# Quantitative analysis of gas chromatography-coupled electroantennographic detection (GC-EAD) of plant volatiles by insects

**DOI:** 10.1101/2024.12.01.626223

**Authors:** Kelsey J.R.P. Byers, Robert N. Jacobs

**Affiliations:** Department of Cell and Developmental Biology, John Innes Centre, Norwich Research Park, Norwich, NR4 7UH, United Kingdom; 6 Greenways, Norwich, NR4 6HE, United Kingdom

**Keywords:** electroantennography, electrophysiology, first forward difference, floral scent, GC-EAD, herbivory, plant volatiles, pollination

## Abstract

**Premise:** Plant-interacting insects receive plant volatile signals through their antennae, and voltage changes across an antenna exposed to volatile stimuli can be measured to determine if insects can perceive them, often by coupling gas chromatography to electroantennographic detection (GC-EAD). Current methods for analysing GC-EAD data rely on *qualitative* observation, rather than using *quantitative* analysis, and are thus prone to bias.

**Methods and Results:** We developed a novel *quantitative* methodology for analysis of GC-EAD data using a signal processing technique on EAD data to compare responses of hawkmoths and bumble bees to a library of common floral volatiles. Responses varied between species and sexes and these responses were in some cases affected by compound type or modification. Our method also works with existing, published GC-EAD datasets.

**Conclusions:** Our novel GC-EAD analysis technique is robust to baseline drift and low signal:noise ratios commonly found in GC-EAD data, and provides a way forward for quantitative studies of plant volatile-mediated plant-insect interactions.

## INTRODUCTION

Plants produce a wide array of secondary metabolites – i.e. those that are not immediately tied to growth and reproduction – with over 200,000 isolated and described across the plant kingdom (Kessler and Kalske, 2018). Many of these compounds play direct roles in the interactions of plants with the abiotic and biotic environment, and have been exploited for this purpose by humanity with uses in industries such as agriculture (e.g. the terpenoid volatile geraniol, which repels eggplant-infesting pest moths (Ghosh et al., 2023; Byers, 2023)) and flavour and fragrance (e.g. vanillin, a volatile product of the phenylpropanoid pathway, which is the dominant compound in natural vanilla extract (Gallage et al., 2014)). Although these compounds can be exploited in such ways, and although chemical structures of numerous plant metabolites have been described, their biological functions remain unknown in many cases (Dixon and Dickinson, 2024). Many of these compounds are volatiles, which are especially well known for influencing plants’ interactions with insects by attracting both beneficial and antagonistic species (e.g. pollinators and herbivores (Raguso, 2008)), with significant implications for plant ecology and evolution (Byers, 2021) as well as agriculture (Pickett and Khan, 2016).

Identifying which volatile cues emitted by plants are perceived by plant-interacting insects is key to our understanding of these cross-kingdom interactions. In particular, identifying insect-relevant volatiles can facilitate work in plant secondary metabolism, ecology, and evolution. For example, identifying pest-resistant crop varieties by screening for known repellent volatiles is cheaper and easier than performing field trials (e.g. Ghosh et al., 2023), while determining which volatiles are key to the ecology and evolution of plant-pollinator interactions can facilitate genetic mapping of only those volatiles rather than the entire plant headspace (e.g. Byers et al., 2014a, 2014b). In short, identifying insect-relevant volatiles – often only a fraction of a plant’s emitted volatiles and individually not always the most abundant volatiles (Clavijo McCormick et al., 2014) – can dramatically reduce the experimental scope and increase the tractability of plant secondary metabolite study.

Electroantennography – the detection and measurement of electrical signals from whole insect antennae – is often used to screen plant volatiles for insect perceptibility. In this technique, a pair of electrodes are mounted at either end of an excised insect antenna or at the tip of the antenna and another reference point such as the base of the head (typically for smaller insects). The continuous voltage difference across the antenna is then amplified and sent to a computer. The insect antenna can be stimulated using single air puffs passed across filter paper spotted with single compounds or a mix of compounds (electroantennography, abbreviated EAG), or instead can be exposed to the effluent from a gas chromatography (GC) column (GC-coupled electroantennographic detection, or GC-EAD). The two signals – from the GC’s main detector (usually a flame ionization detector, or FID) and from the insect antenna – are then aligned and compared by eye to determine whether a response has occurred to a given GC-FID peak. The compounds to which the insect antenna “responds” with some voltage change are taken to be physiologically active and may be further tested via behavioural assays with single pure or mixed compounds or with plant lines known to be high or low in the relevant compounds. While GC-EAD has significant advantages over EAG assays due to its ability to test responses to commercially or synthetically unavailable or unknown compounds present in a chemical mixture, the resulting signal is typically fairly noisy and can be difficult to interpret.

Although these techniques have been in use for nearly seventy years, being first described in 1957 (Schneider, 1957), the resulting data are always presented qualitatively – “response/no response” (e.g. Peakall et al., 2010) – with no quantitative, repeatable measure used to define antennal “response” versus background activity. Indeed, a comprehensive and well-cited overview of GC-EAD includes no mention of how data analysis is performed or what criteria are used to define an “active” GC-EAD peak (Schiestl and Marion-Poll, 2002). This has hampered the field by limiting reproducibility and cross-study or cross-observer comparisons, introducing error, and reducing ease of application to broader plant volatile studies. This may be a particular issue with many plant volatiles, which – unlike insect pheromones – may not have the same degree of “dedicated hardware” on the insect antenna (Akers and Getz, 1993; Schiestl and Marion-Poll, 2002) and thus may produce weaker signals, with the exception of sexually deceptive insect pollination systems which use insect pheromone compounds to attract pollinators.

In this Protocol Note, we present “best practice” statistical approaches incorporating techniques from signal processing to analyse gas chromatography-coupled electroantennographic detection (GC-EAD) data in a quantitative fashion. Signal processing, an engineering technique where mathematical transformations are used to enhance signals and remove background noise, has been used in medical electrophysiology (Venkatachalam et al., 2011; van Drongelen, 2018). To our knowledge it has never been applied to electroantennographic data specifically, despite this being a type of electrophysiological study. Our method uses signal processing analyses for correcting for baseline drift and electrical noise (specifically, the use of the first forward difference and median filtering), background antennal noise (specifically, comparison with response to GC outputs between GC-FID peaks), and antennal sensitivity loss (via use of windowing) are presented. Median filtering is a technique used to reduce impulsive noise and smooth data; it functions by setting the value at a given timepoint to be the median of a surrounding “window” of data of a given length (Lim, 1990), in our case with the goal of removing random electrical noise often present in electroantennography data. The first forward difference (FFD), another signal processing technique, outputs the step change from the value at time *t* to the value at time *t*+*n*, where *n* is 1 for the “first” forward difference used here (Oppenheim and Schafer, 1975). The FFD is the simplest possible way to remove low frequency components, including the baseline drift present in electroantennography data. This baseline drift, often seen as a slow, low frequency wiggle of the antennal response, can superficially resemble legitimate responses and thus needs to be controlled for (Schiestl and Marion-Poll, 2002). As the antenna can lose responsiveness over time and the signal often becomes noisier as it degrades (Byers et al., 2020), our method “windows” the EAD data into discrete time chunks and calculates outlier values for each time chunk separately, rather than calculating the outliers for the entire data set at once. By doing so, we avoid assuming that a significant response in e.g. the 12-16 minute time window (when the antenna may have degraded somewhat) must look identical to a significant response in the earlier e.g. 0-4 minute time window when the antenna is perhaps more sensitive.

A sample data set from six *Manduca sexta* hawkmoths (three males and three females), four *Bombus terrestris audax* bumble bees (all workers), and 28 common volatile stimuli of different volatile chemical compound classes (all present in a variety of plant species) are used to demonstrate the methods and their application in detail and show that they work for a variety of volatile plant secondary metabolites and two major model pollinating insect species. We show that quantitative analyses of GC-coupled electroantennogram data are indeed possible and quite feasible, pointing the way forward for robust, statistically-based analyses of antennal detection ability in volatile-driven plant-insect interactions.

## METHODS AND RESULTS

Please see Appendix 1 for a detailed equipment list and protocol. Perl and R scripts may be found at https://github.com/plantpollinator/GCEADAnalysis/

### Methods: Gas chromatography-coupled electroantennographic detection (GC-EAD) setup and recording

#### Insect sourcing

Adult *Manduca sexta* (L.) (Lepidoptera: Sphingidae) hawkmoth males and females were sourced from a maintained colony reared on artificial diet (Frontiers Scientific Services, Newark, Delaware, USA) at the John Innes Centre Entomology platform facility; adults were not fed but females were allowed to oviposit freely on live *Nicotiana benthamiana* plants and males and females were kept together prior to collection. Worker *Bombus terrestris audax* (Harris) (Hymenoptera: Apidae) were sourced from a single Bumblebee New Season Hive (Biobest, purchased from Agralan Ltd, Wiltshire, UK) which were allowed to freely forage in a custom-built acrylic cage with 25% sucrose and ground pollen *ad libitum* (Sevenhills Wholefoods Organic Raw Bee Pollen, Dinnington, UK). Both species of insect were maintained prior to sampling in growth chambers at 25°C and constant day length (16 hours light, 8 hours dark).

#### EAD test mixture preparation and analysis

To assess antennal responses to multiple compounds delivered at approximately equal flame ionization detector (FID) sensitivities, a mixture of 28 authentic reference standards was assembled, representing 29 GC-FID peaks (farnesol is sold as a mixture of isomers and both elute separately on the GC-FID).

Standards were obtained from a variety of established sources (see Table S1 for vendor addresses and Table S2 for vendor and product numbers) and diluted to 10μg/μl in hexane (Table S2, HPLC-grade, ca. 95% purity). The final compound list was selected to include a variety of chemical compound classes (terpenoids, aromatics, fatty acid derived compounds, and one nitrogenous compound) that would separate well on a 10°C per minute GC oven ramp program. Due to differences in FID responsiveness to the individual compounds (i.e. some compounds may produce different peak areas on the FID despite being at equal injected amounts), different volumes of the 10μg/μl dilutions were added to the experimental mixture (see Table S2 for final concentrations, which ranged from 93ng/μl for (R)-(+)- limonene to 2004ng/μl (2.004μg/μl) for indole).

To determine retention times and confirm mixture GC-FID peak identities, 3ul of the test mixture was manually injected into an Agilent 7890BGC (Agilent, Stockport, UK) fitted with an Agilent 5977A/B MS detector (GC-MS) and a Phenomenex Zebron ZB-5HT column (35m x 0.25mm x 0.10μm; Phenomenex UK Ltd, Macclesfield, UK). The GC run parameters were as follows: the column flow was 1.0603 ml/min of helium carrier gas with an oven program of 50°C for 4 minutes followed by a temperature ramp of 10°C/minute to 230°C, which was then held for 4 minutes. The inlet was splitless and held at 250°C and the MS source was held at 230°C. Compounds were identified using Agilent Unknowns software (v10.1, Agilent, Stockport, UK) and via comparison to Kovats Retention Indices of the individual compounds run separately using the same GC-MS system, calculated using an alkane ladder (49452-U, Sigma-Aldrich, Watford, UK). Two additional “unknown” GC-FID peaks were identified in the final mixture which did not represent any of the chemical standards added or the solvent. As many of the standards used are derived from natural sources rather than synthetic ones and are not completely pure as a result, we expect that these represent additional minor natural products from these sources. Due to our inability to remove these unknown compounds from the mixture, we treated them as a separate chemical compound class and excluded them from downstream analyses that targeted chemical compound classes or modification types.

#### EAD antennal preparation and assembly

*Manduca sexta* male and female adults: single antennae were excised using microdissection scissors from three individuals each of living adult male and female *Manduca sexta*. Due to biosecurity requirements for this non-native potential plant pest species, excised antennae were placed in a 15ml conical tube on wet tissue paper to maintain antennal humidity levels and carried to the electrophysiology setup in a separate building. The fine tip of an antenna (representing ca. 7 segments, cut where the tip abruptly broadens) was removed using a microscalpel under a dissection microscope. Tip removal is commonly done (though is not mandatory) in electroantennographic studies, as it may reduce electrical resistance and thus improve signal (Schoonhoven 1973). The excised and trimmed antenna was then mounted on an antenna electrode fork (Syntech PRG-2, Ockenfels Syntech, Buchenbach, Germany) using electrode gel (Signa Gel, Parker Laboratories, Fairfield, New Jersey, USA), which was then plugged in to the electrode mount (Syntech MP-15 with EAG Combi Probe) and moved into position in front of the GC eluent-carrying air stream approximately 1cm from the air stream outlet. Approximately ten minutes elapsed between antennal excision and mounting on the antenna electrode fork due to the walk between buildings.

*Bombus terrestris audax* workers: individuals were removed from the Entomology platform facility in 50ml conical tubes, then kept at 4°C for 30 minutes to enable safe antennal removal. A single antenna was then removed using microdissection scissors under red light (with the insect held using a set of forceps) and the tip of the final flagellar segment removed using a microscalpel under a dissection microscope, then treated as above.

#### GC-EAD setup and recording procedure

The GC-FID setup consisted of an Agilent 8890 GC fitted with a flame ionization detector (FID). The column (a Phenomenex Zebron ZB-5HT column (35m x 0.25mm x 0.10μm), identical to the one used for GC-MS analysis) was fitted to a splitless inlet kept at 250°C. The detector end of the column was fitted to the bottom end of an inert glass Y-splitter (Agilent UltraInert Universal PressFit Y-Splitter, part 5190- 6980). The two other arms of the splitter were fitted with deactivated column segments of equal 30cm length (Agilent CP803210, 0.32mm inner diameter), with one end running to the FID (which was kept at 300°C) and the other end leaving the GC oven through a port on the side which was fitted with a Syntech EC-03 heated transfer tube kept at 250°C. The GC used hydrogen as a carrier gas, which was generated with a Parker Hydrogen Generator (H2PEM-100, Thames Restek UK Ltd, Saunderton, United Kingdom). Nitrogen gas at a flow of 25ml/min was used as a make-up gas, and the FID was also fed with compressed air at 400ml/min to ensure adequate flame gas supply. The GC run parameters were as follows: the column flow was 4.7 ml/min with an oven program of 50°C for 4 minutes followed by a temperature ramp of 10°C/minute to 230°C, which was then held for 4 minutes. Each manual sample injection consisted of 3ul of the EAD test mixture, for a total of 0.279-6.012μg of each compound injected. GC-FID chromatogram waveforms were exported from Agilent OpenLab CDS v2.5 software as CSV files consisting of an FID amplitude value (in pA) for each given sampled timepoint.

The EAD recording setup consisted of a Syntech IDAC-2 connected to the desktop computer controlling the GC-FID system. A constant air flow (setting 8 on the CS-55) was delivered using a Syntech CS-55 Stimulus Controller throughout the experiment. Prior to and shortly after each GC-FID run, a puff of the EAD test mixture (via a Pasteur pipette fitted with filter paper, with 10μl of the EAD test mixture added onto the paper, also setting 8 on the CS-55) was manually delivered using the CS-55 to ensure antennal viability had been maintained throughout the GC-FID run. EAD signals were captured using Syntech GcEAD 2014 v1.2.5 software with default settings and exported as CSV waveforms in GcEAD 2014 (measured as millivolts) prior to analysis.

### Methods: Data processing and analysis

#### GC-FID peak identification

A custom Perl script (Perl, 2023; see Appendix for script) was developed to determine the “onset” and “offset” timings (i.e. when a peak began and when it ended) of GC-FID peaks above a certain user-defined threshold from the FID CSV data. In the presented GC-FID experimental data, the threshold was set at 50pA to capture all significant GC-FID peaks while avoiding inclusion of small GC-FID peaks likely representing contamination from the hexane solvent. Note that this resulted in the merger of the closely eluting caryophyllene oxide and ethyl laurate GC-FID peaks, which were manually removed from the downstream analyses of chemical compound class and compound modification impact on GC-EAD responses as a result.

#### Data loading and initial processing

EAD and FID waveforms were loaded as CSV files into R (R Core Team, 2024; see Appendix for example script). GC-FID onsets and offsets determined in the previous section were also loaded. As the EAD setup samples at a rate of 6000 datapoints per second and does not export a timestamp to its CSV file, a new time variable was defined corresponding to the sampling rate. The GC-FID injection time and a 20 minute endpoint (after all mixture GC-FID peaks had eluted) relative to the EAD time were also defined, as were EAD timestamps corresponding to each GC-FID peak onset and offset. As injections were done manually for the GC-FID, there was occasionally a slight disparity between GC-FID peak onsets and the onset of a significant-seeming EAD pattern, and so a user-defined offset was determined by comparing the most significant EAD pattern onset with the previously determined corresponding GC-FID peak onset.

#### Median filtering, First Forward Difference (FFD) processing, and significant FFD value identification

To remove random electrical noise from the EAD signal, we first applied a median filter using the runmed() function in base R (specifically as `runmed(EADamplitude,17)`). A window of 17 samples was chosen based on the width of the random electrical spikes seen in the *Acleris* data from Dr. Glenn Svensson (**Figure 5B**). In order to remove the effects of the “wandering baseline” common to GC-EAD, we used the `diff()` function in base R (specifically as `diff(EADamplitude,lag=1,differences=1)`) to calculate the First Forward Difference of this median filtered antennal amplitude. The FFD vector was subsetted into five windows of four minutes each for a total GC-FID run of 20 minutes for the hawkmoth and bumble bee data. For the data from Dr. Svensson, we instead used three minutes each for a total GC-FID run of 21 and 14.5 minutes for the *Acleris* and *Plodia* data respectively. For each window, the average and standard deviation of the FFD within that window was calculated. A significance threshold of 3σ above or below the average was calculated, and data points that exceeded the significance threshold for their respective windows were marked as significant and termed “significant spikes”.

#### Calculating the number of significant FFD spikes per second for each identified GC-FID peak

We first calculated how many GC-EAD FFD spikes above the 3σ threshold were within each GC-FID peak. For each GC-FID peak identified, the number of significant GC-EAD FFD spikes as defined above within its onset-offset window was determined using the ‘between’ function in the R package `dplyr` (Wickham et al., 2023), which identified significant spikes that were within a GC-FID peak window. This value was divided by the length of the GC-FID peak in seconds (offset time minus onset time) to get the significant FFD spike frequency of spikes/second. As EAD data can have significant background noise (Schiestl and Marion-Poll, 2002), we defined the significant FFD spike frequency of the space between GC-FID peaks as the threshold for whether a GC-FID peak attracted significant EAD activity for each window: if the significant FFD spike frequency for a given GC-FID peak exceeded the background significant FFD spike frequency of its time window, the GC-FID peak’s EAD response was counted as ”EAD active”, and otherwise considered non-active.

#### Statistical analysis

To compare results between individuals within a sex/caste and between species, a “significant response” across each species-sex/caste category (female *M. sexta*, male *M. sexta*, and worker *B. terrestris*) was identified for chemical compounds where >2/3 of individuals tested were “EAD active” for that peak. To investigate whether the results were repeatable between moths, we calculated the Pearson correlation coefficient between the significant spike frequency for all pairwise combinations of moths within a sex using cor.test() in R. We then used the linear model function `lm()` in R to calculate the slope and intercept of the relationship between each pair of moths within a sex. Finally, we used `lm()` and the estimated marginal means function `emmeans()` (Lenth, 2024) to determine whether chemical compound class (e.g. terpenoid) or modification type (e.g. alcohol) had an influence on significant spike frequency within each species-sex/caste group (female *M. sexta*, male *M. sexta*, and worker *B. terrestris*).

#### Further test with unpublished and published data

Unpublished and published GC-EAD data were kindly donated by Dr. Glenn Svensson (Pheromone Group, Lund University). Data from *Plodia interpunctella* (Lepidoptera: Pyralidae) were of a single male in response to a synthetic mixture of four female pheromone components. Data from *Acleris comariana* (Lepidoptera: Tortricidae) were of a single male in response to a natural female pheromone gland extract. As we had difficulty employing our GC-FID automated peak detector on these data due to significant noise and a long solvent peak trailing end in the GC-FID signal, we instead manually annotated GC-FID peaks in GcEad 2014 software and used those retention times in the downstream analysis. Due to the different length of the GC-EAD runs, we chose 3 minute windows for calculation of significant FFD spikes instead of 4 minute windows as in our earlier experiments.

## Results

### The new method allows quantitative analysis and avoids baseline drift

Here we present a method that avoids the otherwise significant biases inherent in GC-EAD *qualitative* analysis methodology by the use of signal processing techniques, specifically *median filtering* and the *first forward difference* (FFD), to remove the influence of electrical noise and baseline drift respectively, essentially flattening out the EAD signal (**Figure 1B**) and replacing its waveform with a series of individual spikes (**Figure 1C**, see also **Figures S16-S30**). A significance threshold may then be set for each window (horizontal lines in **Figure 1C** and **Figures S16-S30**, both showing ± 3 standard deviations above/below the mean for that window), and the frequency of spikes with an amplitude over that significance threshold during a given GC-FID peak may be easily calculated and analysed using traditional statistical methods. This results in a *quantitative* rather than *qualitative* assessment of whether a GC-FID peak is physiologically active, i.e. provokes a significant GC-EAD response, or not.

**Figure 1:**
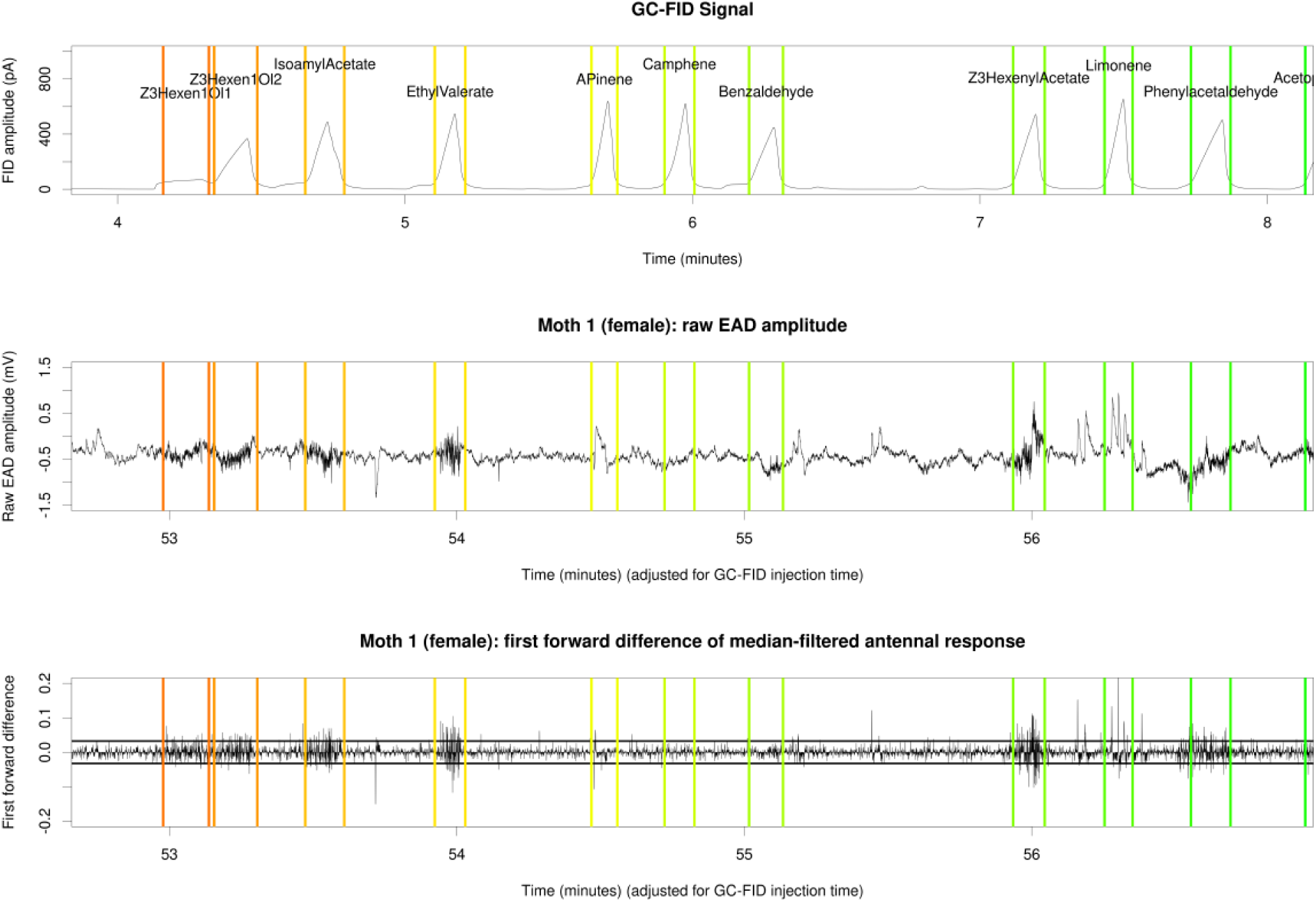
Comparison of raw EAD and first forward difference-processed EAD traces from a single female *Manduca sexta* hawkmoth antenna between 4-8 minutes of GC elapsed time.(A) GC-FID trace annotated with automatically detected GC-FID peak starts and ends (colored vertical lines) and GC-FID peak identities. (B) Raw EAD amplitude annotated with GC-FID peak starts and ends. (C) First forward difference-processed EAD amplitude annotated with GC-FID peak starts and ends. Note the “wandering baseline” in (B) that is absent in (C).

### GC-FID peaks with significant EAD responses

**Table 1** shows the list of compounds tested in the EAD test mixture along with their significant spike rates for female and male moths and worker bumble bees and how many individuals of each had a significant spike rate over the background spike rate. In general, female moths were most responsive to fatty acid-derived compounds (FADs, red in **Figure 2A**, see also **Figure S43**) and to some but not all aromatic volatiles (blue in **Figure 2A**). Responses to terpenoids were more mixed (green in **Figure 2A**), with some compounds (e.g. (geraniol) eliciting significant spike frequencies, while others (e.g. camphene) eliciting no significant spike requencies in any female. The sole tested nitrogenous compound, indole (yellow in **Figure 2A**), also elicited significant spike frequencies in all three female moths. Overall, female moths responded significantly to 20 of the 31 tested compounds (excluding the hexane GC-FID peaks). Male moths were generally less responsive than females to the majority of compounds, with overall significant spike frequencies to 20 of the 31 tested compounds (excluding the hexane GC-FID peaks) but with no significant difference in significant spike frequency between chemical compound classes (e.g. terpenoids versus aromatics) (**Figure S43A,B**); 2 of 3 males were responsive to indole. Worker bumble bees responded significantly to 8 of the 30 compounds (“unknown 2” was missing from this analysis as it was not picked up by the GC-FID peak detector script). Bees responded equally (i.e. had equal significant spike frequencies) to all types of chemical compound classes (**Figure S43C**), although only one of four tested bees was significantly responsive to indole, the sole nitrogenous compound tested.

**Figure 2:**
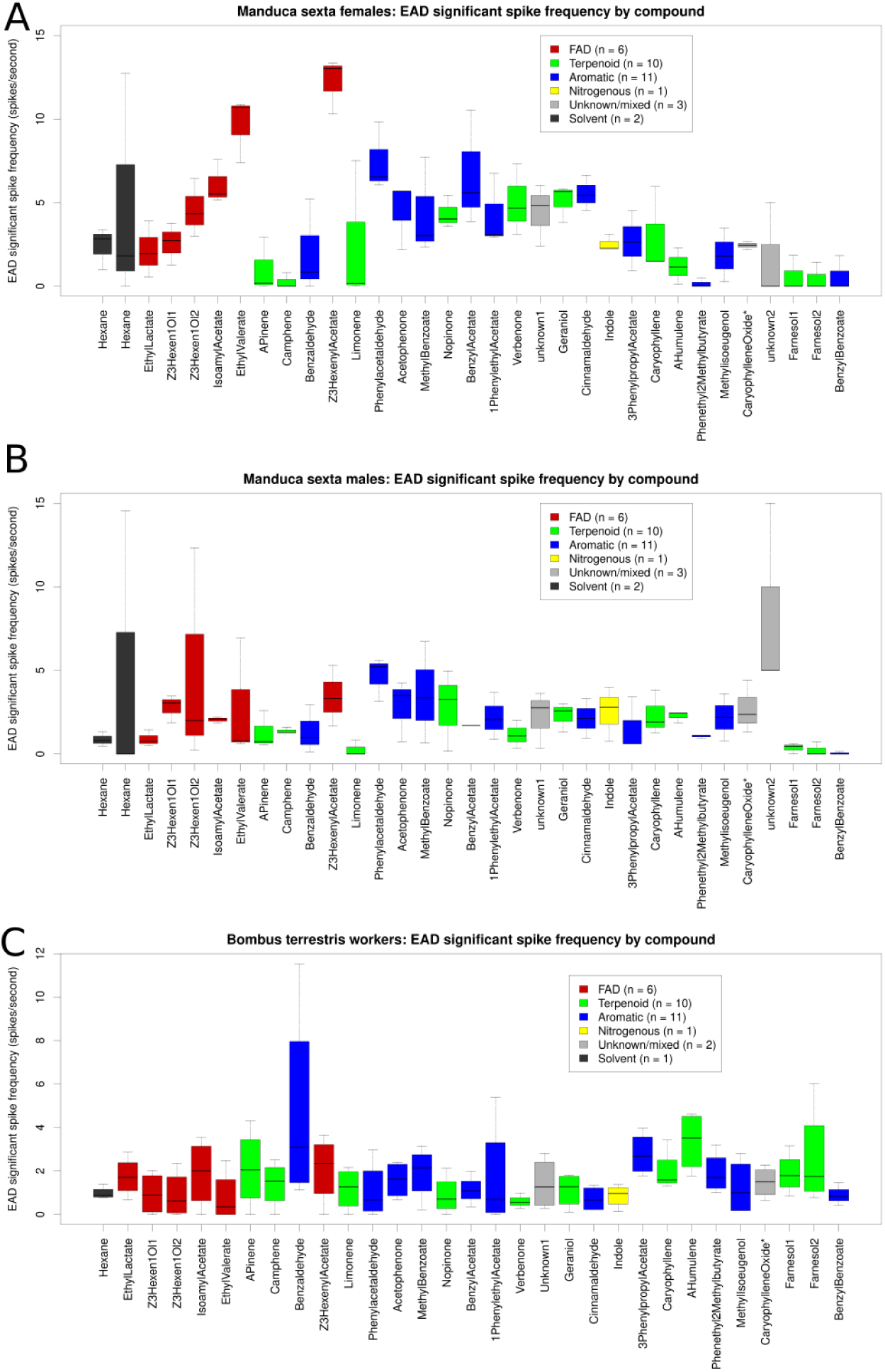
First forward difference-processed EAD responses to 33 compounds from the test mixture, colored by compound type and sorted by retention time (note that the Y-axis scaling differs for (C)). Note for caryophyllene oxide*: this box includes responses to ethyl laurate, as the GC-FID peaks were merged in the GC-FID annotation thresholding step so are treated as a single GC-FID peak by the analysis. (A) Data from three *Manduca sexta* females. (B) Data from three *Manduca sexta* males. (C) Data from four *Bombus terrestris audax* workers.

**Table 1:** Compounds used in the EAD test mixture with details of their retention time, chemical compound class and subclass or modification categorization, and average significant spike frequency (spikes per second) for female and male *Manduca sexta* hawkmoths and worker *Bombus terrestris audax* bumble bees. The ratio after the significant spike frequency indicates the number of samples where the given compound elicited a greater response than the background level of spikes versus the total number of samples with that peak present. Bolded responses: >2/3 of antennal samples reacted to that peak for a given species-sex/caste combination. Unknown superscripts: 1^A^: an unclear oxygenated cyclic molecule from GC-MS results; 2^B^: likely an unknown sesquiterpenoid oxidefrom GC-MS analysis results.

| Retent<br>ion<br>time<br>(GC-FID,<br>minute<br>s) | Compound<br>identity | Chemical<br>compound<br>class | Modifications<br>/ subtype | Average<br>*Manduca sexta*<br>female<br>response<br>(significant hit<br>frequency,<br>spikes/second) | Average<br>*Manduca*<br>*sexta* male<br>response<br>(significant<br>hit<br>frequency,<br>spikes/secon<br>d) | Average<br>*Bombus*<br>*terrestris*<br>*audax* worker<br>response<br>(significant hit<br>frequency,<br>spikes/second) |
| --- | --- | --- | --- | --- | --- | --- |
| 1.973 | Hexane, main<br>GC-FID peak | Solvent | NA | 2.408 (NA) | 0.873 (NA) | 0.975 (NA) |
| 2.687 | Hexane, small<br>GC-FID peak | Solvent | NA | 4.848 (NA) | 4.848 (NA) | NA (NA) |
| 3.755 | Ethyl L-lactate | FAD | Ester | **2.133 (2/3)** | **0.900 (2/3)** | **1.738 (3/4)** |
| 4.291 | (Z)-3-hexen-1-ol,<br>shoulder GC-FID<br>peak | FAD | Alcohol | **2.583 (3/3)** | **2.792 (3/3)** | 0.950 (2/4) |
| 4.458 | (Z)-3-hexen-1-ol,<br>main GC-FID<br>peak | FAD | Alcohol | **4.593 (3/3)** | **4.852 (2/3)** | 0.897 (2/4) |
| 4.741 | Isoamyl acetate | FAD | Ester | **6.094 (3/3)** | **2.045 (3/3)** | 1.886 (2/4) |
| 5.184 | Ethyl valerate | FAD | Ester | **9.659 (3/3)** | 2.782 (1/3) | 0.796 (1/4) |
| 5.716 | (1R)-(+)-α-pinene | Terpenoid | Monoterpene | 1.049 (1/3) | 1.296 (1/3) | **2.097 (3/4)** |
| 5.985 | Camphene | Terpenoid | Monoterpene | 0.267 (0/3) | **1.387 (3/3)** | 1.396 (2/4) |
| 6.292 | Benzaldehyde | Aromatic | Aldehyde | **2.019 (2/3)** | 1.362 (1/3) | **4.715 (4/4)** |
| 7.200 | (Z)-3-hexenyl<br>acetate | FAD | Ester | **12.222 (3/3)** | **3.434 (3/3)** | **2.087 (3/4)** |
| 7.505 | (R)-(+)-limonene | Terpenoid | Monoterpene | 2.564 (1/3) | 0.285 (0/3) | 1.176 (2/4) |
| 7.847 | Phenylacetaldeh<br>yde | Aromatic | Aldehyde | **7.475 (3/3)** | **4.646 (3/3)** | 1.074 (2/4) |
| 8.222 | Acetophenone | Aromatic | Ketone | **4.526 (3/3)** | **2.822 (2/3)** | 1.581 (2/4) |
| 8.691 | Methyl benzoate | Aromatic | Ester | **4.370 (3/3)** | **3.585 (2/3)** | **1.912 (3/4)** |
| 9.355 | (1R)-(+)-nopinone | Terpenoid | Monoterpenoid ketone | **4.341 (3/3)** | **2.791 (2/3)** | 0.885 (2/4) |
| 9.818 | Benzyl acetate | Aromatic | Ester | **6.667 (3/3)** | **1.705 (3/3)** | 1.126 (2/4) |
| 10.279 | 1-phenylethyl acetate | Aromatic | Ester | **4.265 (3/3)** | **2.206 (3/3)** | 1.696 (2/4) |
| 10.511 | (1S)-(-)-verbenone | Terpenoid | Monoterpenoid ketone | **5.030 (3/3)** | 1.158 (1/3) | 0.594 (0/4) |
| 10.609 | Unknown 1<sup>A</sup> | Unclear | Unclear | **4.425 (3/3)** | **2.241 (2/3)** | 1.408 (2/4) |
| 11.240 | Geraniol | Terpenoid | Monoterpenoid alcohol | **5.111 (3/3)** | **2.306 (2/3)** | 1.117 (2/4) |
| 11.459 | (E)-cinnamaldehyde | Aromatic | Aldehyde | **5.532 (3/3)** | **2.128 (3/3)** | 0.718 (2/4) |
| 11.777 | Indole | Nitrogenous | Aromatic | **2.545 (3/3)** | **2.509 (2/3)** | 0.856 (1/4) |
| 12.691 | 3-phenylpropyl acetate | Aromatic | Ester | **2.687 (2/3)** | 1.550 (1/3) | **2.772 (4/4)** |
| 13.254 | β-caryophyllene | Terpenoid | Sesquiterpene | 2.979 (1/3) | **2.340 (2/3)** | 1.974 (2/4) |
| 13.683 | α-humulene | Terpenoid | Sesquiterpene | 1.199 (1/3) | **2.254 (2/3)** | **3.352 (4/4)** |
| 14.111 | Phenethyl 2-methylbutyrate | Aromatic | Ester | 0.159 (0/3) | 1.058 (0/3) | **1.900 (3/4)** |
| 14.285 | Methyl isoeugenol | Aromatic | Alcohol | 1.849 (1/3) | 2.194 (1/3) | 1.240 (2/4) |
| 15.265 | Caryophyllene oxide + ethyl laurate (GC-FID peaks merged by GC-FID peak identification script) | Terpenoid + FAD | Sesquiterpenoid oxide + ester | **2.449 (3/3)** | **2.694 (2/3)** | 1.477 (2/4) |
| 15.476 | Unknown 2<sup>B</sup> | Terpenoid | Unknown sesquiterpenoid oxide | 1.667 (1/3) | **8.333 (3/3)** | NA (NA) |
| 16.474 | Farnesol, isomer 1 | Terpenoid | Sesquiterpenoid alcohol | 0.615 (0/3) | 0.359 (1/3) | 1.895 (2/4) |
| 16.734 | Farnesol, isomer 2 | Terpenoid | Sesquiterpenoid alcohol | 0.476 (0/3) | 0.238 (1/3) | 2.573 (2/4) |
| 17.189 | Benzyl benzoate | Aromatic | Ester | 0.611 (0/3) | 0.051 (0/3) | 0.885 (0/4) |

Compounds containing oxygenated modifications, particularly esters, seemed to produce the highest level of significant spike frequency in female moths, but not in males (**Figure S43D,E**). At least two-thirds of females responded significantly to 8 of 10 presented esters (analyzed without the co-eluting GC-FID peaks ethyl laurate and caryophyllene oxide), 3/6 alcohols, 3/3 aldehydes, and 3/3 ketones, but not to any of the five presented unmodified monoterpenes ((1R)-α-pinene, camphene, and R-(+)-limonene) or sesquiterpenes (caryophyllene, α-humulene), both groups of which lack oxygen atoms. Male moths responded to 6 of 10 esters, 3/6 alcohols, 2/3 aldehydes, and 2/3 ketones but, unlike females, responded to 3/5 of the unmodified (mono/sesqui)terpenes. Males also responded significantly overall to the “Unknown 2” compound, whose identity remains unclear despite GC-MS analysis but which likely also is oxygenated based on tentative NISTlibrary matches. For bees, there was no difference in response based on compound modification, oxygenated or otherwise (**Figure S43F**). At least 2/3 of bees responded to 5 of 10 esters, 1/3 aldehydes, and 2/5 unmodified (mono/sesqui)terpenes, but fewer than 2/3 of bees responded to the 6 alcohols or 3 ketones.

### Repeatability within moth sexes and comparisons between sexes

Responses to non-hexane compounds were highly repeatable between female moths (**Figure S44A**, Pearson’s correlation coefficient range 0.537 to 0.762, all significant), but not between male moths (**Figure S44B**, Pearson’s correlation coefficient range 0.025 to 0.476, only one significant). In general, male moths were substantially less responsive to non-hexane chemical stimuli in this experiment, as represented by their lower average significant spike frequency (**Table 1**, range 0.051-8.333 significant spikes/second) versus females (range 0.159-12.222 significant spikes/second). Responses between males and females trended towards correlation (Pearson’s r = 0.342, *P* = 0.051, **Figure S45**).

### Validation of the method with existing external data

To validate our method with existing GC-EAD data from another lab and setup, we obtained both published and unpublished GC-EAD data from Dr. Glenn Svennson (Pheromone Research Group, Lund University). These data represent responses of *Plodia interpunctella* (Lepidoptera: Pyralidae; **Figure 4, Table S3**) and *Acleris comariana* (Lepidoptera: Tortricidae; **Figure 5, Table S4**) to synthetic and natural pheromones respectively. In both cases our method was able to identify significant GC-EAD peaks, including confirming the published *A. comariana* response to the pheromone component (*E*)-11,13- tetradecadienal (Svensson et al., 2019). Of note, we identified multiple significant spike frequency responses in both datasets that were not identified as significant by previous qualitative analyses (**Tables S3, S4,** Svensson et al., 2019); as our method more unambiguously identifies low-amplitude responses, we suggest ranking potential compounds of interest by their significant FFD hit frequency when prioritizing which compounds to take further via behavioural analysis. The most significant spike frequency by far in the *A. comariana* data, for example, was to the published bioactive compound (**Figure 5, Table S4**).

**Figure 3:**
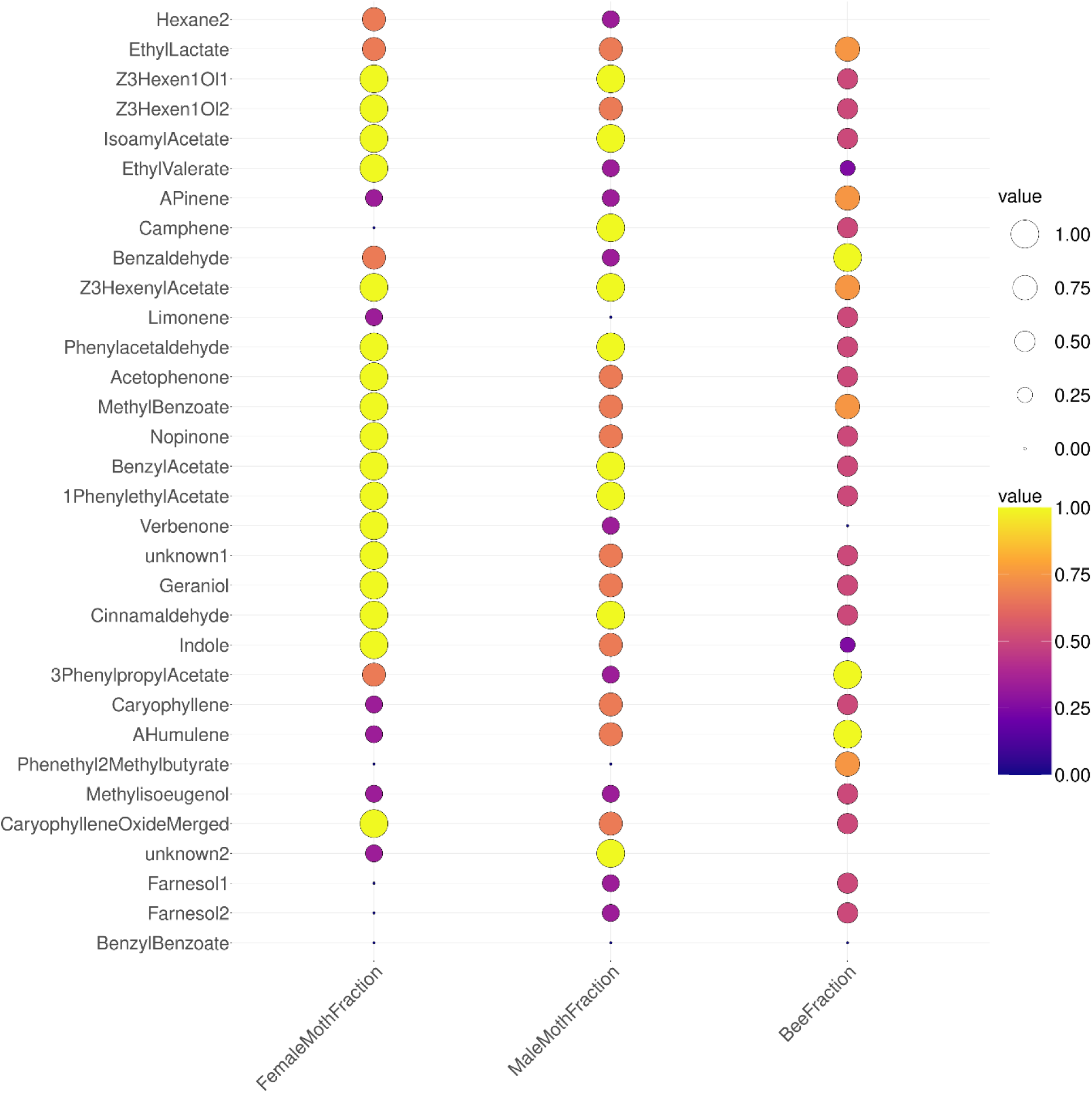
Proportion of individual *Manduca sexta* hawkmoths (“FemaleMothFraction” and “MaleMothFraction”) and *Bombus terrestris audax* worker bumble bees (“BeeFraction”) that had significant responses to each chemical compound tested. Larger circles and warmer colors indicate that more of the individuals were responsive to each compound, while smaller circles and cooler colors indicate that no individuals or only one were responsive. Absent points indicate the specific peak was not detected in the GC-FID run for a given insect.

**Figure 4:**
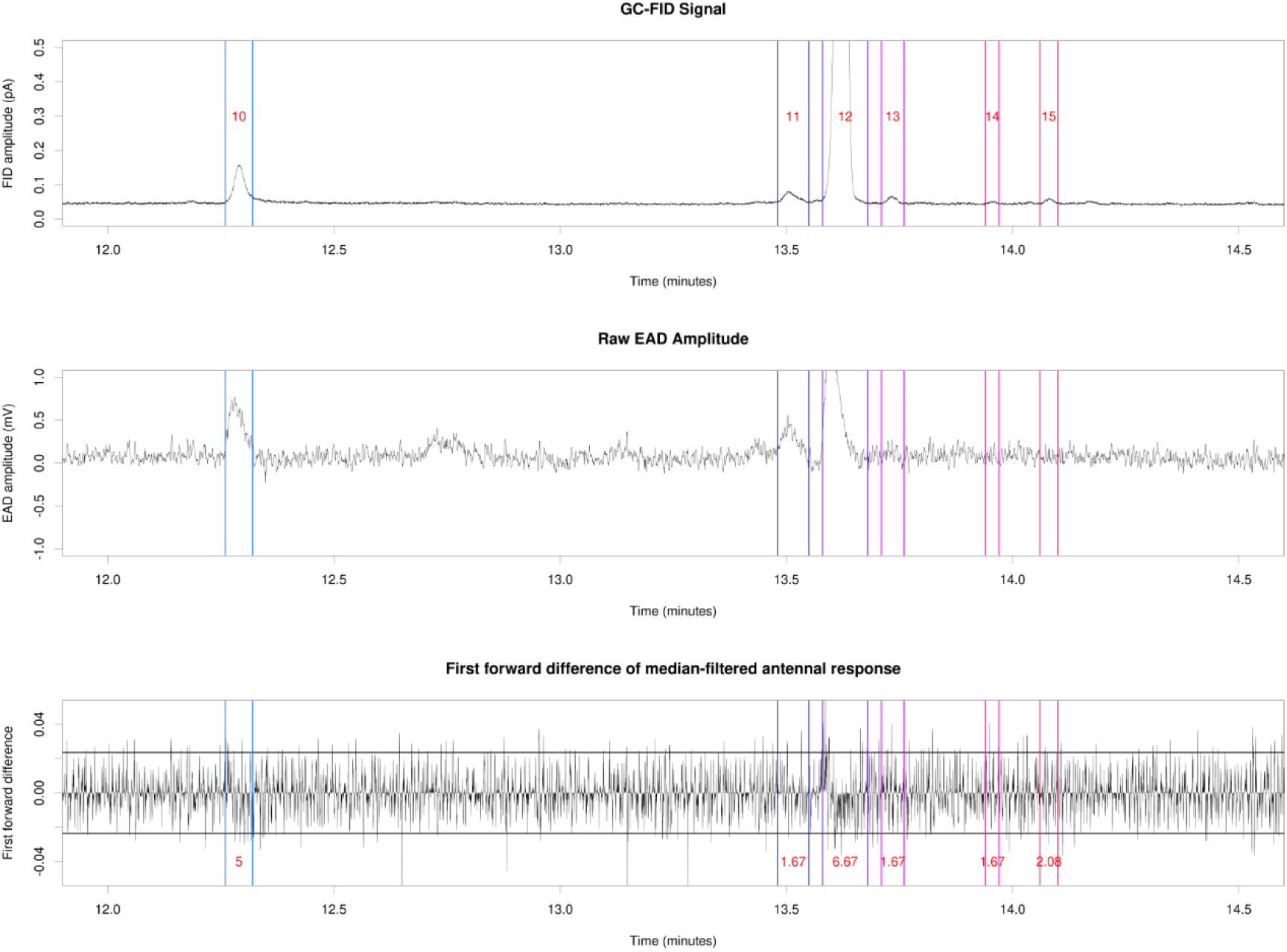
Reanalysis of unpublished GC-EAD data from the Pheromone Group, University of Lund (raw data kindly provided by Dr. Glenn Svensson). Data show the response of a male *Plodia interpunctella* (Lepidoptera: Pyralidae) to a synthetic mixture of four female pheromone compounds. (A) GC-FID trace annotated with manually identified GC-FID peak starts and ends (colored vertical lines) and GC-FID peak numbers. (B) Raw EAD amplitude annotated with GC-FID peak starts and ends. (C) First forward difference-processed median-filtered EAD amplitude annotated with GC-FID peak starts and ends. GC- FID peaks showing significant EAD spike frequency are annotated with that frequency in red text in (C) (all annotated GC-FID peaks within this time window).

**Figure 5:**
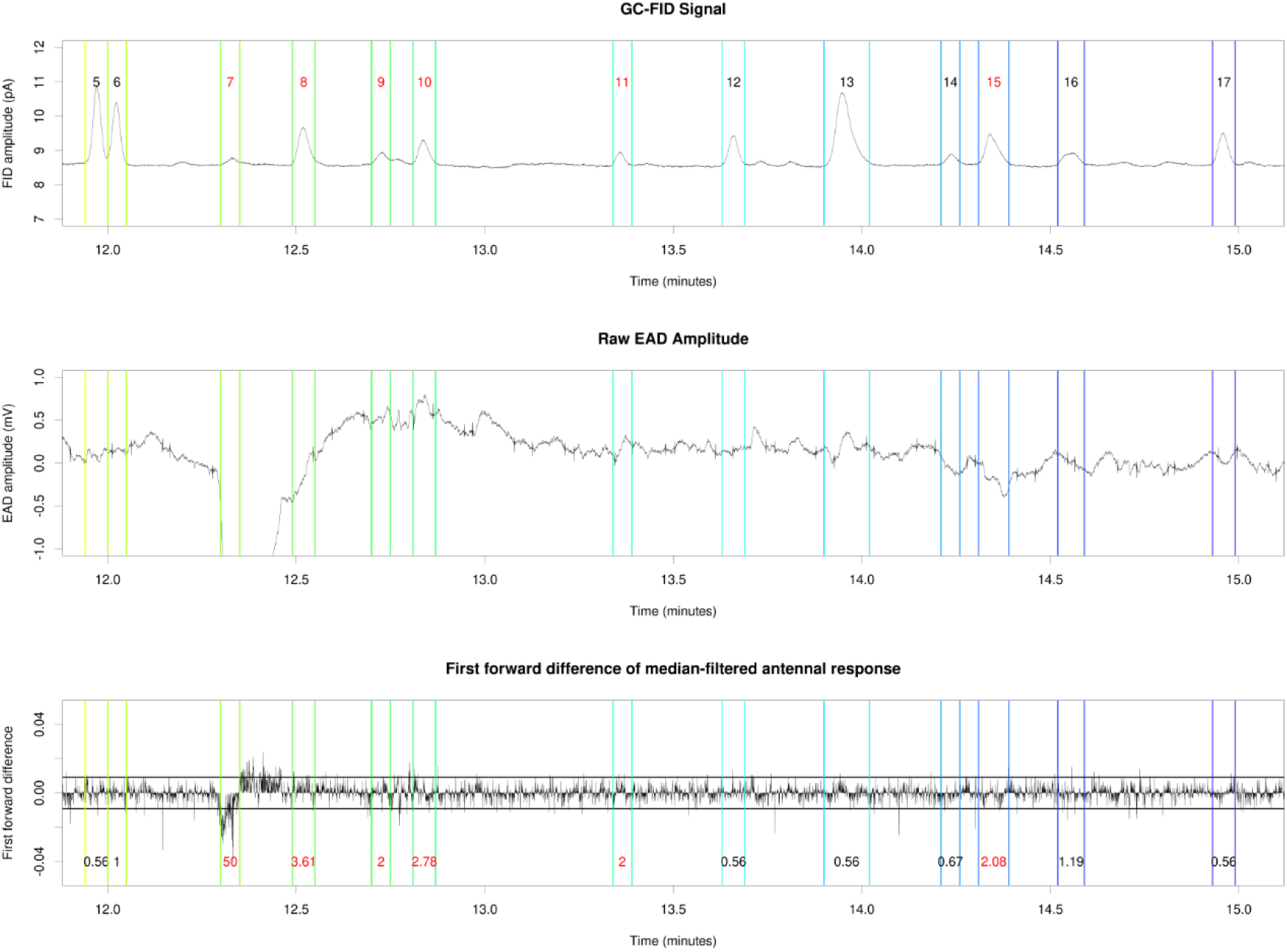
Reanalysis of published GC-EAD data from the Pheromone Group, University of Lund (raw data kindly provided by Dr. Glenn Svensson) (Svensson et al., 2019). Data show the response of a male *Acleris comariana* (Lepidoptera: Tortricidae) to an extract of the female pheromone gland. (A) GC-FID trace annotated with manually identified GC-FID peak starts and ends (colored vertical lines) and GC-FID peak numbers. (B) Raw EAD amplitude annotated with GC-FID peak starts and ends; note the regular small spikes every few seconds, presumably due to electrical noise, which were later removed using a median filter. (C) First forward difference-processed median-filtered EAD amplitude annotated with GC-FID peak starts and ends. GC-FID peaks showing significant EAD spike frequency are annotated with that frequency in red text in (C). GC-FID peak 7 corresponds to (*E*)-11,13-tetradecadienal, the biologically active pheromone component in this species. Note that other GC-FID peaks also provoked a significant response in the male antenna, although they are not listed as EAD-active in (Svensson et al., 2019).

## CONCLUSIONS

In this Protocol Note, we have presented a novel signal processing-based methodology for use with gas chromatography-coupled electroantennographic detection (GC-EAD) measurements of insect antennae in response to plant volatiles. Both female and male *Manduca sexta* hawkmoths were responsive to approximately 2/3 of tested compounds in a highly repeatable fashion, although male hawkmoths of the same species were far less repeatable between individuals. In female moths, both chemical compound class (FAD, aromatic, terpenoid, or nitrogenous) and modification (none, alcohol, aldehyde, ester, ketone, or oxide) played a significant role in response patterns, while in male moths this was true for neither modification type nor chemical compound class. Worker *Bombus terrestris audax* bumble bees showed an intermediate level of response, with significant spike frequency responses to one quarter of tested compounds, and with no difference in response rate relative to chemical compound class or modification.

To ensure that our results were not simply due to some property of *Manduca sexta* antennae, we also analysed the same stimulus mixture with four workers of the polylectic (wide dietary breadth, Rasmont et al., 2008) bumble bee *Bombus terrestris audax* using the same data collection and analysis methodologies. We found three compounds that were significant in >2/3 of tested individuals across all three species-sex/caste combinations (female hawkmoths, male hawkmoths, worker bumble bees): ethyl L-lactate, (*Z*)-3-hexenyl acetate, and methyl benzoate (**Table 1**). Results from all three groups are presented in **Figure 3** and **Table 1**. We also observed overall lower significant spike frequencies in bumble bees compared with hawkmoths, despite bumble bees’ robust responses to single puffs of the EAD stimulus mixture. The signal to noise ratio threshold in the bumble bee antennae appears to be weaker than in the hawkmoths (as shown by much lower first forward difference significance thresholds and a lower range of significant average spike frequencies (range 0.594-4.715 hits/second compared with hawkmoth values of 0.159-12.222 and 0.594-4.715 for females and males respectively). We separately conducted a non-exhaustive literature review to identify whether our tested compounds had previously been identified as electroantennographically active in either *Manduca sexta* or *Bombus terrestris*, with the results shown in **Table S3**. In general, many of our tested compounds had not previously been tested with GC-EAD or puff-based (without GC separation) electroantennography, and those that had been often produced results that differed between studies, further suggesting the need for a quantitative analysis technique such as the one presented here.

Consistent with our findings, some compounds were significant in some studies but not others, emphasizing the importance of testing stimuli on multiple individuals of a given species rather than relying on one or two individuals, given that we found variability even between individuals of the same commercial bumble bee hive and hawkmoth colony. The variability commonly seen in GC-EAD experiments – including ours - can also be due to experimental factors such as humidity, gel coverage of antennal sensillae, insect age and time between antennal excision and mounting. The weaker responses seen in bumble bees further necessitate the use of this new technique to ensure unbiased assessment of electroantennographic detection studies in potentially less strongly responsive insects.

By studying responses to common plant volatiles in two model pollinator species, we have shown that responses can be species- and sex-specific, and that it may be difficult to generalize results from one pollinator group to another, or even between populations or genotypes of the same species. Further study will be needed to determine whether results can be generalized within a genus of pollinators, or between those that are generalists and specialists. However, at least one species of pollinating insect was responsive to the majority of the plant compounds tested here. Interestingly, there was no consistent GC-EAD response in any species tested to the monoterpene (R)-(+)-limonene. Limonene is the most widely distributed floral volatile in terms of angiosperm families in which it is found (Schiestl, 2010), and is known to provoke a responsive in the antennal lobe of *Bombus vosnesenskii* (Byers et al., 2014a).

The data analysis methods introduced make use of existing typical GC-EAD recording setups and experimental methods, differing only in the downstream processing of collected GC-EAD data, and thus may be used with both existing as well as novel datasets. In addition, they are simple to execute, requiring only minimal implementation in R or another statistical processing language (example code is provided in the Appendix). This study provides the first truly quantitative analysis of insect responses to common plant volatiles using GC-EAD, and paves the way forward for future GC-EAD analyses to incorporate quantitative, unbiased data processing methodologies to determine whether plant volatiles are physiologically active to interacting insect species.

## AUTHOR CONTRIBUTIONS

KJRPB conceived the project, designed and executed the data collection and data analysis methodology, and wrote the manuscript. RNJ contributed significant signal processing methodology ideas and expertise to the data analysis methodology development. Both authors approved the final version of the manuscript.

## ACKNOWLEDGMENTS

The authors thank the John Innes Centre (JIC) Entomology platform for maintaining the *Manduca sexta* and *Bombus terrestris* colonies and the JIC Metabolomics platform for GC-MS maintenance. We also thank Dr. Glenn Svensson (Pheromone Research Group, Lund University) for providing published and unpublished GC-EAD data, allowing us to test our method on well-validated results from another GC-EAD equipment setup. Funding was provided by the UK Biotechnology and Biological Sciences Research Council Institute Strategic Programmes (Harnessing Biosynthesis for Sustainable Food and Health (HBio) (grant no. BB/X01097X/1) and Building Robustness in Crops (BRiC) (grant no. BB/X01102X/1)). We also thank anonymous reviewers and the editor for their helpful suggestions, which greatly enhanced the manuscript and method.

## DATA AVAILABILITY

All raw data (GC-FID CSV files, electroantennography CSV files, and relevant GC-FID injection timestamps) are available on FigShare (doi: XXX). Scripts referenced in the main text and appendices are also available on GitHub (https://github.com/plantpollinator/GCEADAnalysis).

## Notes

### Competing Interest Statement

The authors have declared no competing interest.

### Summary of Updates

We have updated the analysis method to include median filtering and a comparison against background significant spike frequency rather than the spike frequency of the solvent peak. We have also added analysis of an unpublished and published GC-EAD dataset to show the method works with data beyond our own. In addition, the text of the manuscript has been updated to include a more thorough description of the methods used as an introduction for the reader.

