## Supporting Information Figures and Tables for "Quantitative analysis of gas chromatography-coupled electroantennographic detection (GC-EAD) of plant volatiles by insects"

**Short title: Byers & Jacobs – Quantitative insect GC-EAD for plant interaction analysis**

### SUPPORTING INFORMATION: SUPPLEMENTAL FIGURES AND TABLES

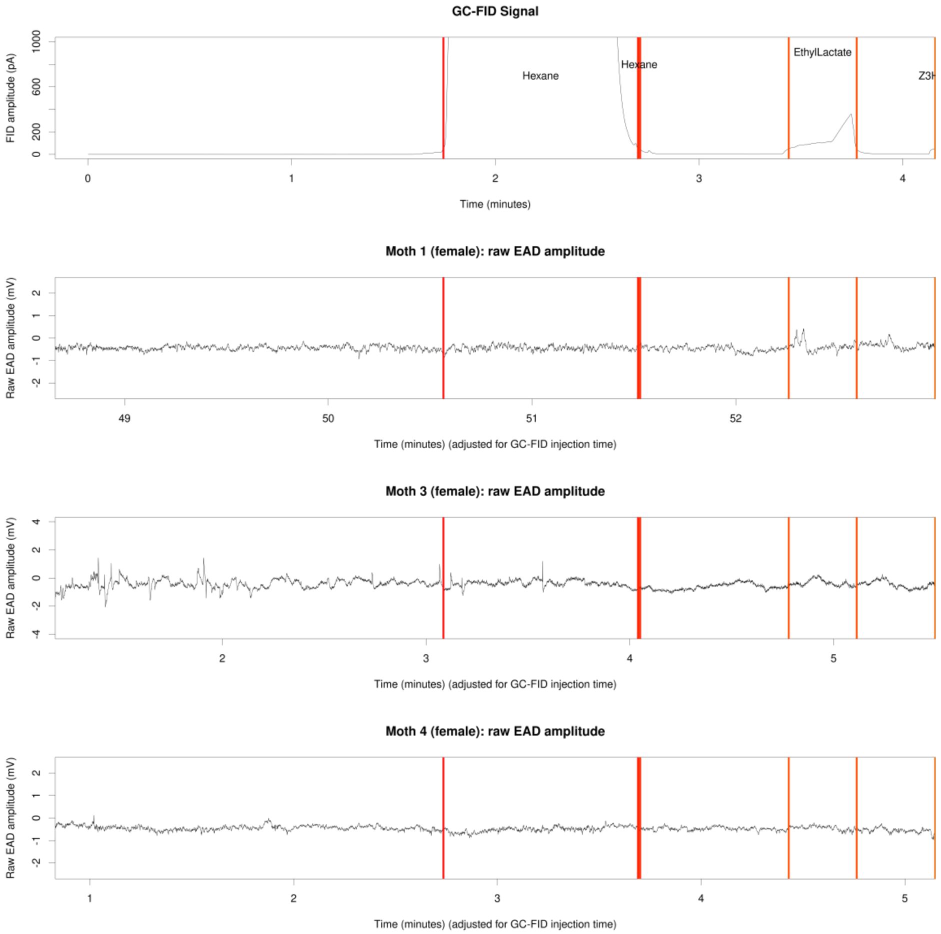


**Figure S1:** Raw EAD amplitude for three female hawkmoths, 0-4 minutes of GC-FID elapsed time. Top pane: GC-FID trace annotated with GC-FID peak starts and ends (colored vertical lines) and GC-FID peak identities. Bottom three panes: individual female hawkmoth EAD traces annotated with GC-FID peak starts and ends (colored vertical lines).

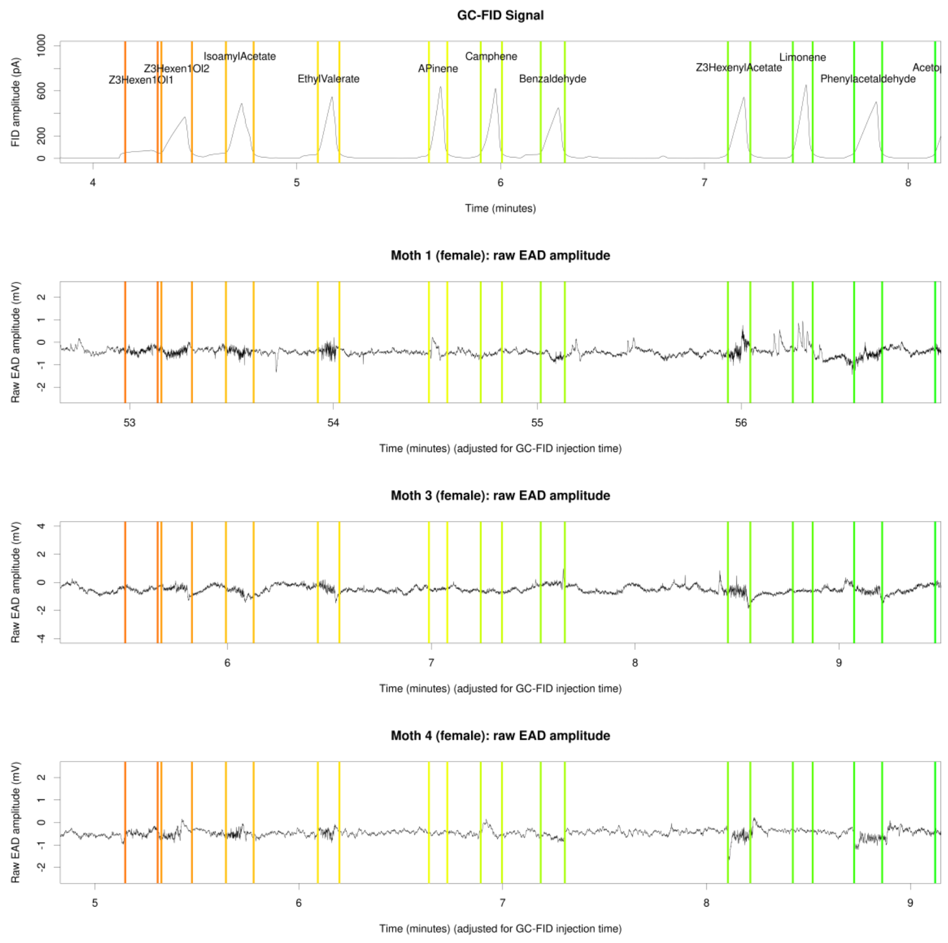


**Figure S2:** Raw EAD amplitude for three female hawkmoths, 4-8 minutes of GC-FID elapsed time. Top pane: GC-FID trace annotated with GC-FID peak starts and ends (colored vertical lines) and GC-FID peak identities. Bottom three panes: individual female hawkmoth EAD traces annotated with GC-FID peak starts and ends (colored vertical lines).

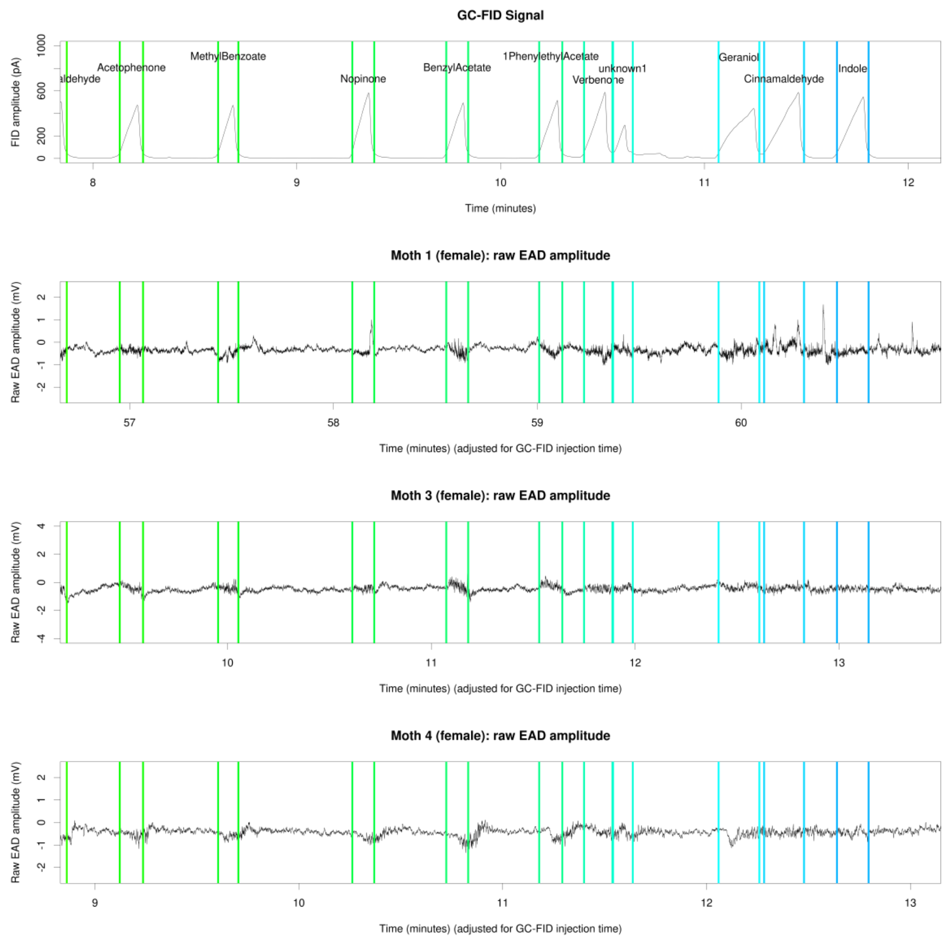


**Figure S3:** Raw EAD amplitude for three female hawkmoths, 8-12 minutes of GC-FID elapsed time. Top pane: GC-FID trace annotated with GC-FID peak starts and ends (colored vertical lines) and GC-FID peak identities. Bottom three panes: individual female hawkmoth EAD traces annotated with GC-FID peak starts and ends (colored vertical lines).

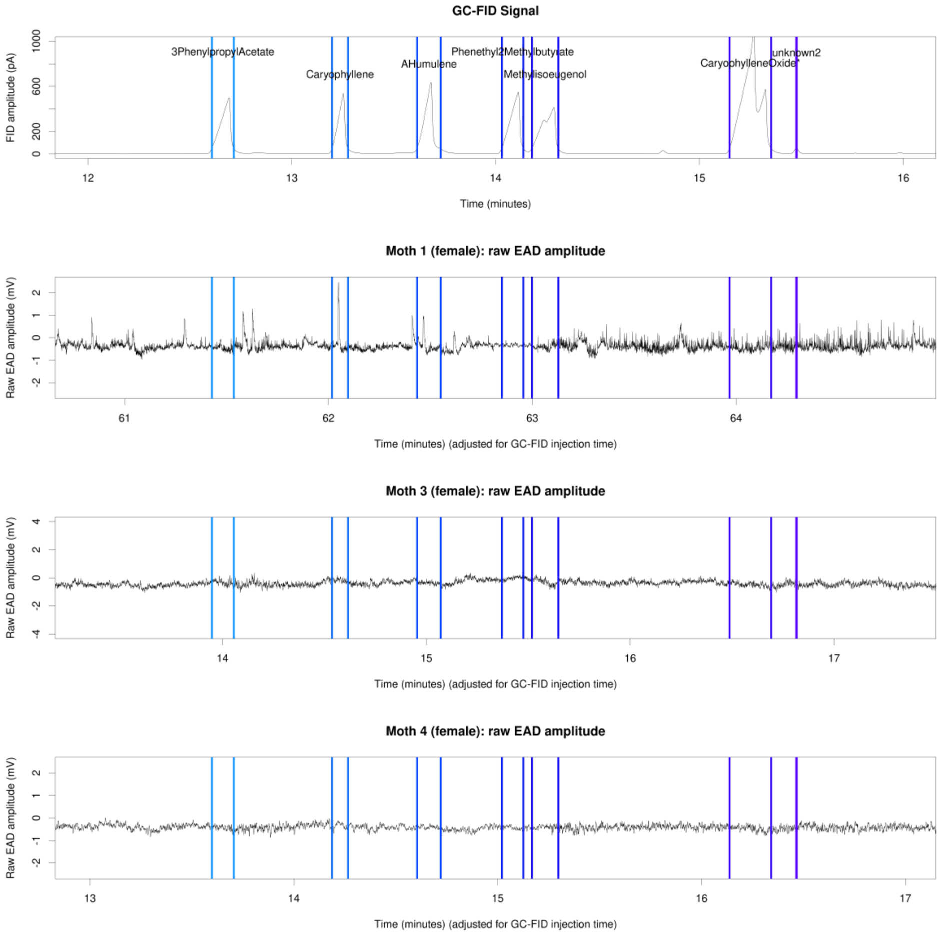


**Figure S4:** Raw EAD amplitude for three female hawkmoths, 12-16 minutes of GC-FID elapsed time. Top pane: GC-FID trace annotated with GC-FID peak starts and ends (colored vertical lines) and GC-FID peak identities. Bottom three panes: individual female hawkmoth EAD traces annotated with GC-FID peak starts and ends (colored vertical lines).

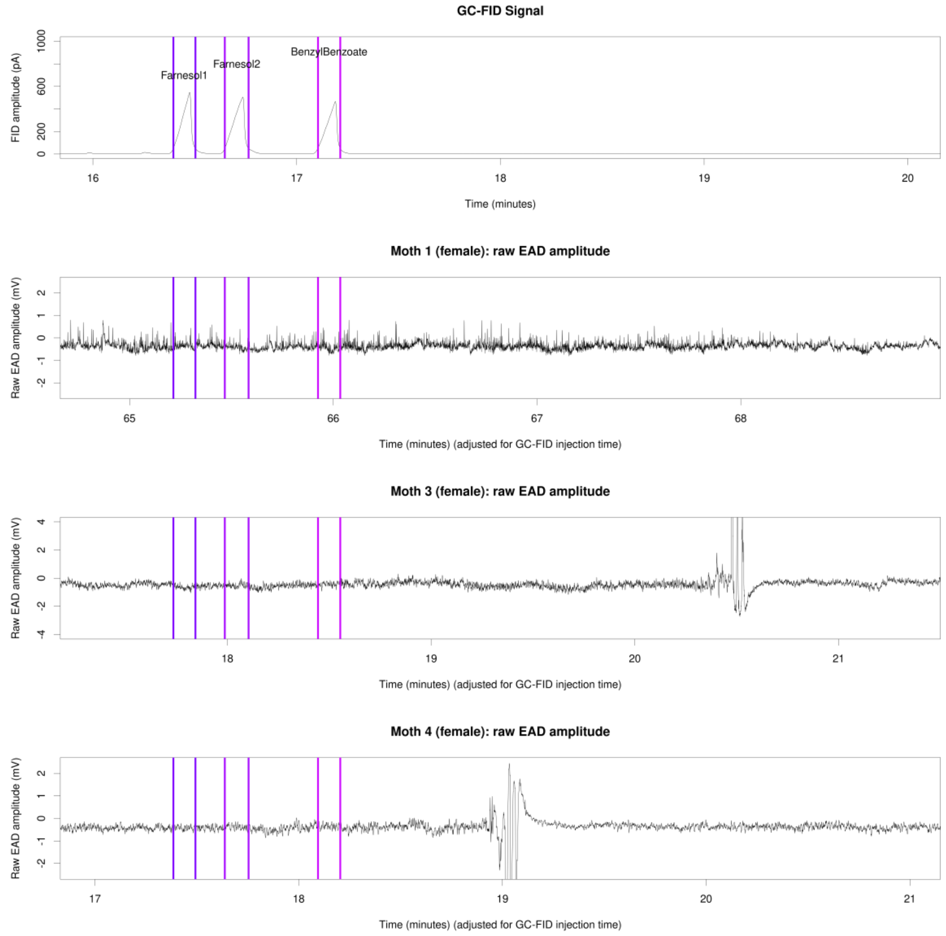


**Figure S5:** Raw EAD amplitude for three female hawkmoths, 16-20 minutes of GC-FID elapsed time. Top pane: GC-FID trace annotated with GC-FID peak starts and ends (colored vertical lines) and GC-FID peak identities. Bottom three panes: individual female hawkmoth EAD traces annotated with GC-FID peak starts and ends (colored vertical lines). The large peaks in the second two EAD traces after the benzyl benzoate interval are mixture puffs to test antennal viability.

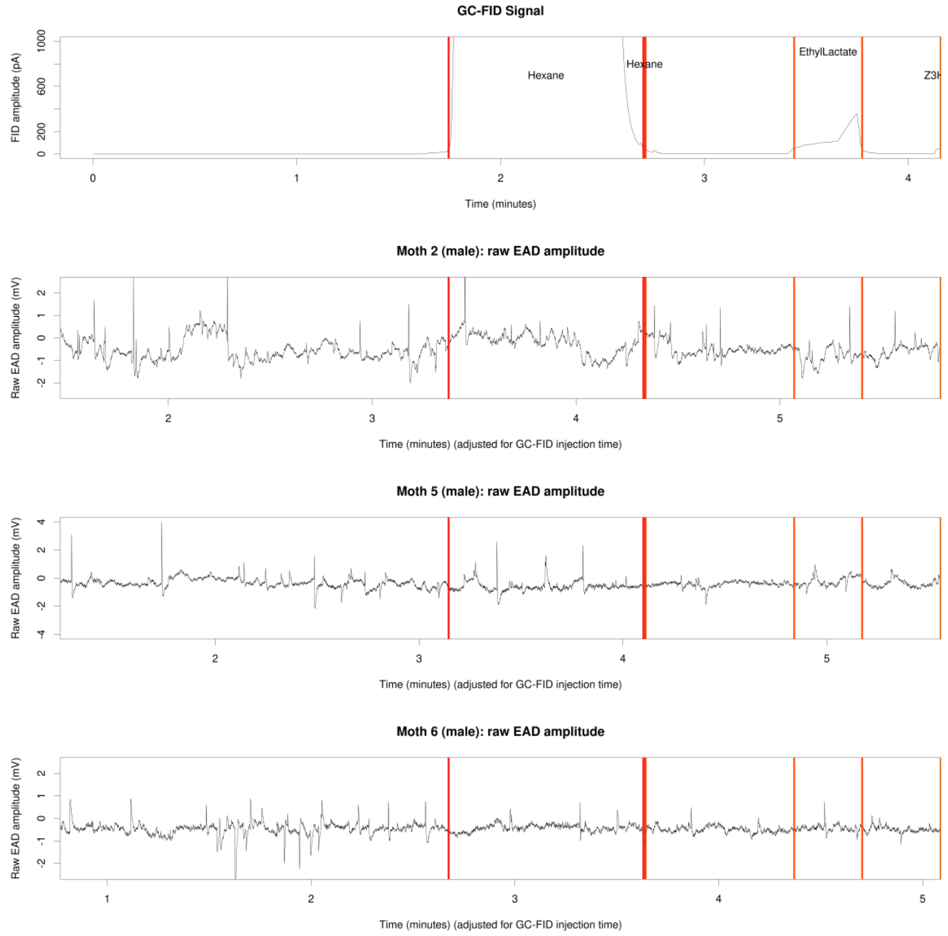


**Figure S6:** Raw EAD amplitude for three male hawkmoths, 0-4 minutes of GC-FID elapsed time. Top pane: GC-FID trace annotated with GC-FID peak starts and ends (colored vertical lines) and GC-FID peak identities. Bottom three panes: individual male hawkmoth EAD traces annotated with GC-FID peak starts and ends (colored vertical lines).

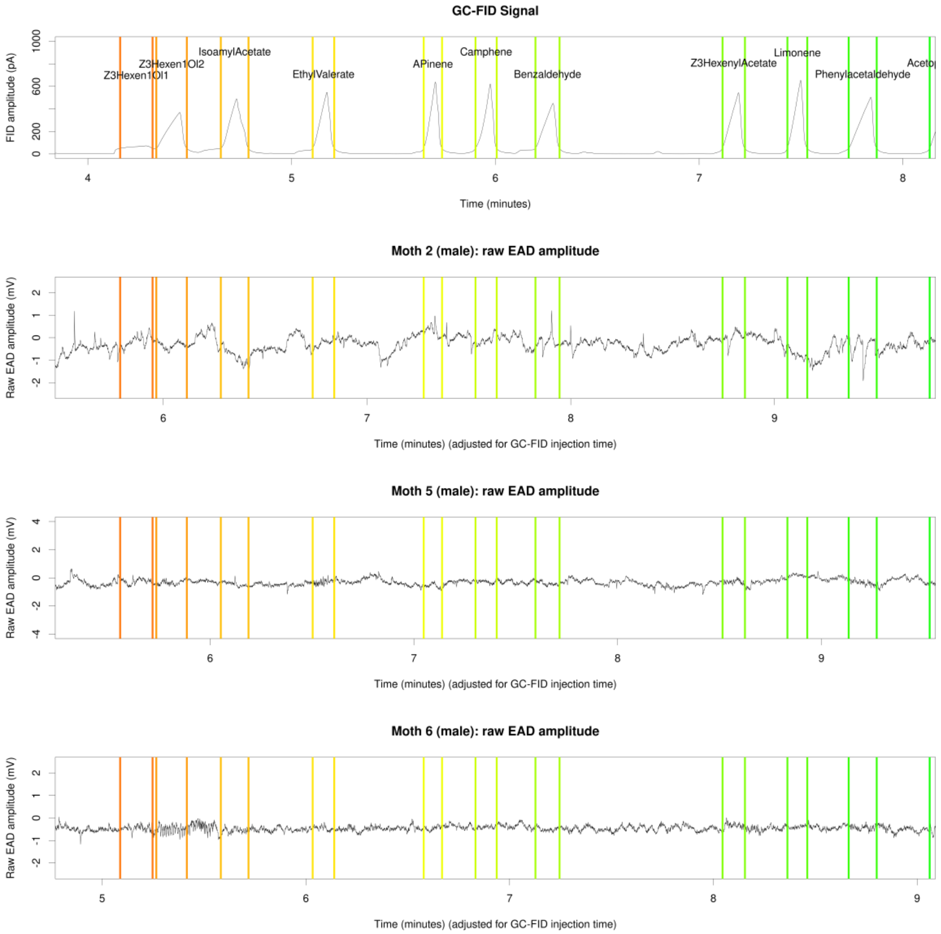


**Figure S7:** Raw EAD amplitude for three male hawkmoths, 4-8 minutes of GC-FID elapsed time. Top pane: GC-FID trace annotated with GC-FID peak starts and ends (colored vertical lines) and GC-FID peak identities. Bottom three panes: individual male hawkmoth EAD traces annotated with GC-FID peak starts and ends (colored vertical lines).

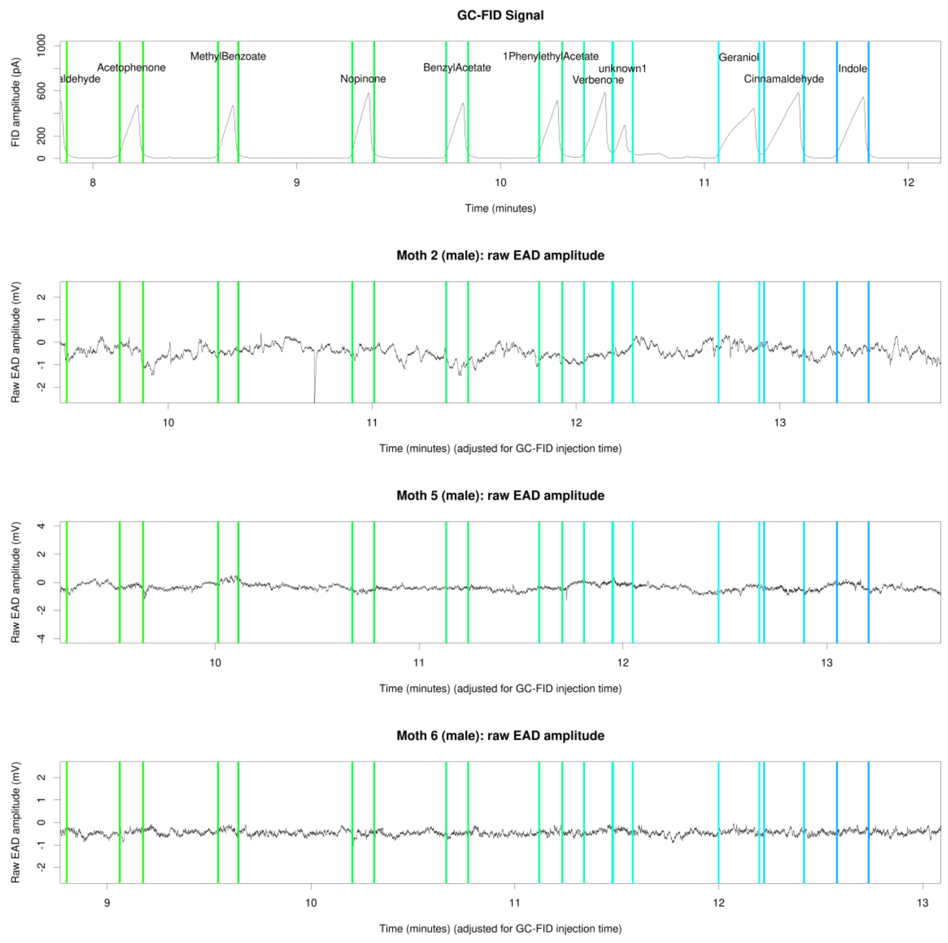


**Figure S8:** Raw EAD amplitude for three male hawkmoths, 8-12 minutes of GC-FID elapsed time. Top pane: GC-FID trace annotated with GC-FID peak starts and ends (colored vertical lines) and GC-FID peak identities. Bottom three panes: individual male hawkmoth EAD traces annotated with GC-FID peak starts and ends (colored vertical lines).

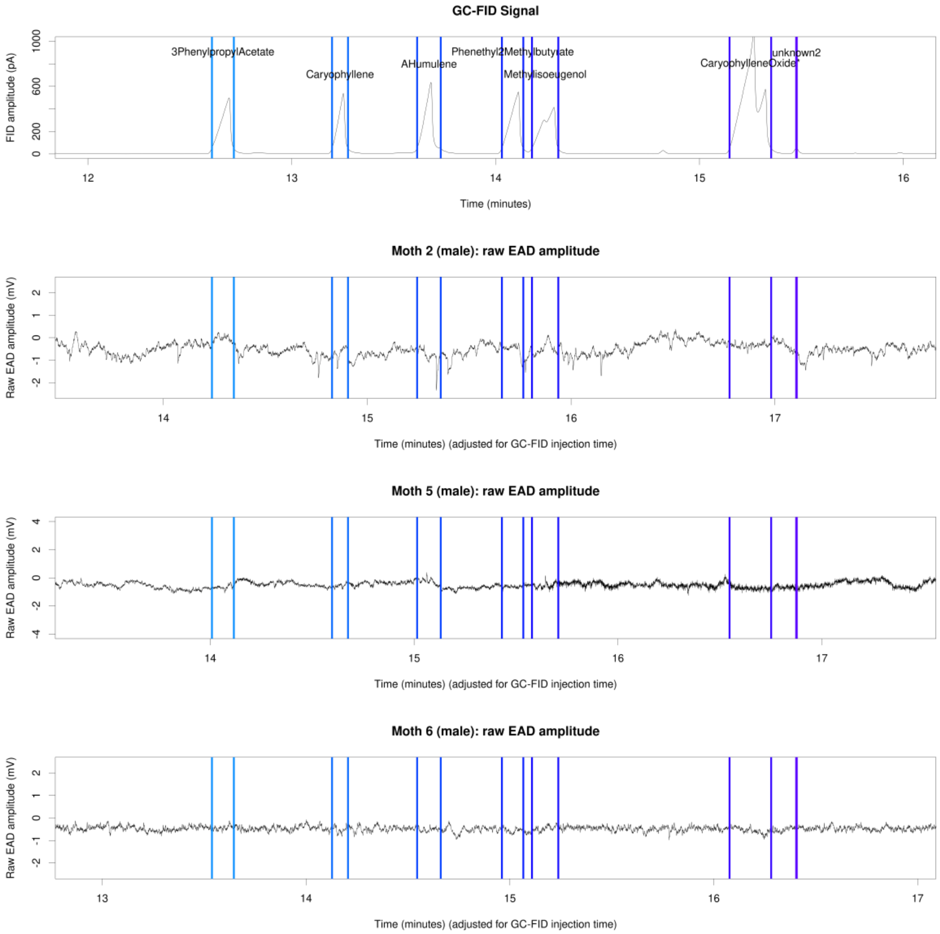


**Figure S9:** Raw EAD amplitude for three male hawkmoths, 12-16 minutes of GC-FID elapsed time. Top pane: GC-FID trace annotated with GC-FID peak starts and ends (colored vertical lines) and GC-FID peak identities. Bottom three panes: individual male hawkmoth EAD traces annotated with GC-FID peak starts and ends (colored vertical lines).

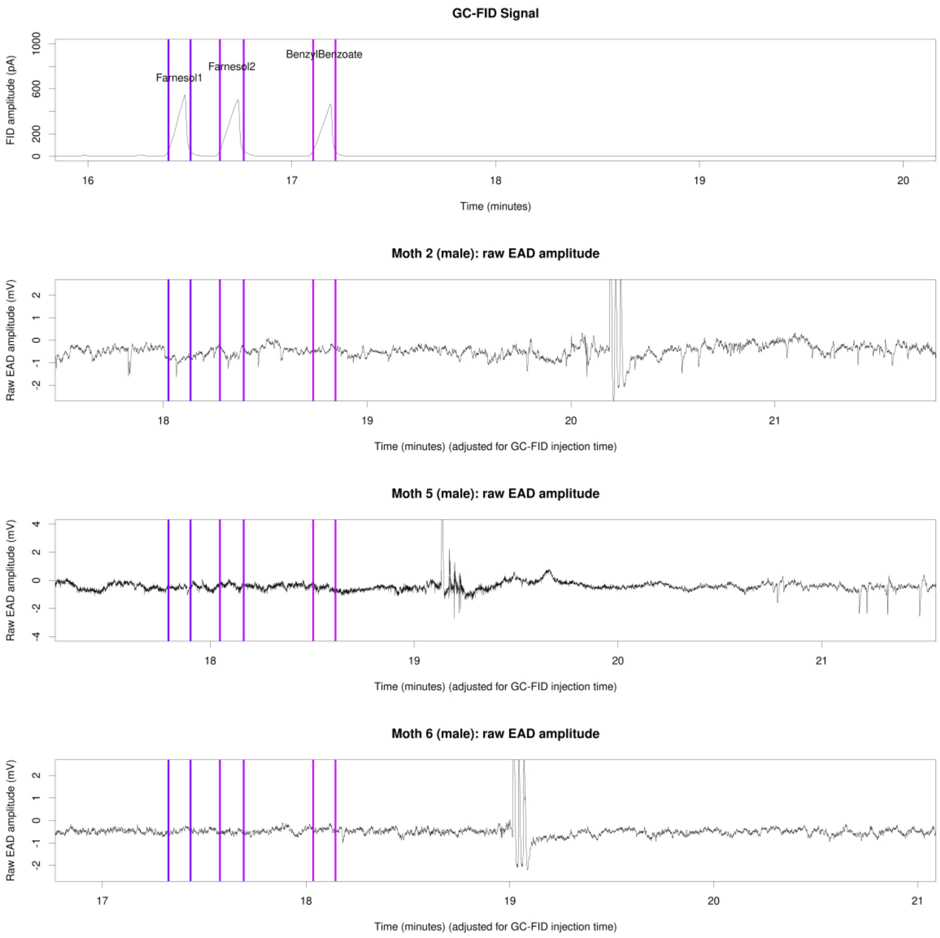


**Figure S10:** Raw EAD amplitude for three male hawkmoths, 16-20 minutes of GC-FID elapsed time. Top pane: GC-FID trace annotated with GC-FID peak starts and ends (colored vertical lines) and GC-FID peak identities. Bottom three panes: individual male hawkmoth EAD traces annotated with GC-FID peak starts and ends (colored vertical lines). The large peaks in the EAD traces after the benzyl benzoate interval are mixture puffs to test antennal viability.

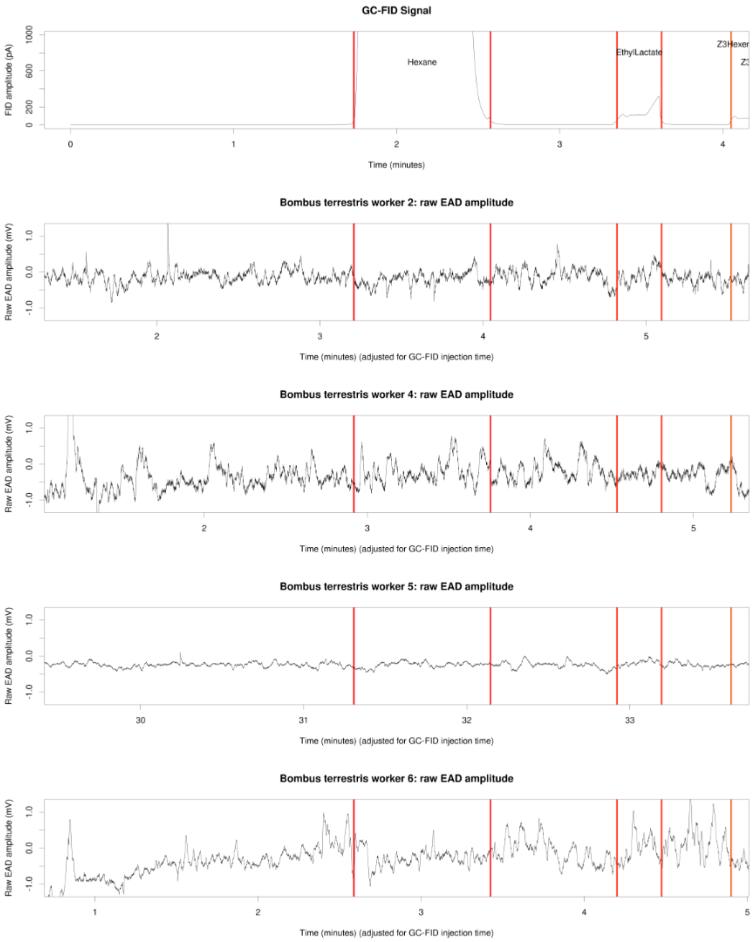


**Figure S11:** Raw EAD amplitude for four bumble bee workers, 0-4 minutes of GC-FID elapsed time. Top pane: GC-FID trace annotated with GC-FID peak starts and ends (colored vertical lines) and GC-FID peak identities. Bottom four panes: individual bumble bee worker EAD traces annotated with GC-FID peak starts and ends (colored vertical lines).

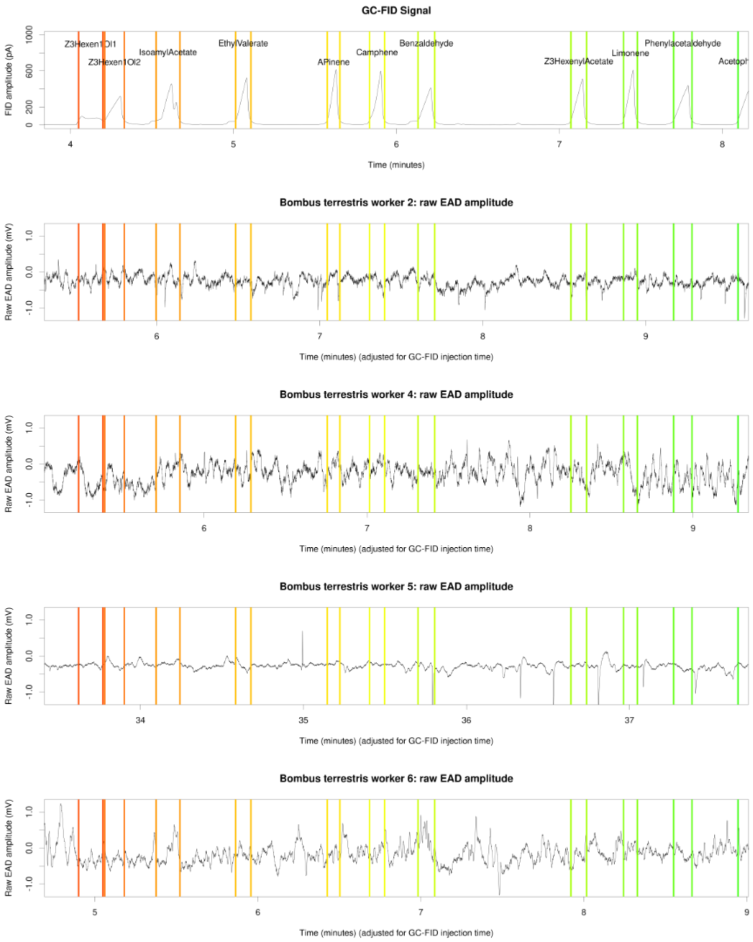


**Figure S12:** Raw EAD amplitude for four bumble bee workers, 4-8 minutes of GC-FID elapsed time. Top pane: GC-FID trace annotated with GC-FID peak starts and ends (colored vertical lines) and GC-FID peak identities. Bottom four panes: individual bumble bee worker EAD traces annotated with GC-FID peak starts and ends (colored vertical lines).

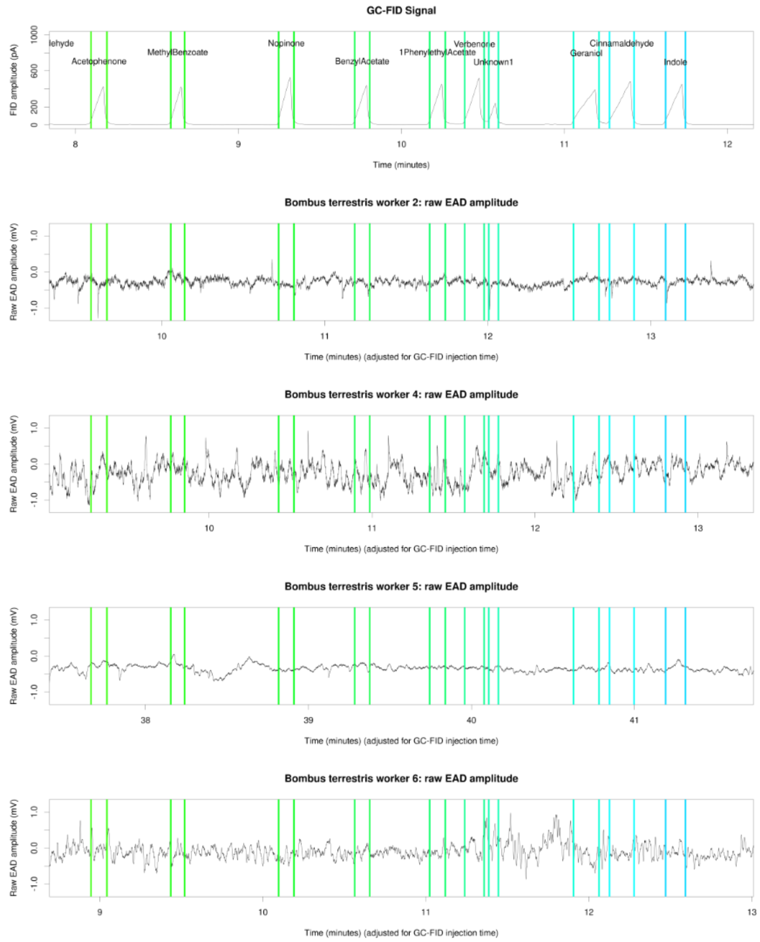


**Figure S13:** Raw EAD amplitude for four bumble bee workers, 8-12 minutes of GC-FID elapsed time. Top pane: GC-FID trace annotated with GC-FID peak starts and ends (colored vertical lines) and GC-FID peak identities. Bottom four panes: individual bumble bee worker EAD traces annotated with GC-FID peak starts and ends (colored vertical lines).

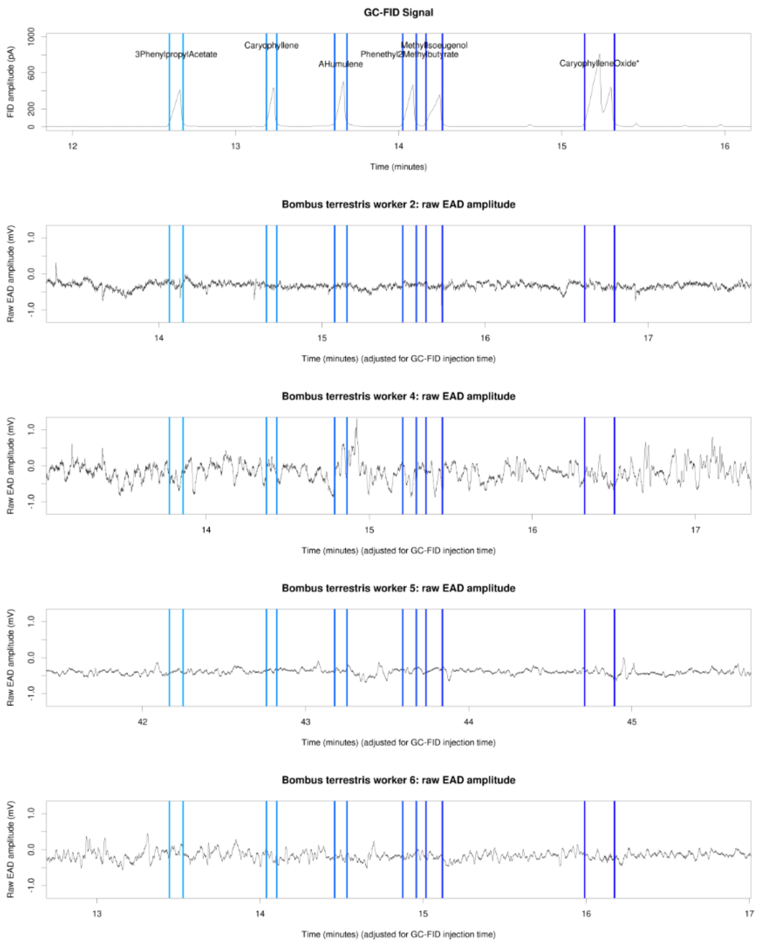


**Figure S14:** Raw EAD amplitude for four bumble bee workers, 12-16 minutes of GC-FID elapsed time. Top pane: GC-FID trace annotated with GC-FID peak starts and ends (colored vertical lines) and GC-FID peak identities. Bottom four panes: individual bumble bee worker EAD traces annotated with GC-FID peak starts and ends (colored vertical lines).

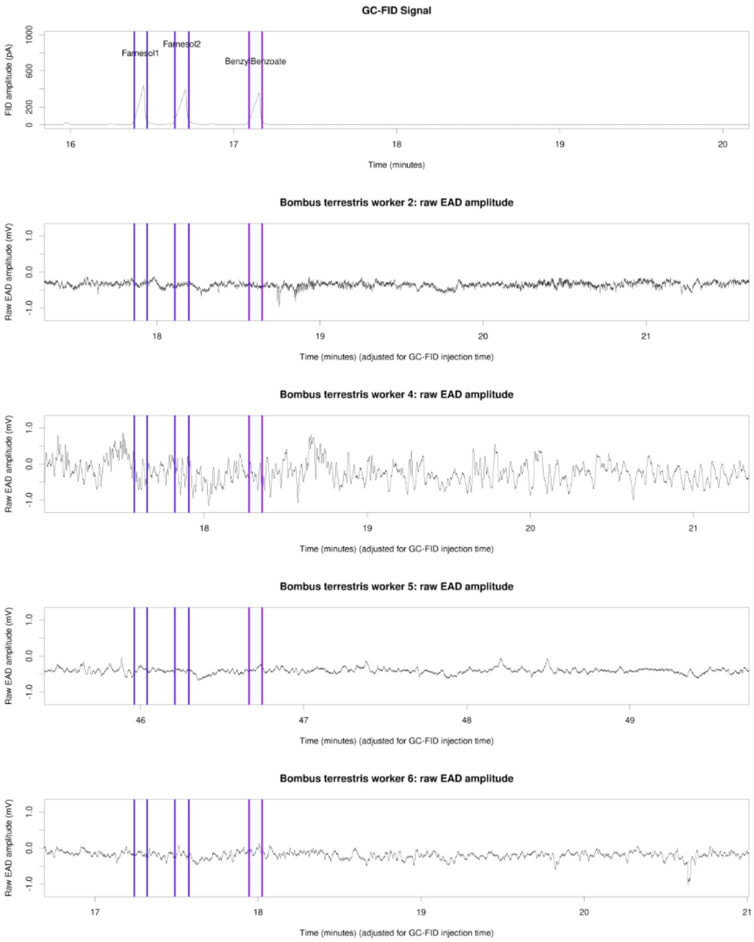


**Figure S15:** Raw EAD amplitude for four bumble bee workers, 16-20 minutes of GC-FID elapsed time. Top pane: GC-FID trace annotated with GC-FID peak starts and ends (colored vertical lines) and GC-FID peak identities. Bottom four panes: individual bumble bee worker EAD traces annotated with GC-FID peak starts and ends (colored vertical lines).

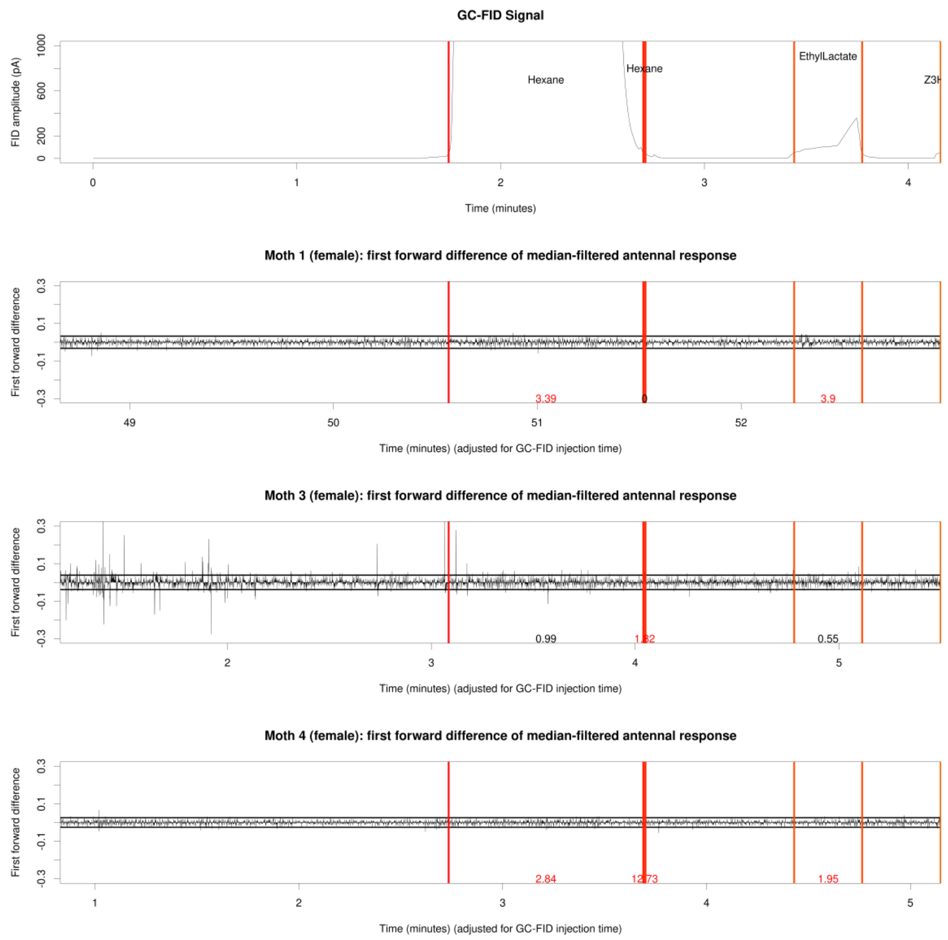


**Figure S16:** First forward difference-processed EAD amplitude for three female hawkmoths, 0-4 minutes of GC-FID elapsed time. Top pane: GC-FID trace annotated with GC-FID peak starts and ends (colored vertical lines) and GC-FID peak identities. Bottom three panes: individual female hawkmoth EAD traces annotated with GC-FID peak starts and ends (colored vertical lines) and significant GC-EAD spike hit frequency (spikes/second), with red indicating frequency over the background threshold.

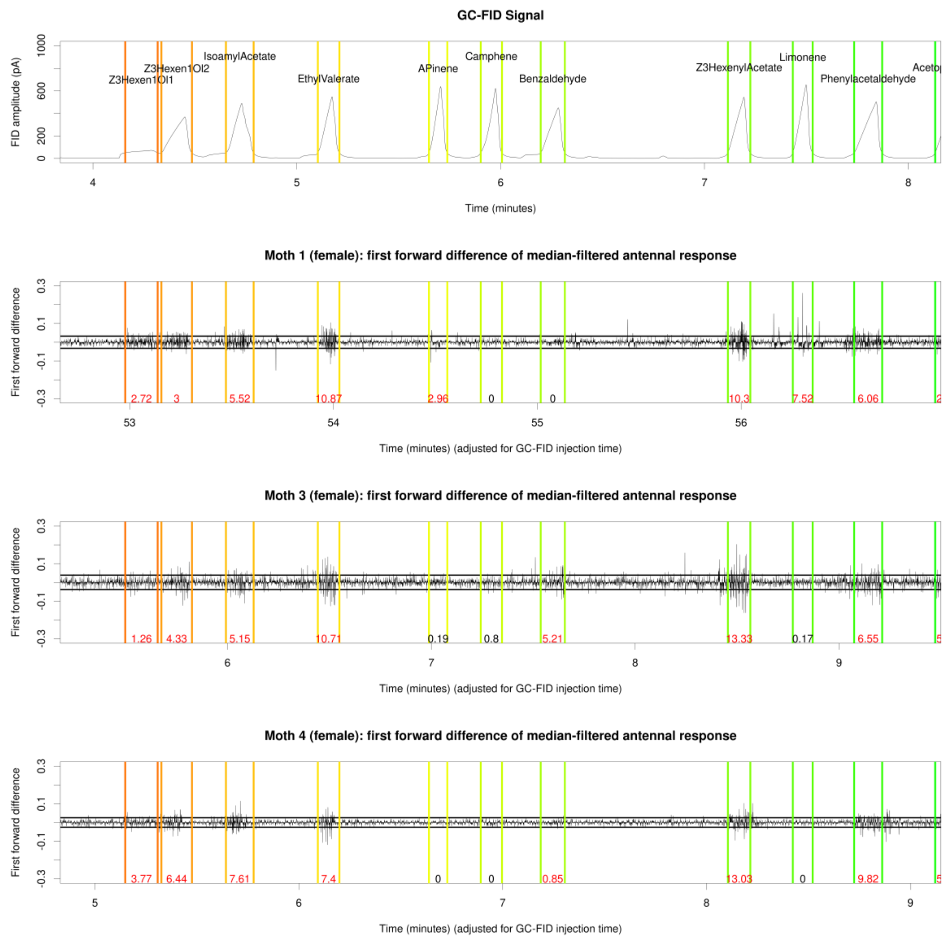


**Figure S17:** First forward difference-processed EAD amplitude for three female hawkmoths, 4-8 minutes of GC-FID elapsed time. Top pane: GC-FID trace annotated with GC-FID peak starts and ends (colored vertical lines) and GC-FID peak identities. Bottom three panes: individual female hawkmoth EAD traces annotated with GC-FID peak starts and ends (colored vertical lines) and significant GC-EAD spike hit frequency (spikes/second), with red indicating frequency over the background threshold.

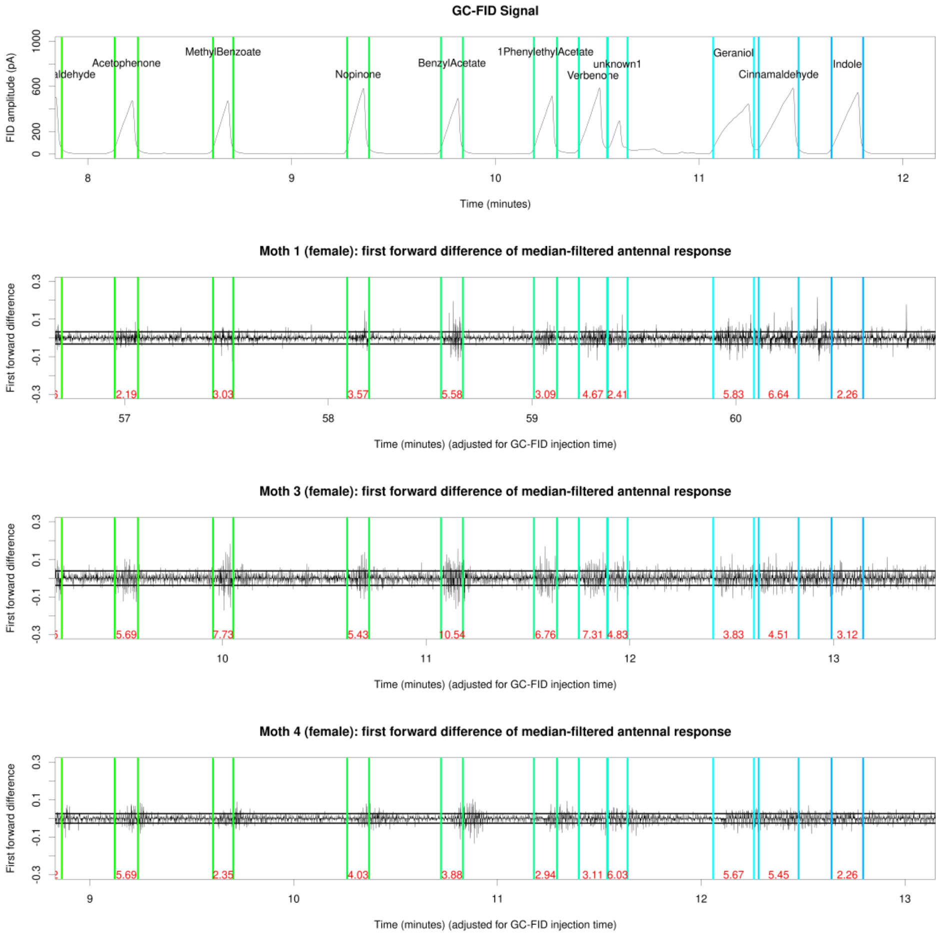


**Figure S18:** First forward difference-processed EAD amplitude for three female hawkmoths, 8-12 minutes of GC-FID elapsed time. Top pane: GC-FID trace annotated with GC-FID peak starts and ends (colored vertical lines) and GC-FID peak identities. Bottom three panes: individual female hawkmoth EAD traces annotated with GC-FID peak starts and ends (colored vertical lines) and significant GC-EAD spike hit frequency (spikes/second), with red indicating frequency over the background threshold.

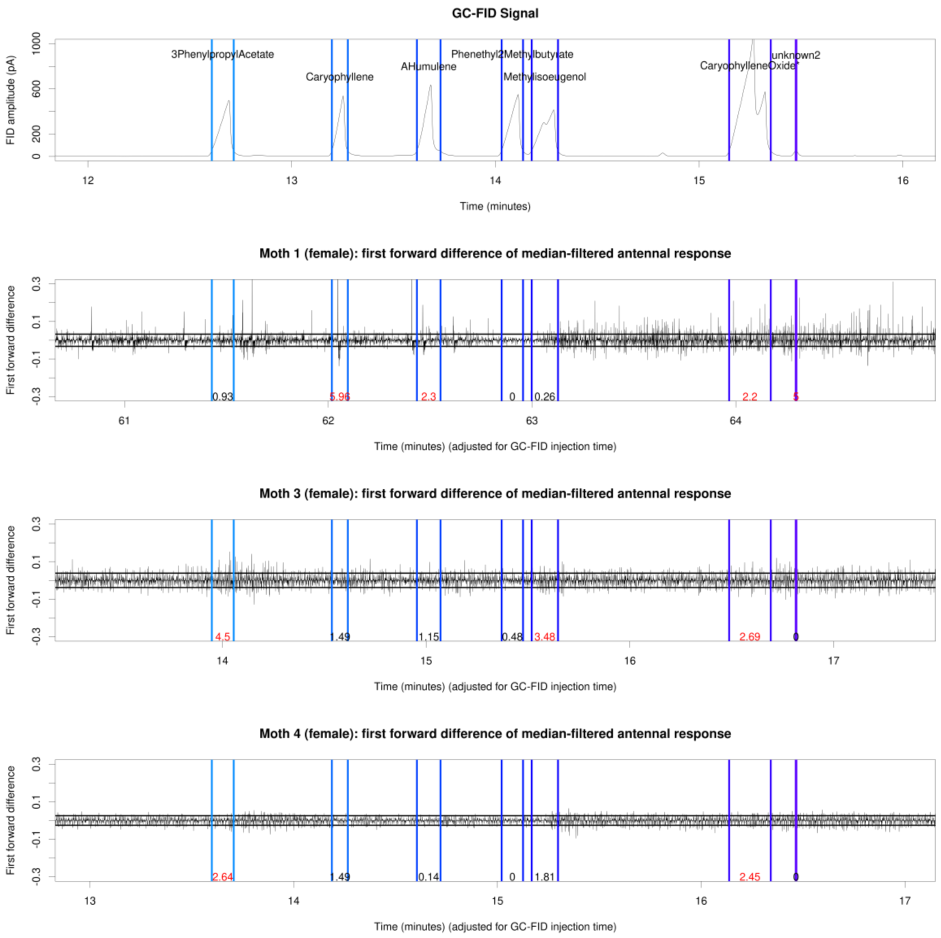


**Figure S19:** First forward difference-processed EAD amplitude for three female hawkmoths, 12-16 minutes of GC-FID elapsed time. Top pane: GC-FID trace annotated with GC-FID peak starts and ends (colored vertical lines) and GC-FID peak identities. Bottom three panes: individual female hawkmoth EAD traces annotated with GC-FID peak starts and ends (colored vertical lines) and significant GC-EAD spike hit frequency (spikes/second), with red indicating frequency over the background threshold.

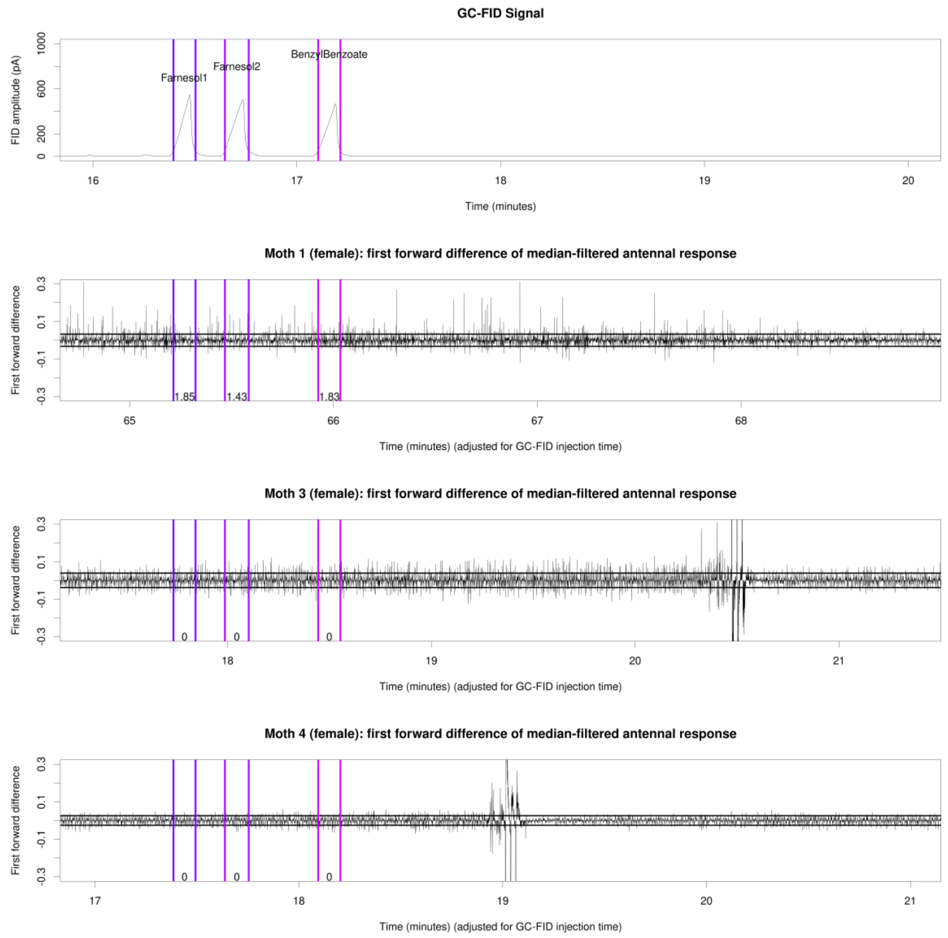


**Figure S20:** First forward difference-processed EAD amplitude for three female hawkmoths, 16-20 minutes of GC-FID elapsed time. Top pane: GC-FID trace annotated with GC-FID peak starts and ends (colored vertical lines) and GC-FID peak identities. Bottom three panes: individual female hawkmoth EAD traces annotated with GC-FID peak starts and ends (colored vertical lines) and significant GC-EAD spike hit frequency (spikes/second), with red indicating frequency over the background threshold. The large peaks in the second two EAD traces after the benzyl benzoate interval are mixture puffs to test antennal viability.

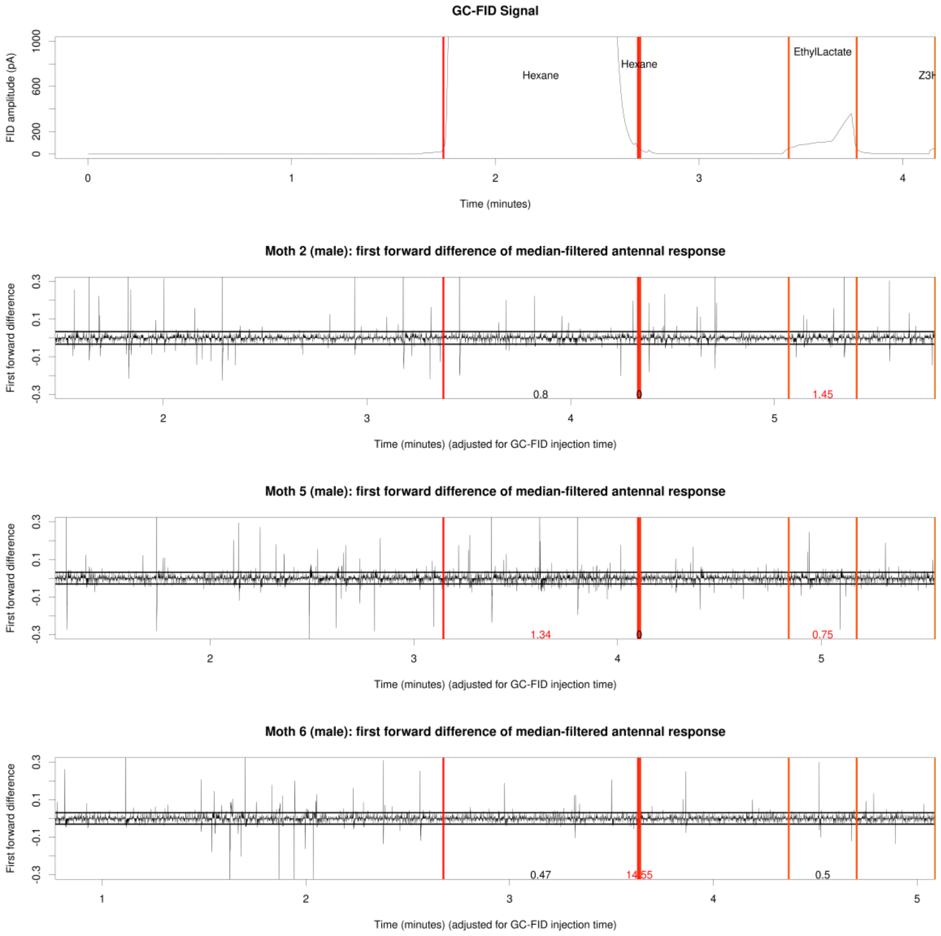


**Figure S21:** First forward differ GC-FID ence-processed EAD amplitude for three male hawkmoths, 0-4 minutes of GC-FID elapsed time. Top pane: GC-FID trace annotated with GC-FID peak starts and ends (colored vertical lines) and GC-FID peak identities. Bottom three panes: individual male hawkmoth EAD traces annotated with GC-FID peak starts and ends (colored vertical lines) and significant GC-EAD spike hit frequency (spikes/second), with red indicating frequency over the background threshold.

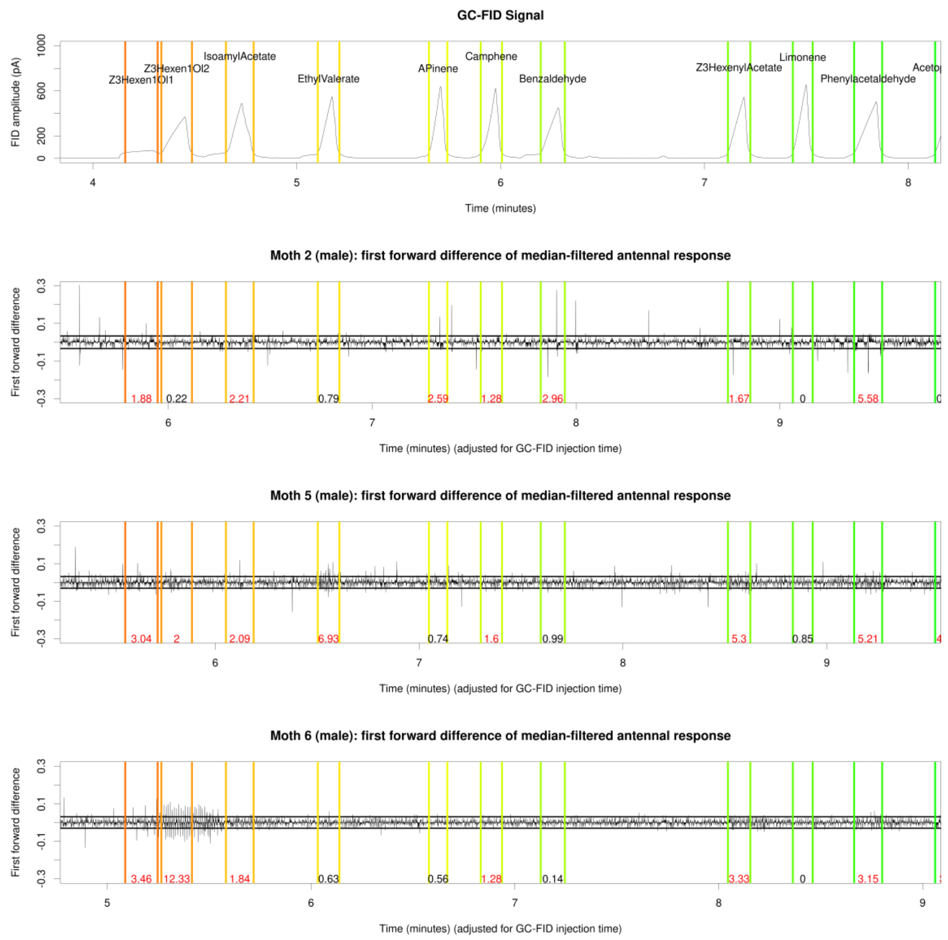


**Figure S22:** First forward difference-processed EAD amplitude for three male hawkmoths, 4-8 minutes of GC-FID elapsed time. Top pane: GC-FID trace annotated with GC-FID peak starts and ends (colored vertical lines) and GC-FID peak identities. Bottom three panes: individual male hawkmoth EAD traces annotated with GC-FID peak starts and ends (colored vertical lines) and significant GC-EAD spike hit frequency (spikes/second), with red indicating frequency over the background threshold.

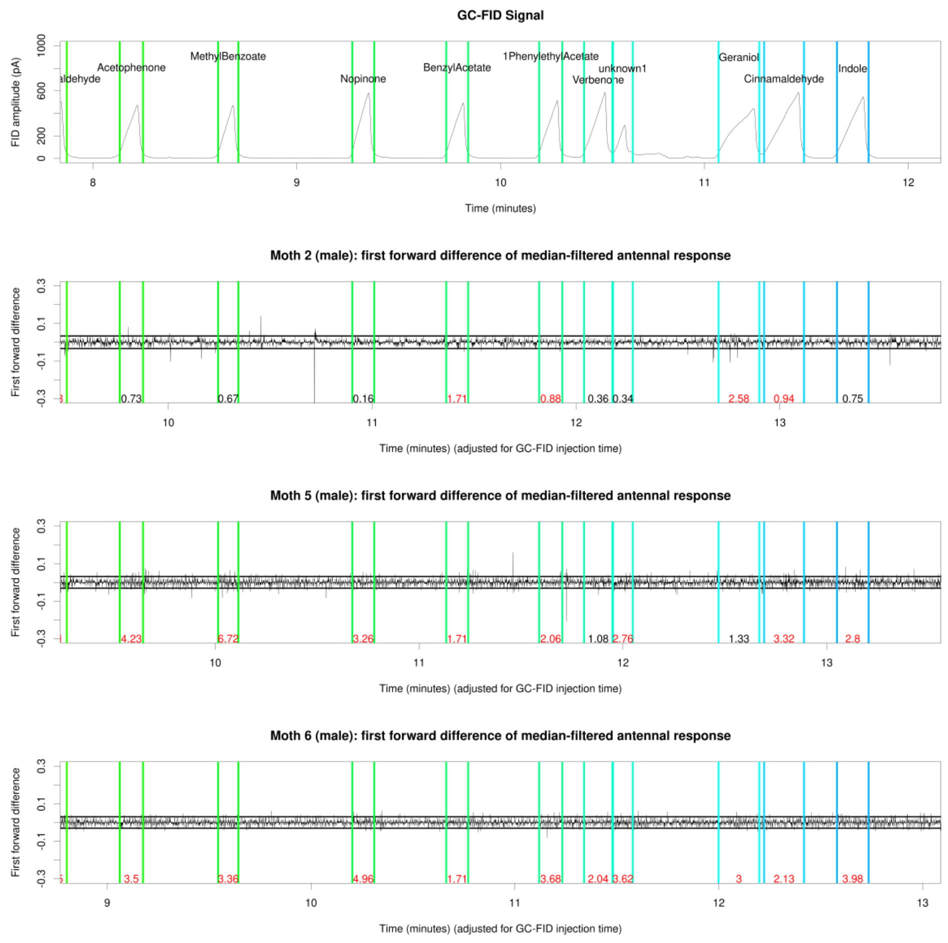


**Figure S23:** First forward difference-processed EAD amplitude for three male hawkmoths, 8-12 minutes of GC-FID elapsed time. Top pane: GC-FID trace annotated with GC-FID peak starts and ends (colored vertical lines) and GC-FID peak identities. Bottom three panes: individual male hawkmoth EAD traces annotated with GC-FID peak starts and ends (colored vertical lines) and significant GC-EAD spike hit frequency (spikes/second), with red indicating frequency over the background threshold.

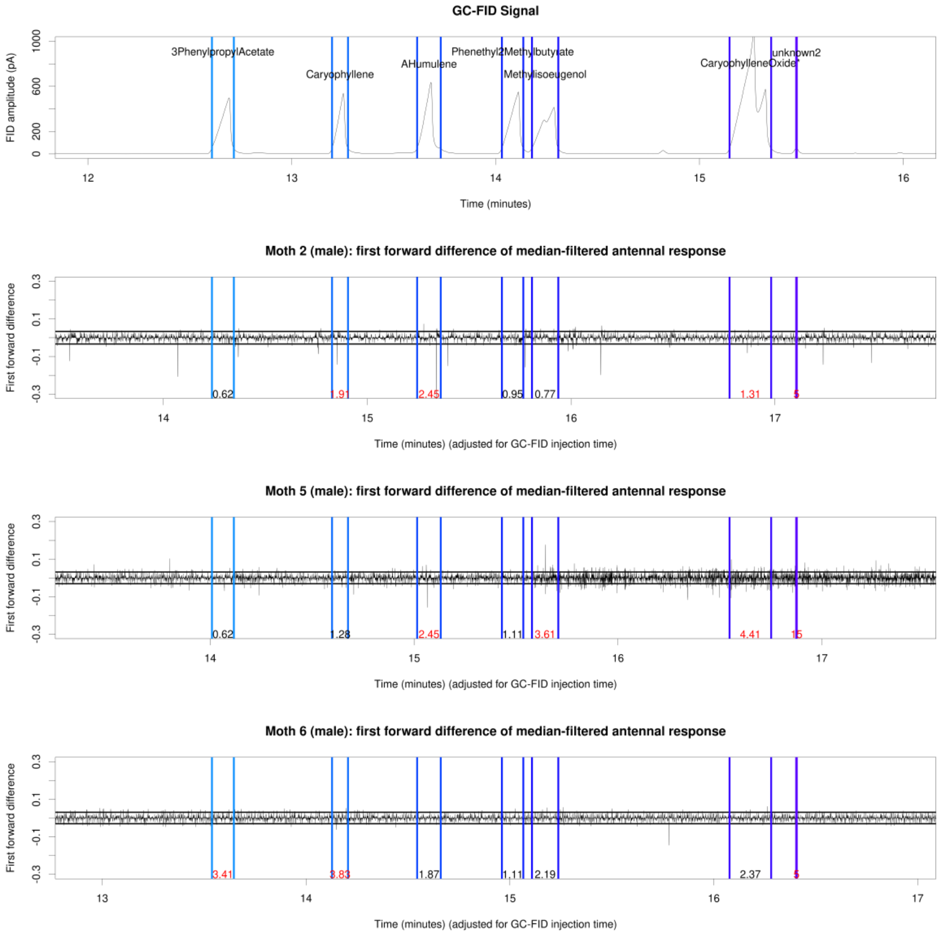


**Figure S24:** First forward difference-processed EAD amplitude for three male hawkmoths, 12-16 minutes of GC-FID elapsed time. Top pane: GC-FID trace annotated with GC-FID peak starts and ends (colored vertical lines) and GC-FID peak identities. Bottom three panes: individual male hawkmoth EAD traces annotated with GC-FID peak starts and ends (colored vertical lines) and significant GC-EAD spike hit frequency (spikes/second), with red indicating frequency over the background threshold.

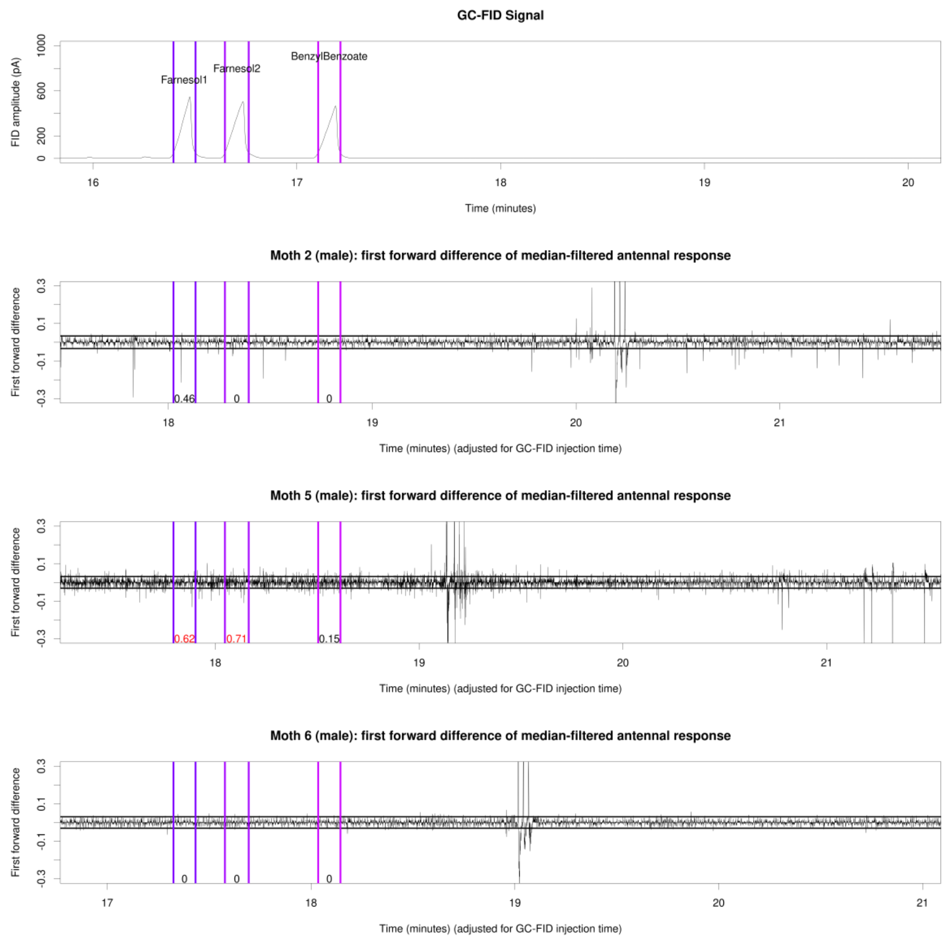


**Figure S25:** First forward difference-processed EAD amplitude for three male hawkmoths, 16-20 minutes of GC-FID elapsed time. Top pane: GC-FID trace annotated with GC-FID peak starts and ends (colored vertical lines) and GC-FID peak identities. Bottom three panes: individual male hawkmoth EAD traces annotated with GC-FID peak starts and ends (colored vertical lines) and significant GC-EAD spike hit frequency (spikes/second), with red indicating frequency over the background threshold. The large peaks in the EAD traces after the benzyl benzoate interval are mixture puffs to test antennal viability.

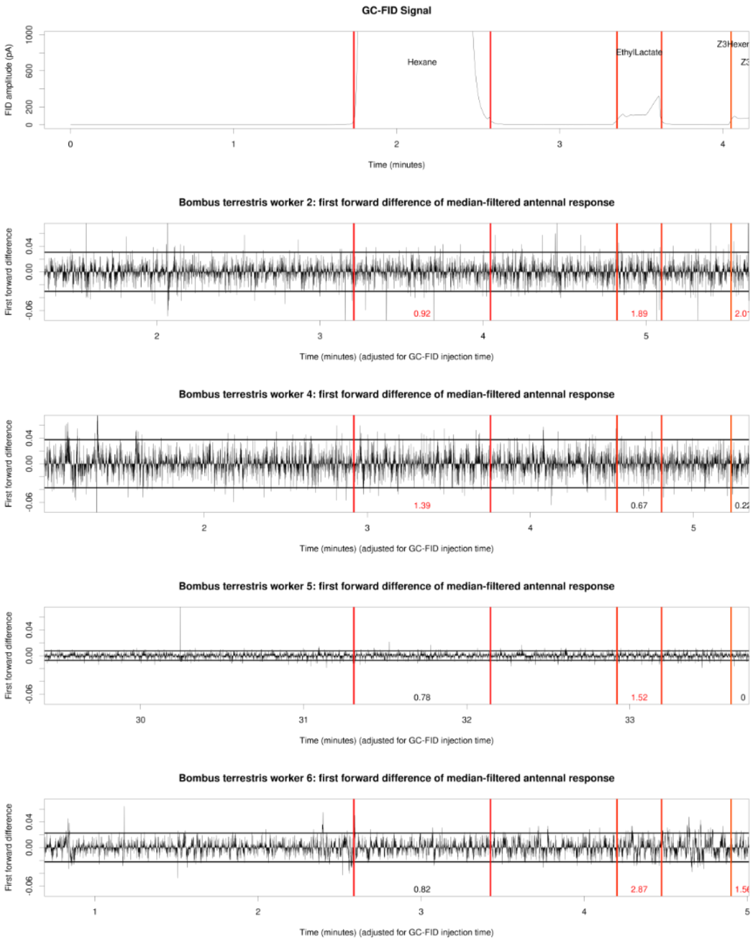


**Figure S26:** First forward difference-processed EAD amplitude for four bumble bee workers, 0-4 minutes of GC-FID elapsed time. Top pane: GC-FID trace annotated with GC-FID peak starts and ends (colored vertical lines) and GC-FID peak identities. Bottom four panes: individual bumble bee worker EAD traces annotated with GC-FID peak starts and ends (colored vertical lines) and significant GC-EAD spike hit frequency (spikes/second), with red indicating frequency over the background threshold.

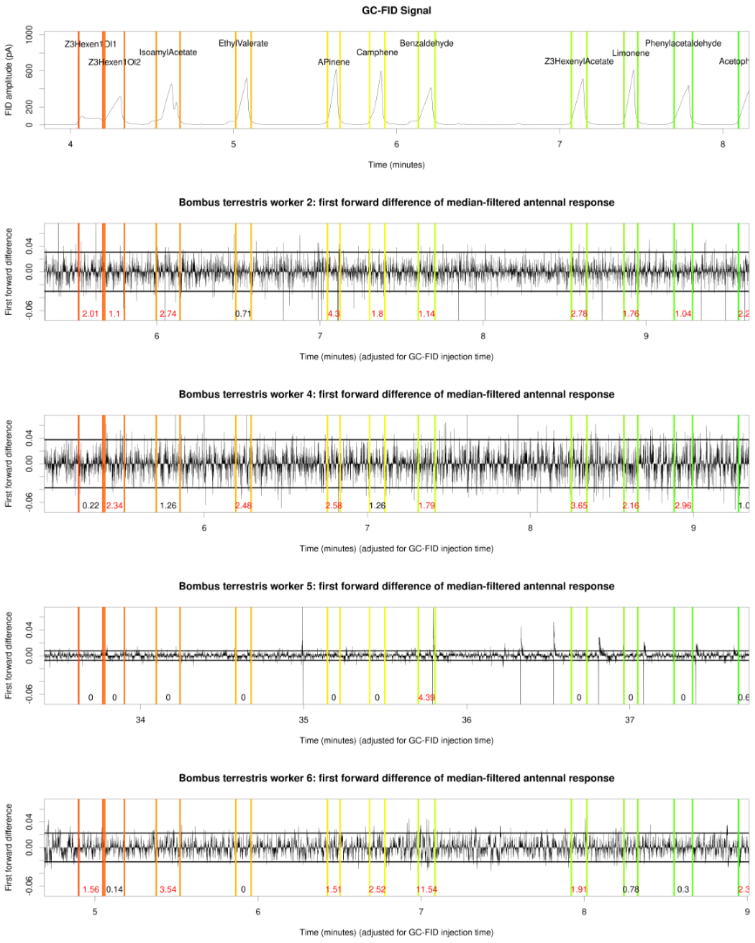


**Figure S27:** First forward difference-processed EAD amplitude for four bumble bee workers, 4-8 minutes of GC-FID elapsed time. Top pane: GC-FID trace annotated with GC-FID peak starts and ends (colored vertical lines) and GC-FID peak identities. Bottom four panes: individual bumble bee worker EAD traces annotated with GC-FID peak starts and ends (colored vertical lines) and significant GC-EAD spike hit frequency (spikes/second), with red indicating frequency over the background threshold.

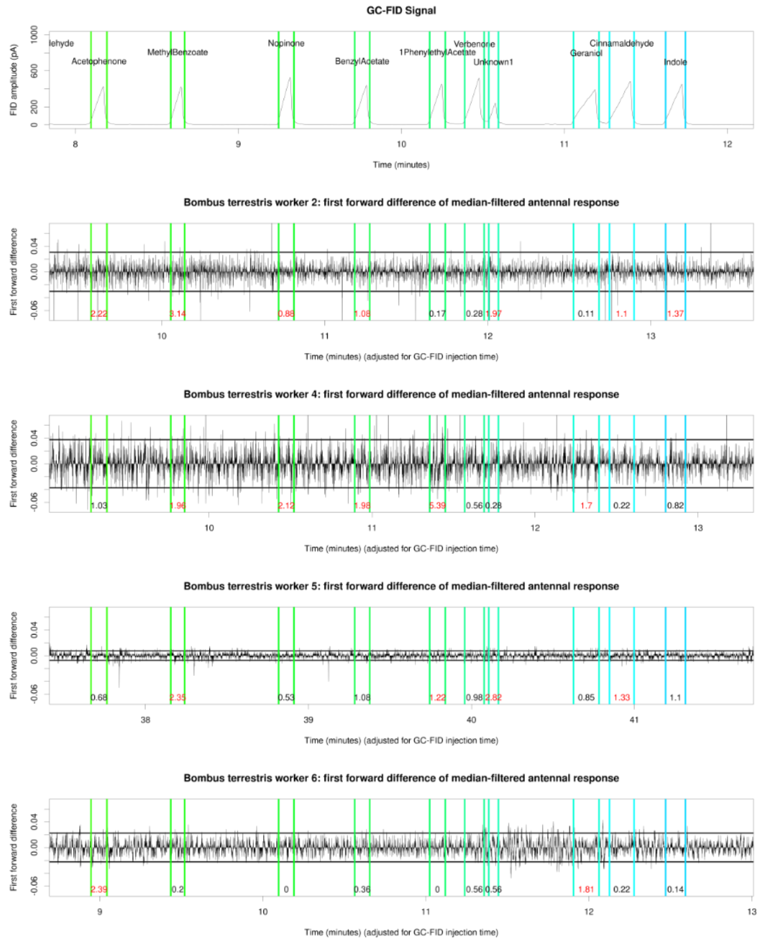


**Figure S28:** First forward difference-processed EAD amplitude for four bumble bee workers, 8-12 minutes of GC-FID elapsed time. Top pane: GC-FID trace annotated with GC-FID peak starts and ends (colored vertical lines) and GC-FID peak identities. Bottom four panes: individual bumble bee worker EAD traces annotated with GC-FID peak starts and ends (colored vertical lines) and significant GC-EAD spike hit frequency (spikes/second), with red indicating frequency over the background threshold.

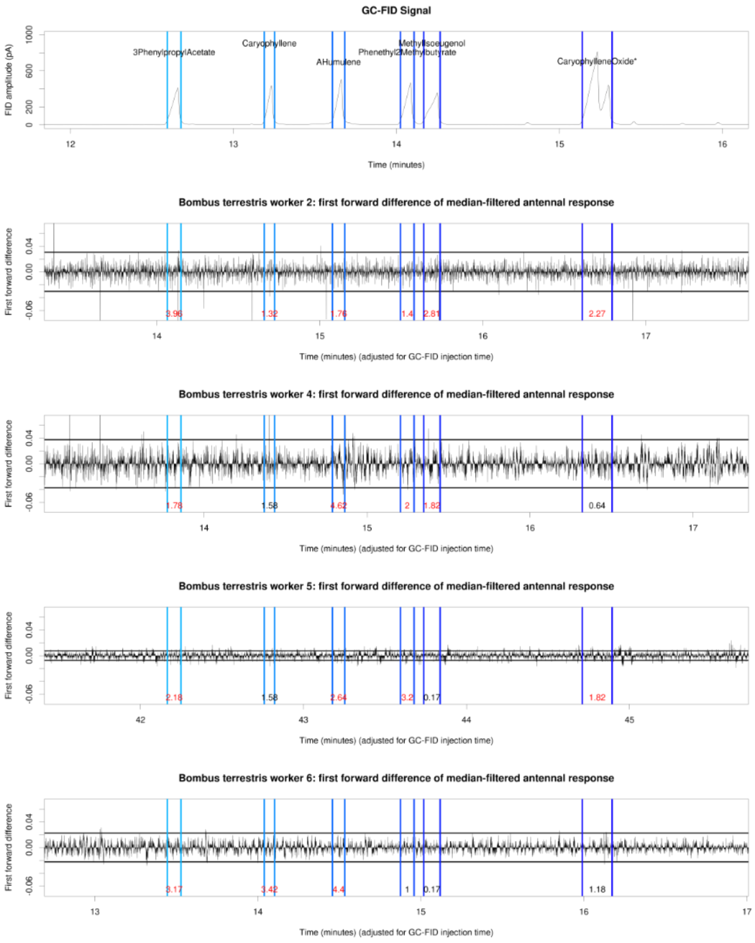


**Figure S29:** First forward difference-processed EAD amplitude for four bumble bee workers, 12-16 minutes of GC-FID elapsed time. Top pane: GC-FID trace annotated with GC-FID peak starts and ends (colored vertical lines) and GC-FID peak identities. Bottom four panes: individual bumble bee worker EAD traces annotated with GC-FID peak starts and ends (colored vertical lines) and significant GC-EAD spike hit frequency (spikes/second), with red indicating frequency over the background threshold.

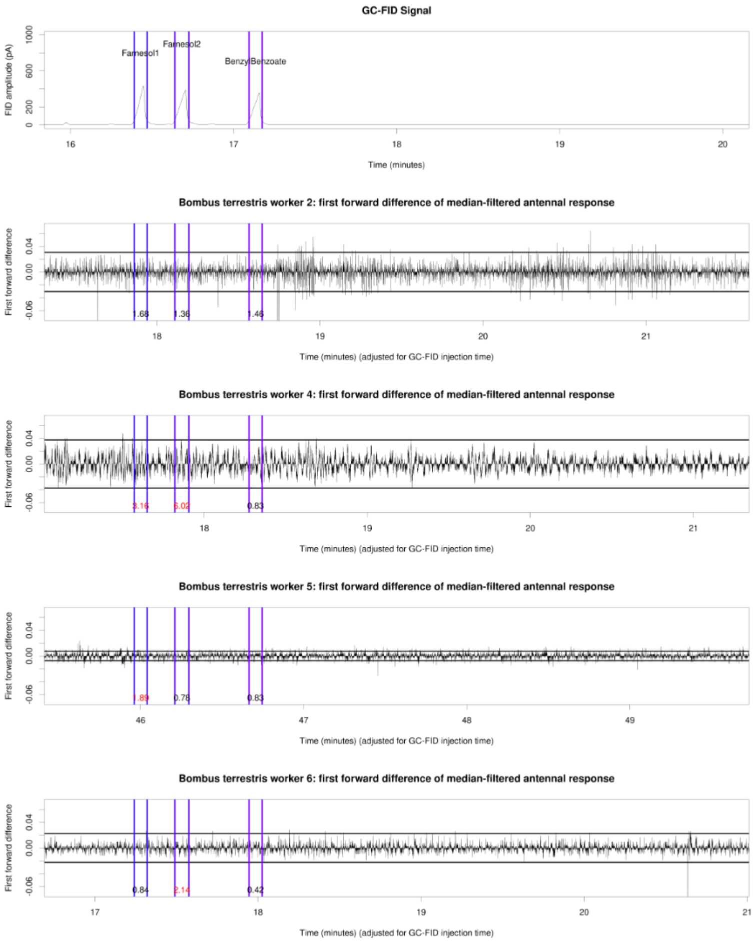


**Figure S30:** First forward difference-processed EAD amplitude for four bumble bee workers, 16-20 minutes of GC-FID elapsed time. Top pane: GC-FID trace annotated with GC-FID peak starts and ends (colored vertical lines) and GC-FID peak identities. Bottom four panes: individual bumble bee worker EAD traces annotated with GC-FID peak starts and ends (colored vertical lines) and significant GC-EAD spike hit frequency (spikes/second), with red indicating frequency over the background threshold.

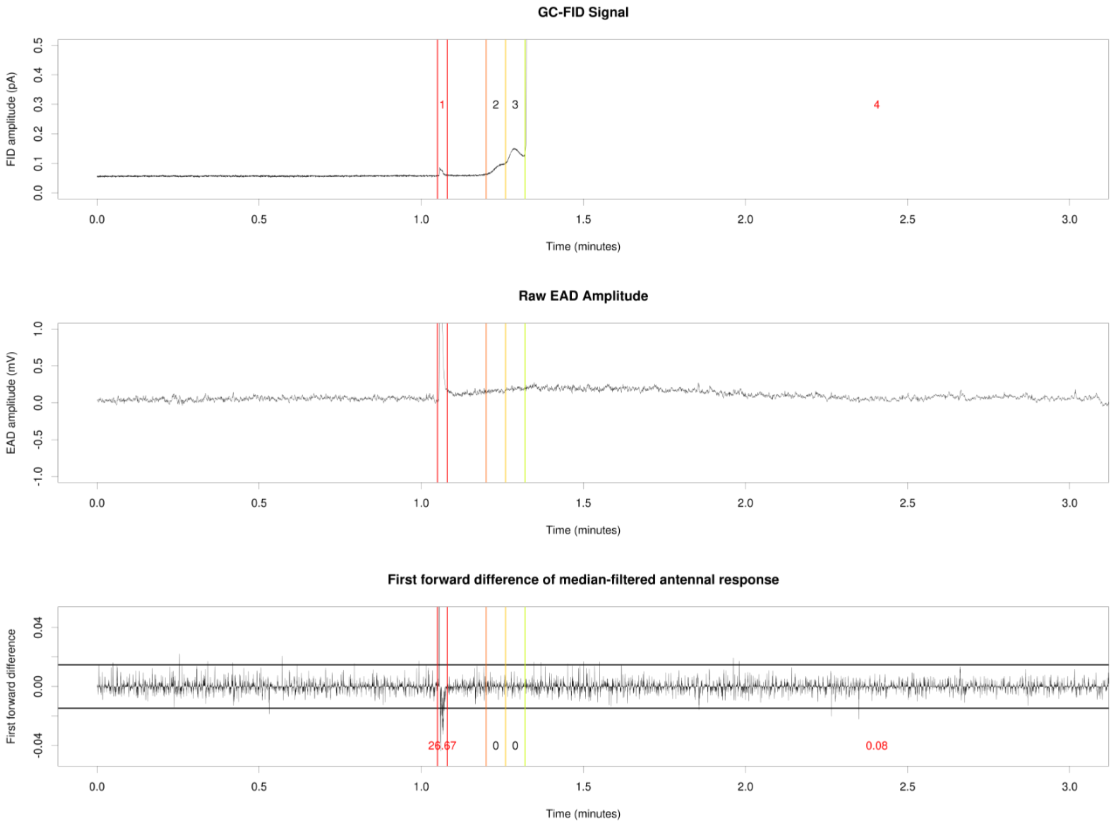


**Figure S31:** Reanalysis of unpublished GC-EAD data from the Pheromone Group, University of Lund (raw data kindly provided by Dr. Glenn Svensson). Data show the response of a male *Plodia interpunctella* (Lepidoptera: Pyralidae) to a synthetic mixture of four female pheromone compounds. (A) GC-FID trace annotated with manually identified GC-FID peak starts and ends (colored vertical lines) and GC-FID peak numbers. (B) Raw EAD amplitude annotated with GC-FID peak starts and ends. (C) First forward difference-processed median-filtered EAD amplitude annotated with GC-FID peak starts and ends. GC-FID peaks showing significant EAD spike frequency are annotated with that frequency in red text in (C) (all annotated peaks within this time window). 0-3 minutes of GC-FID elapsed time.

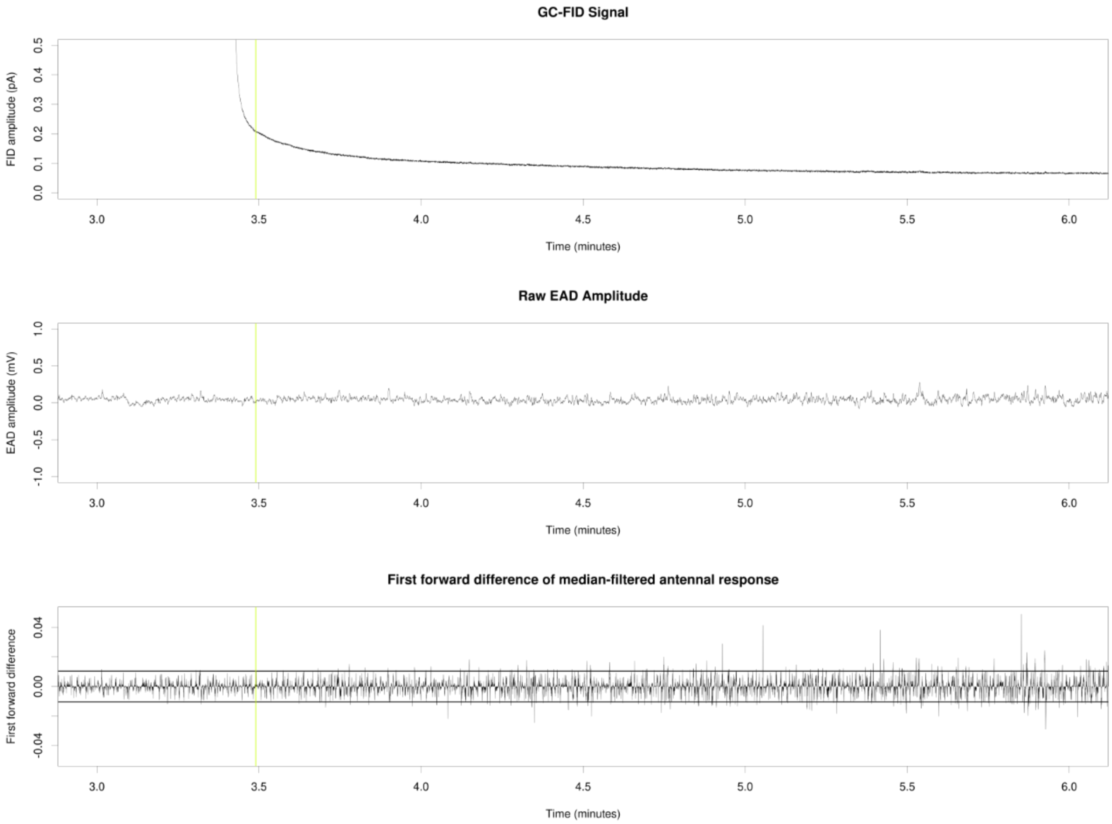


**Figure S32:** Reanalysis of unpublished GC-EAD data from the Pheromone Group, University of Lund (raw data kindly provided by Dr. Glenn Svensson). Data show the response of a male *Plodia interpunctella* (Lepidoptera: Pyralidae) to a synthetic mixture of four female pheromone compounds. (A) GC-FID trace annotated with manually identified GC-FID peak starts and ends (colored vertical lines) and GC-FID peak numbers. (B) Raw EAD amplitude annotated with GC-FID peak starts and ends. (C) First forward difference-processed median-filtered EAD amplitude annotated with GC-FID peak starts and ends. GC-FID peaks showing significant EAD spike frequency are annotated with that frequency in red text in (C) (all annotated peaks within this time window). 3-6 minutes of GC-FID elapsed time.

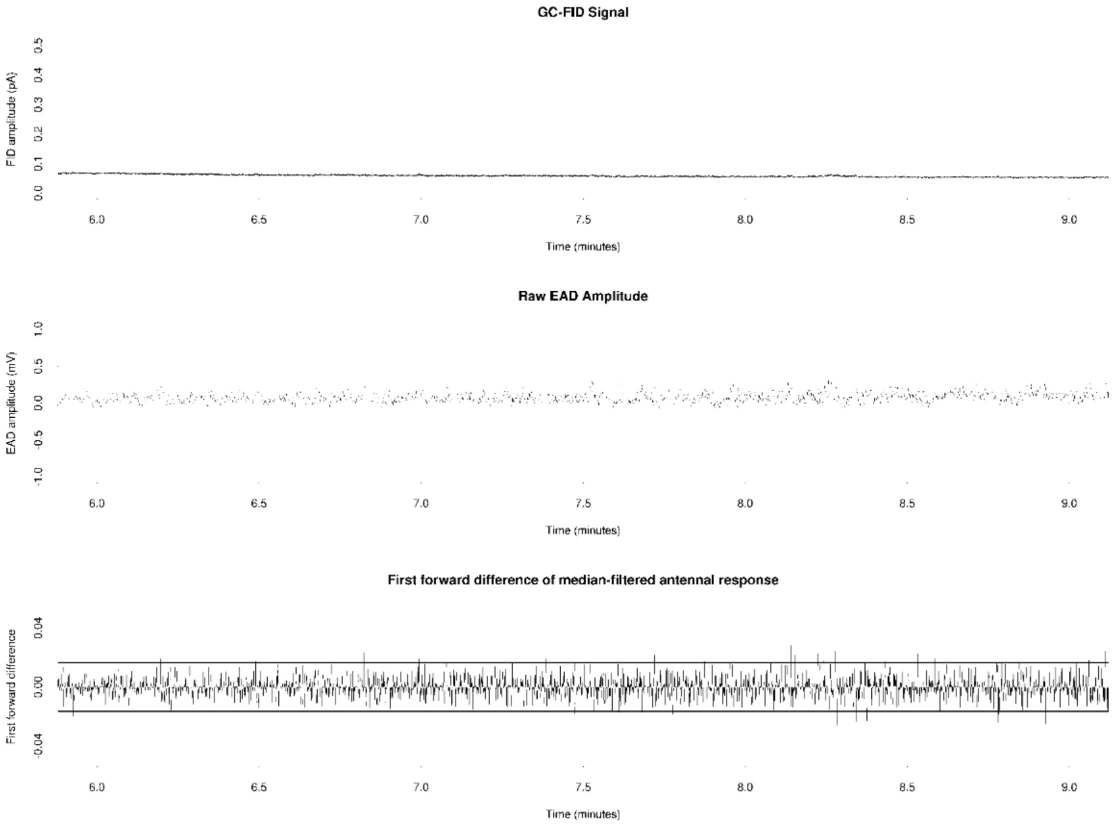


**Figure S33:** Reanalysis of unpublished GC-EAD data from the Pheromone Group, University of Lund (raw data kindly provided by Dr. Glenn Svensson). Data show the response of a male *Plodia interpunctella* (Lepidoptera: Pyralidae) to a synthetic mixture of four female pheromone compounds. (A) GC-FID trace annotated with manually identified GC-FID peak starts and ends (colored vertical lines) and GC-FID peak numbers. (B) Raw EAD amplitude annotated with GC-FID peak starts and ends. (C) First forward difference-processed median-filtered EAD amplitude annotated with GC-FID peak starts and ends. GC-FID peaks showing significant EAD spike frequency are annotated with that frequency in red text in (C) (all annotated peaks within this time window). 6-9 minutes of GC-FID elapsed time.

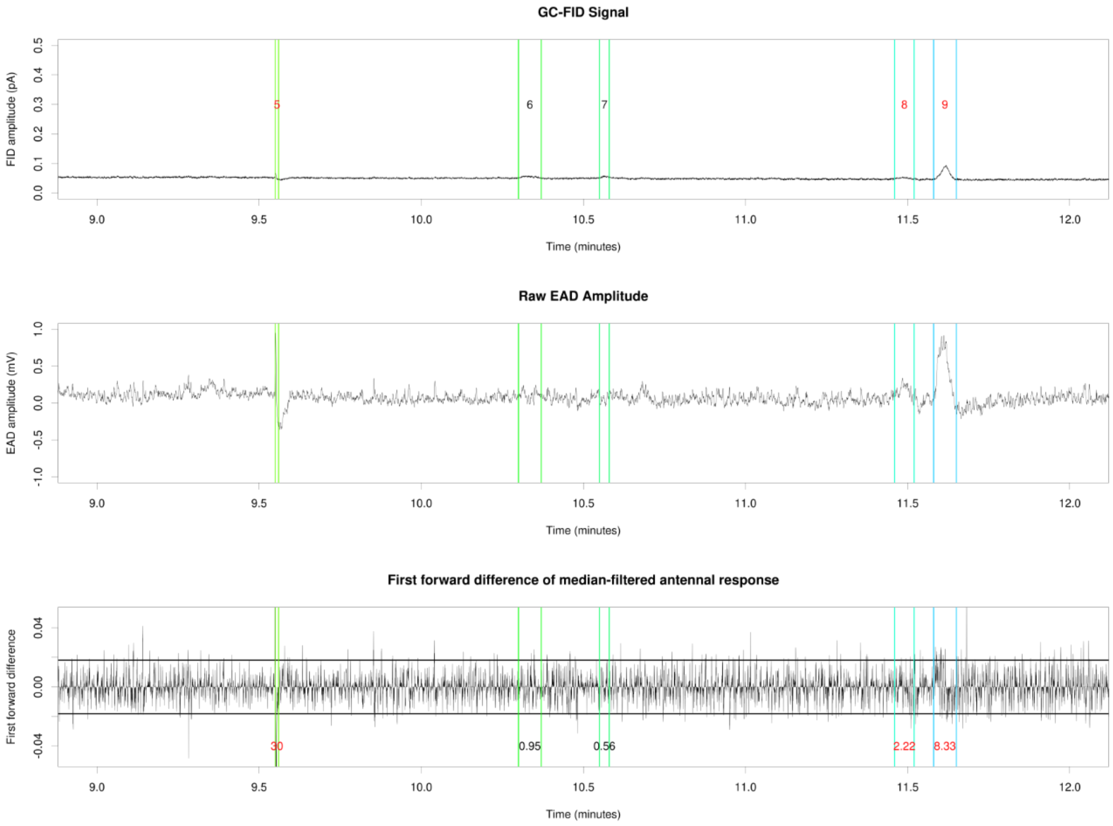


**Figure S34:** Reanalysis of unpublished GC-EAD data from the Pheromone Group, University of Lund (raw data kindly provided by Dr. Glenn Svensson). Data show the response of a male *Plodia interpunctella* (Lepidoptera: Pyralidae) to a synthetic mixture of four female pheromone compounds. (A) GC-FID trace annotated with manually identified GC-FID peak starts and ends (colored vertical lines) and GC-FID peak numbers. (B) Raw EAD amplitude annotated with GC-FID peak starts and ends. (C) First forward difference-processed median-filtered EAD amplitude annotated with GC-FID peak starts and ends. GC-FID peaks showing significant EAD spike frequency are annotated with that frequency in red text in (C) (all annotated peaks within this time window). 9-12 minutes of GC-FID elapsed time.

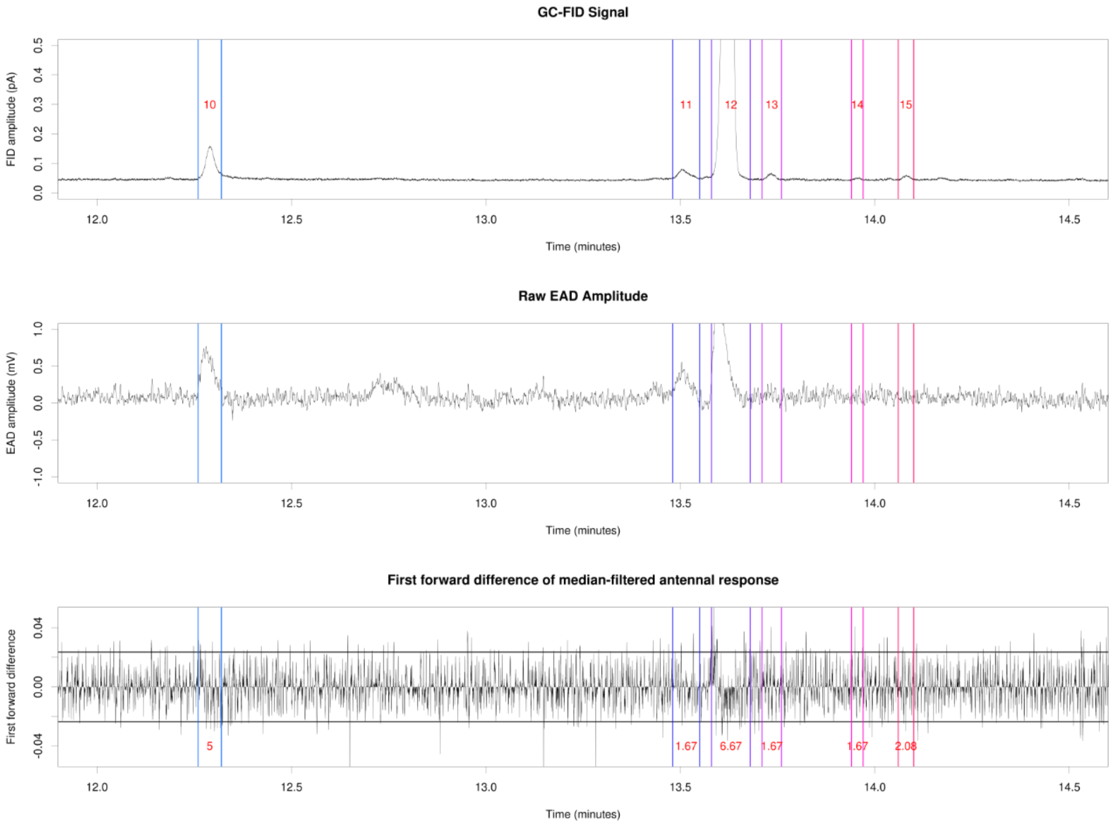


**Figure S35:** Reanalysis of unpublished GC-EAD data from the Pheromone Group, University of Lund (raw data kindly provided by Dr. Glenn Svensson). Data show the response of a male *Plodia interpunctella* (Lepidoptera: Pyralidae) to a synthetic mixture of four female pheromone compounds. (A) GC-FID trace annotated with manually identified GC-FID peak starts and ends (colored vertical lines) and GC-FID peak numbers. (B) Raw EAD amplitude annotated with GC-FID peak starts and ends. (C) First forward difference-processed median-filtered EAD amplitude annotated with GC-FID peak starts and ends. GC-FID peaks showing significant EAD spike frequency are annotated with that frequency in red text in (C) (all annotated peaks within this time window). 12-14.5 minutes of GC-FID elapsed time.

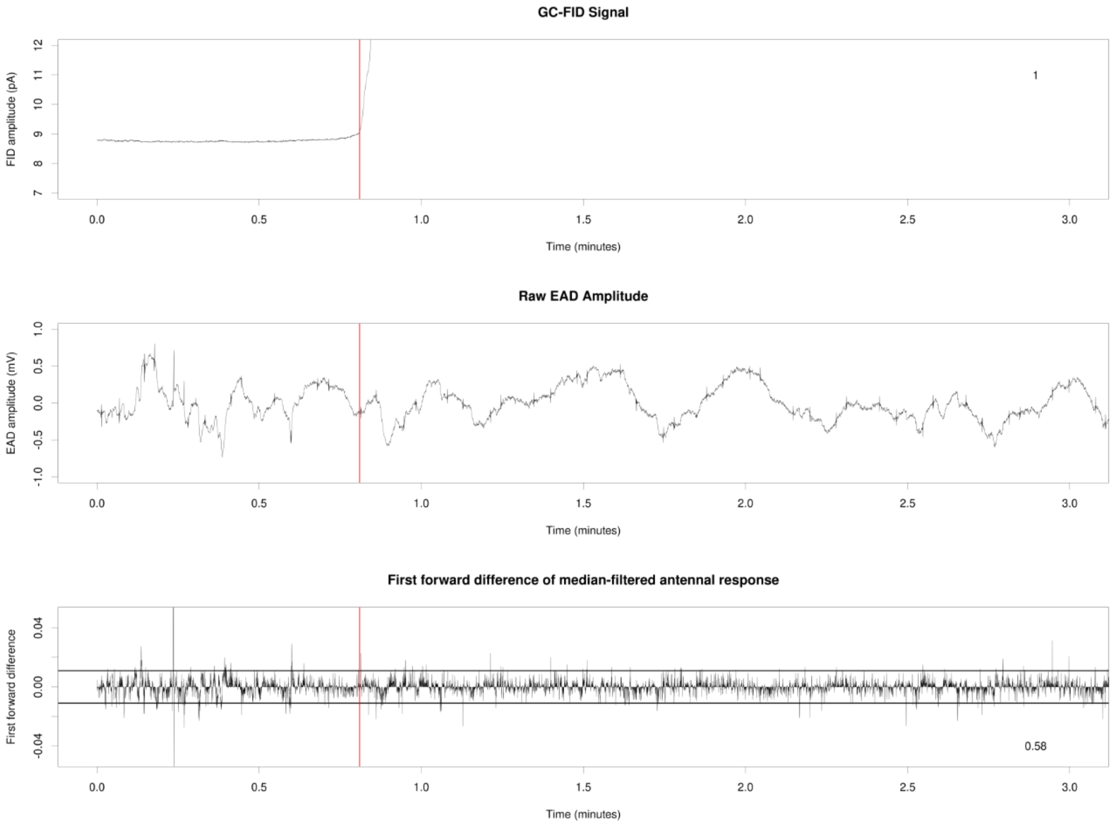


**Figure S36**: Reanalysis of published GC-EAD data from the Pheromone Group, University of Lund (raw data kindly provided by Dr. Glenn Svensson) (Svensson et al., 2019). Data show the response of a male *Acleris comariana* (Lepidoptera: Tortricidae) to an extract of the female pheromone gland. (A) GC-FID trace annotated with manually identified GC-FID peak starts and ends (colored vertical lines) and GC-FID peak numbers. (B) Raw EAD amplitude annotated with GC-FID peak starts and ends; note the regular small spikes every few seconds, presumably due to electrical noise, which were later removed using a median filter. (C) First forward difference-processed median-filtered EAD amplitude annotated with GC-FID peak starts and ends. GC-FID peaks showing significant EAD spike frequency are annotated with that frequency in red text in (C). 0-3 minutes of GC-FID elapsed time.

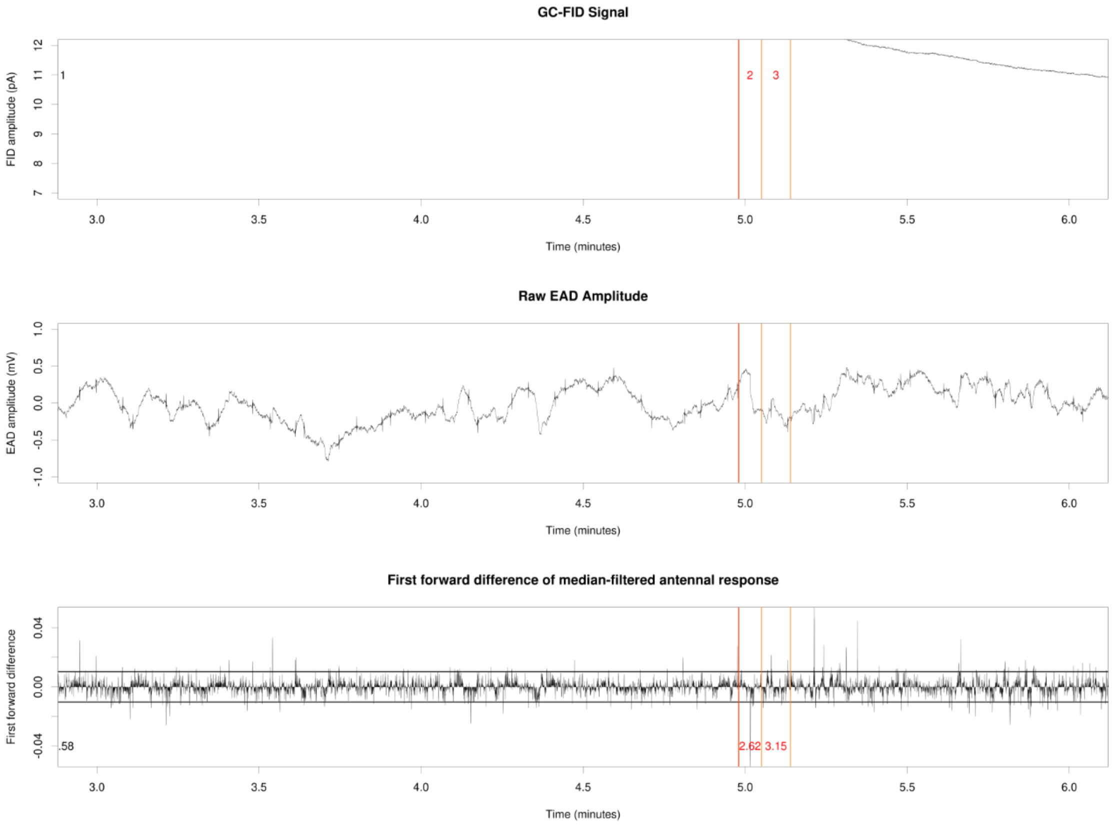


**Figure S37**: Reanalysis of published GC-EAD data from the Pheromone Group, University of Lund (raw data kindly provided by Dr. Glenn Svensson) (Svensson et al., 2019). Data show the response of a male *Acleris comariana* (Lepidoptera: Tortricidae) to an extract of the female pheromone gland. (A) GC-FID trace annotated with manually identified GC-FID peak starts and ends (colored vertical lines) and GC-FID peak numbers. (B) Raw EAD amplitude annotated with GC-FID peak starts and ends; note the regular small spikes every few seconds, presumably due to electrical noise, which were later removed using a median filter. (C) First forward difference-processed median-filtered EAD amplitude annotated with GC-FID peak starts and ends. GC-FID peaks showing significant EAD spike frequency are annotated with that frequency in red text in (C). 3-6 minutes of GC-FID elapsed time.

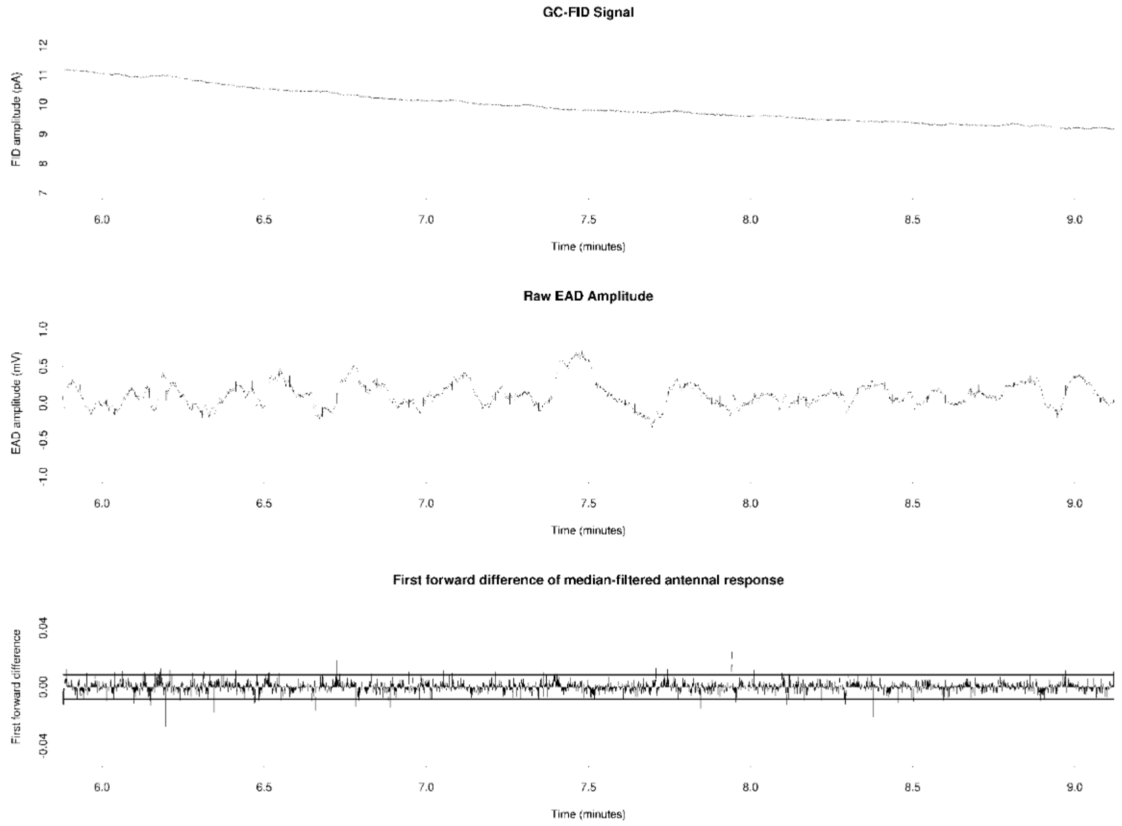


**Figure S38**: Reanalysis of published GC-EAD data from the Pheromone Group, University of Lund (raw data kindly provided by Dr. Glenn Svensson) (Svensson et al., 2019). Data show the response of a male *Acleris comariana* (Lepidoptera: Tortricidae) to an extract of the female pheromone gland. (A) GC-FID trace annotated with manually identified GC-FID peak starts and ends (colored vertical lines) and GC-FID peak numbers. (B) Raw EAD amplitude annotated with GC-FID peak starts and ends; note the regular small spikes every few seconds, presumably due to electrical noise, which were later removed using a median filter. (C) First forward difference-processed median-filtered EAD amplitude annotated with GC-FID peak starts and ends. GC-FID peaks showing significant EAD spike frequency are annotated with that frequency in red text in (C). 6-9 minutes of GC-FID elapsed time.

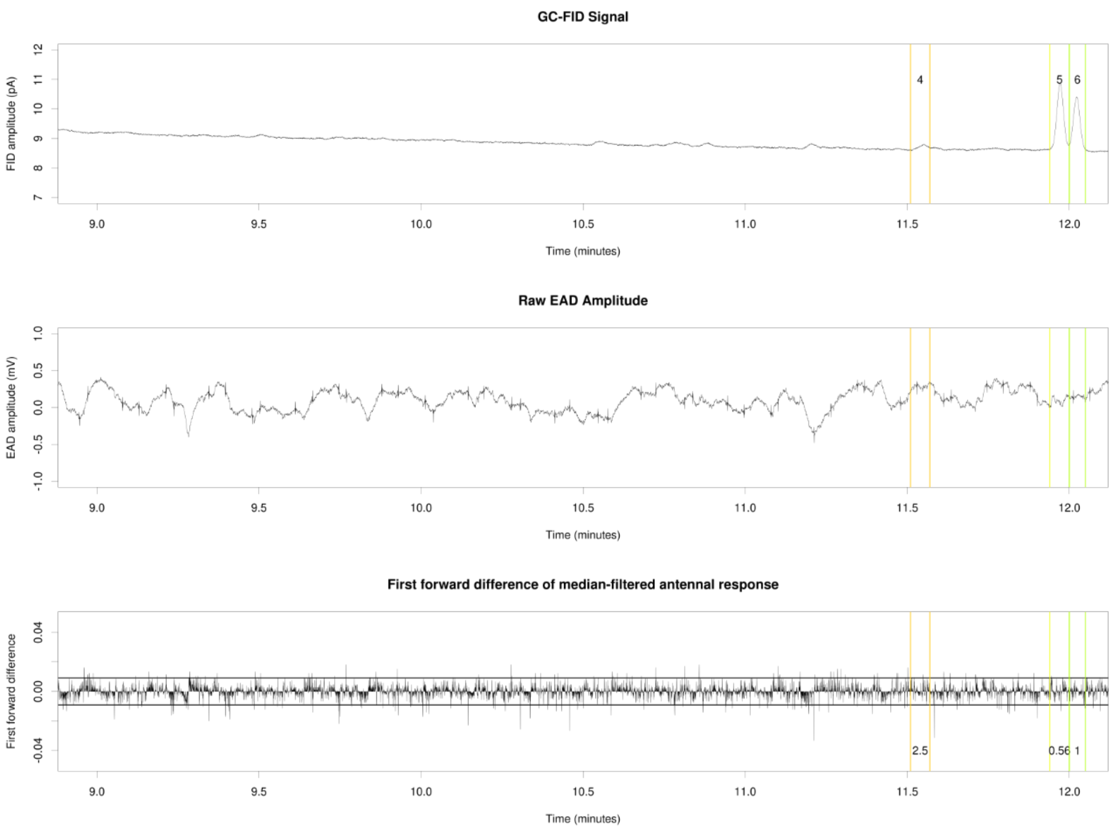


**Figure S39**: Reanalysis of published GC-EAD data from the Pheromone Group, University of Lund (raw data kindly provided by Dr. Glenn Svensson) (Svensson et al., 2019). Data show the response of a male *Acleris comariana* (Lepidoptera: Tortricidae) to an extract of the female pheromone gland. (A) GC-FID trace annotated with manually identified GC-FID peak starts and ends (colored vertical lines) and GC-FID peak numbers. (B) Raw EAD amplitude annotated with GC-FID peak starts and ends; note the regular small spikes every few seconds, presumably due to electrical noise, which were later removed using a median filter. (C) First forward difference-processed median-filtered EAD amplitude annotated with GC-FID peak starts and ends. GC-FID peaks showing significant EAD spike frequency are annotated with that frequency in red text in (C). 9-12 minutes of GC-FID elapsed time.

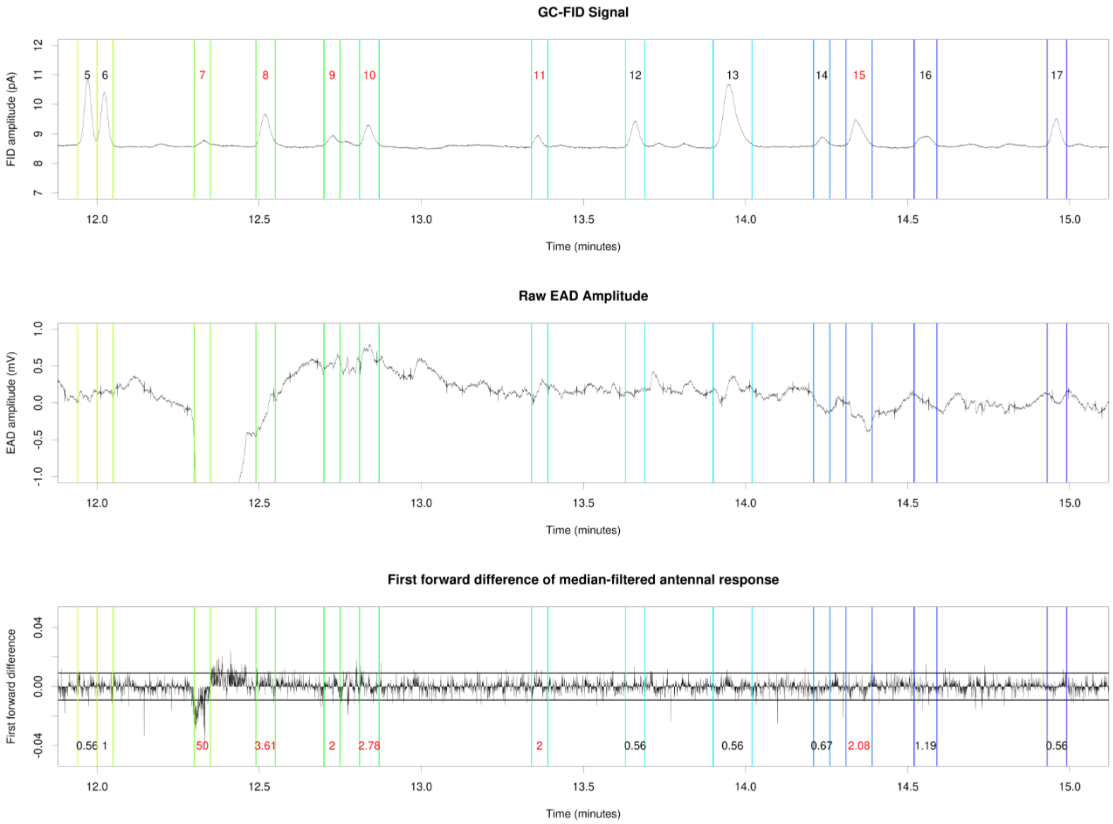


**Figure S40**: Reanalysis of published GC-EAD data from the Pheromone Group, University of Lund (raw data kindly provided by Dr. Glenn Svensson) (Svensson et al., 2019). Data show the response of a male *Acleris comariana* (Lepidoptera: Tortricidae) to an extract of the female pheromone gland. (A) GC-FID trace annotated with manually identified GC-FID peak starts and ends (colored vertical lines) and GC-FID peak numbers. (B) Raw EAD amplitude annotated with GC-FID peak starts and ends; note the regular small spikes every few seconds, presumably due to electrical noise, which were later removed using a median filter. (C) First forward difference-processed median-filtered EAD amplitude annotated with GC-FID peak starts and ends. GC-FID peaks showing significant EAD spike frequency are annotated with that frequency in red text in (C). 12-15 minutes of GC-FID elapsed time.

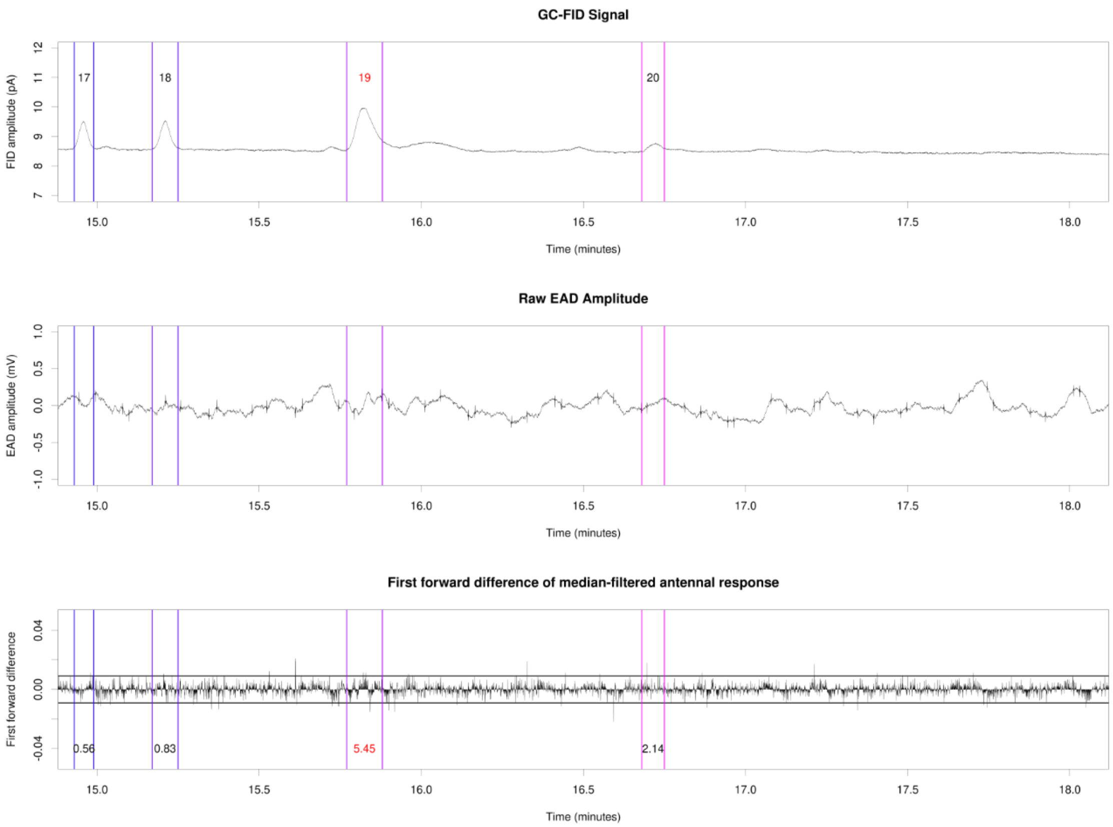


**Figure S41**: Reanalysis of published GC-EAD data from the Pheromone Group, University of Lund (raw data kindly provided by Dr. Glenn Svensson) (Svensson et al., 2019). Data show the response of a male *Acleris comariana* (Lepidoptera: Tortricidae) to an extract of the female pheromone gland. (A) GC-FID trace annotated with manually identified GC-FID peak starts and ends (colored vertical lines) and GC-FID peak numbers. (B) Raw EAD amplitude annotated with GC-FID peak starts and ends; note the regular small spikes every few seconds, presumably due to electrical noise, which were later removed using a median filter. (C) First forward difference-processed median-filtered EAD amplitude annotated with GC-FID peak starts and ends. GC-FID peaks showing significant EAD spike frequency are annotated with that frequency in red text in (C). 15-18 minutes of GC-FID elapsed time.

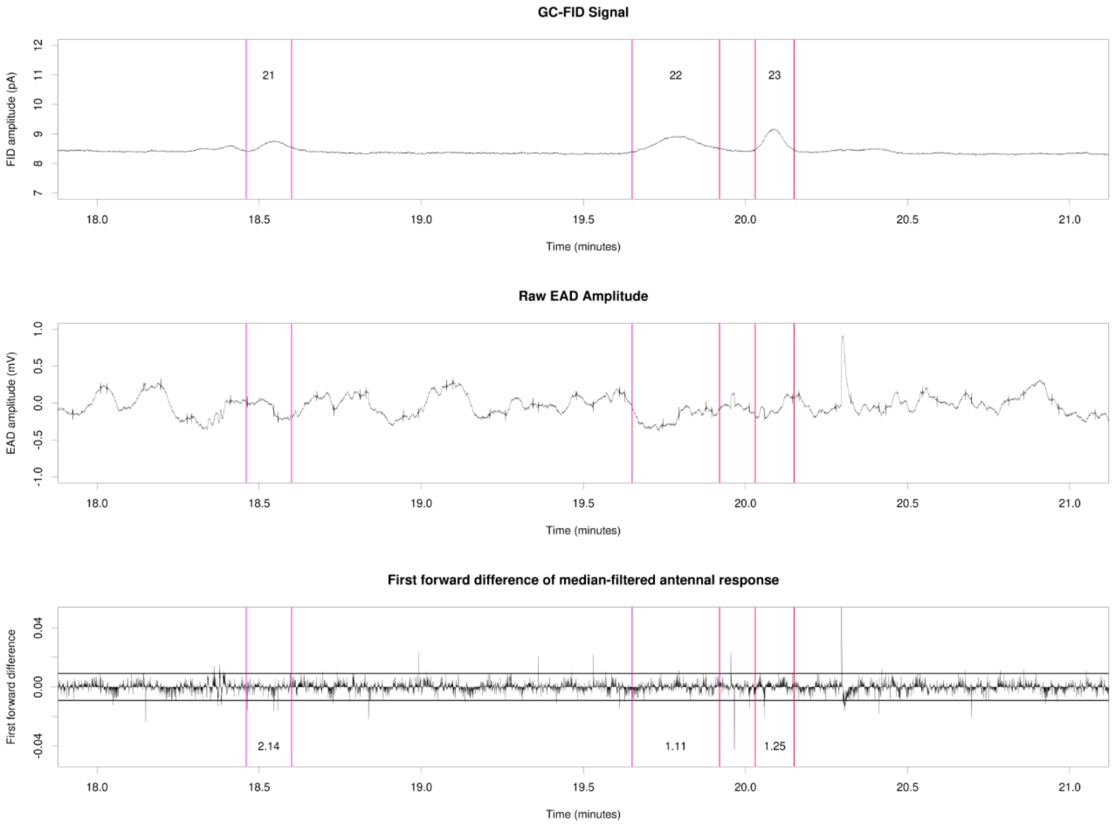


**Figure S42**: Reanalysis of published GC-EAD data from the Pheromone Group, University of Lund (raw data kindly provided by Dr. Glenn Svensson) (Svensson et al., 2019). Data show the response of a male *Acleris comariana* (Lepidoptera: Tortricidae) to an extract of the female pheromone gland. (A) GC-FID trace annotated with manually identified GC-FID peak starts and ends (colored vertical lines) and GC-FID peak numbers. (B) Raw EAD amplitude annotated with GC-FID peak starts and ends; note the regular small spikes every few seconds, presumably due to electrical noise, which were later removed using a median filter. (C) First forward difference-processed median-filtered EAD amplitude annotated with GC-FID peak starts and ends. GC-FID peaks showing significant EAD spike frequency are annotated with that frequency in red text in (C). 18-21 minutes of GC-FID elapsed time.

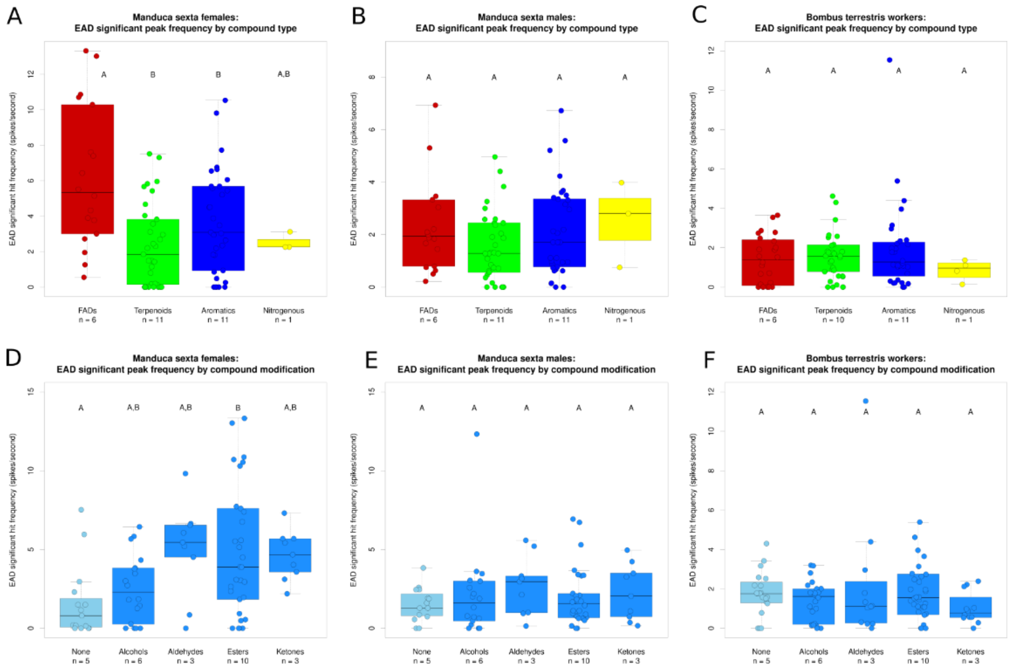


**Figure S43:** First forward difference-processed EAD significant peak hit frequency (spikes/second) across (A,B,C) four chemical skeleton classes (FADs, terpenoids, aromatics, and nitrogenous compounds) and (D,E,F) six compound modification types tested (none, alcohols, aldehydes, esters, and ketones) for (A,D) three *Manduca sexta* females, (B,E) three *Manduca sexta* males, and (C,F) four *Bombus terrestris audax* workers.

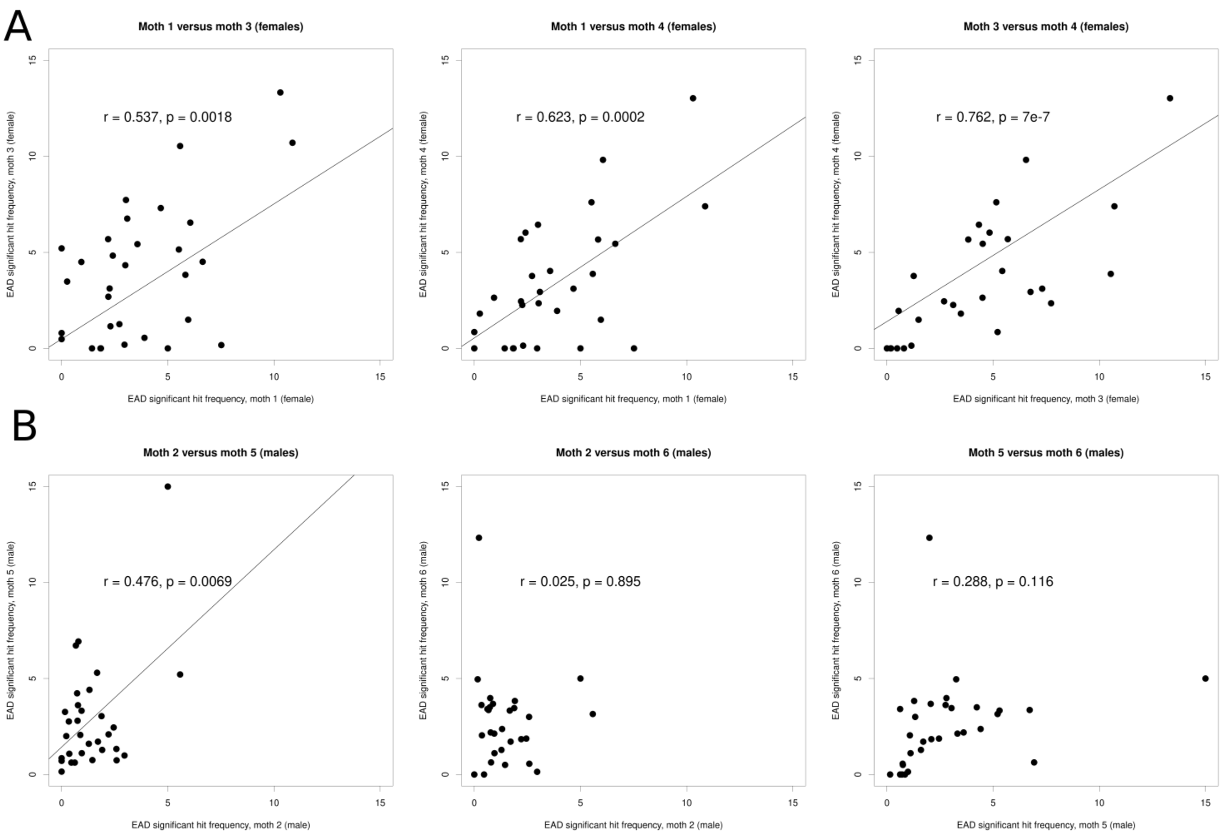


**Figure S44:** Inter-individual EAD response (repeatability) comparisons between *Manduca sexta* hawkmoths of the same sex. (A) All pairwise comparisons between three females. (B) All pairwise comparisons between three males.

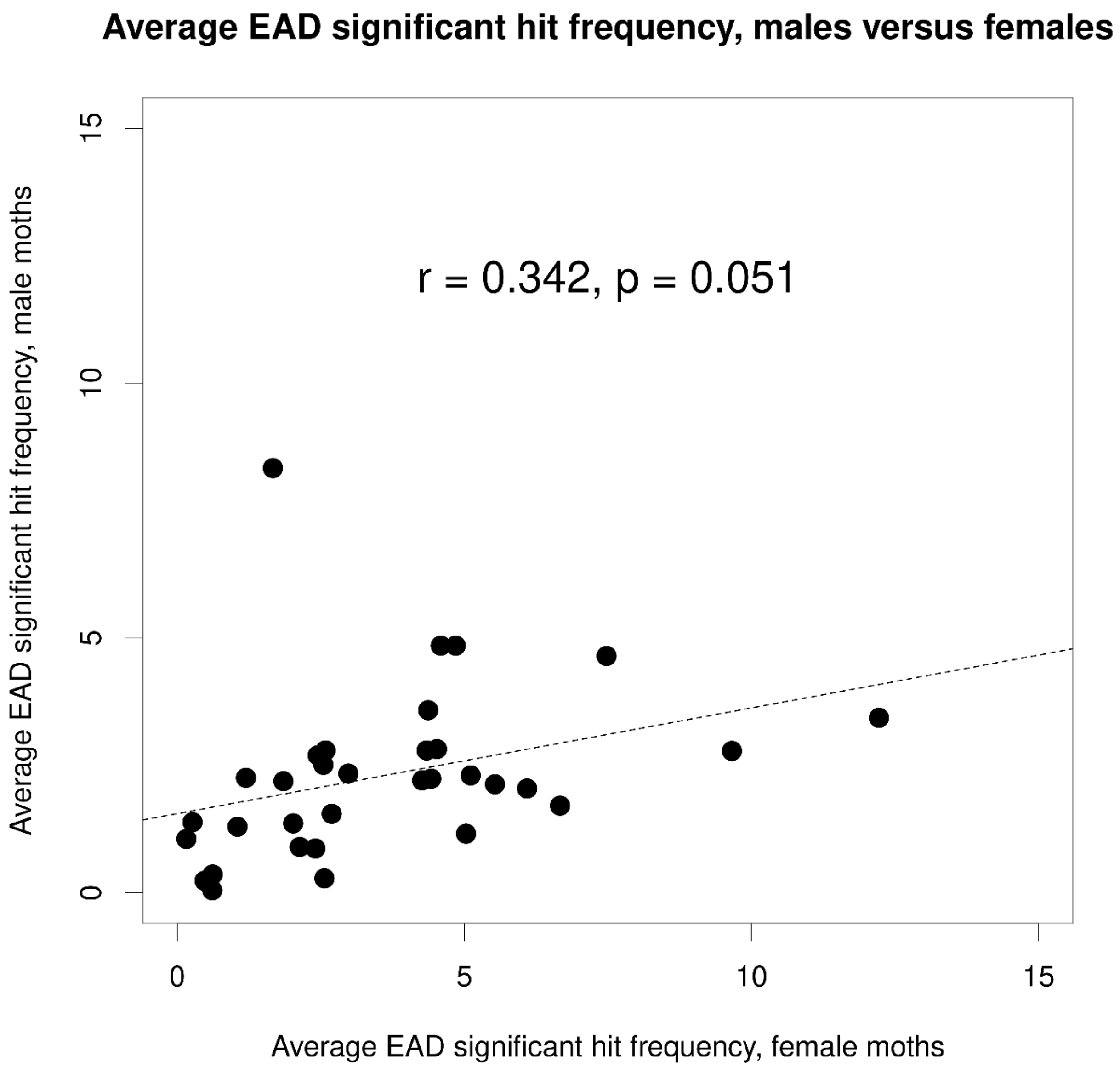


**Figure S45:** Inter-sex EAD response (repeatability) comparisons between the average values of male and female hawkmoths for each GC-FID peak.

**Table S1:** Names and addresses of vendors from whom authentic reference standards were sourced.

| **Vendor** | **Address** |
|---|---|
| Fisher Scientific | Bishop Meadow Rd, Loughborough LE11 5RG United Kingdom |
| Merck (including Millipore) | Suite 21, Building 6, Croxley Green Business Park, Watford, Hertfordshire WD18 8YH United Kingdom |

**Table S2:** Sources of authentic reference standards used in the EAD test mixture and final concentrations used.

| Compound identity | Compound source and catalog number | Compound chemical class | Compound modifications / subcategory | Final concentration in EAD test mixture (ng/µl) |
|---|---|---|---|---|
| Hexane (solvent) | Fisher, H/0406/PB17 | FAD | None | Not applicable |
| Ethyl L-lactate | Fisher, A10900.36 | FAD | Ester | 519 |
| (*Z*)-3-hexen-1-ol | Merck, H12900-10G | FAD | Alcohol | 223 |
| Isoamyl acetate | Millipore, 1.01231.1000 | FAD | Ester | 139 |
| Ethyl valerate | Merck, W246204-SAMPLE-K | FAD | Ester | 213 |
| (1R)-(+)-α-pinene | Merck, W290238-SAMPLE-K | Terpenoid | Monoterpene | 139 |
| Camphene | Merck, W222910-SAMPLE-K | Terpenoid | Monoterpene | 111 |
| Benzaldehyde | Merck, W212709-SAMPLE-K | Aromatic | Aldehyde | 1048 |
| (*Z*)-3-hexenyl acetate | Merck, W247804-SAMPLE-K | FAD | Ester | 195 |
| (R)-(+)-limonene | Merck, 183164-500ML | Terpenoid | Monoterpene | 93 |
| Phenylacetaldehyde | Merck, W287407-SAMPLE-K | Aromatic | Aldehyde | 696 |
| Acetophenone | Merck, A10701-100ML | Aromatic | Ketone | 148 |
| Methyl benzoate | Merck, W268305-SAMPLE | Aromatic | Ester | 167 |
| (1R)-(+)-nopinone | Merck, 327956-1G | Terpenoid | Monoterpenoid ketone | 176 |
| Benzyl acetate | Merck, W213500-SAMPLE-K | Aromatic | Ester | 186 |
| 1-phenylethyl acetate | Merck, 8437850100 | Aromatic | Ester | 111 |
| (1S)-(-)-verbenone | Merck, W506907-SAMPLE-K | Terpenoid | Monoterpenoid ketone | 167 |
| Geraniol | Merck, W250708-SAMPLE-K | Terpenoid | Monoterpenoid alcohol | 297 |
| (*E*)-cinnamaldehyde | Merck, C80687-500G | Aromatic | Aldehyde | 1428 |
| Indole | Merck, W259306-100G-K | Nitrogenous | Aromatic | 2004 |
| 3-phenylpropyl acetate | Merck, W289000-SAMPLE-K | Aromatic | Ester | 176 |
| β-caryophyllene | Merck, W225207-SAMPLE-K | Terpenoid | Sesquiterpene | 102 |

| | | | | |
|---|---|---|---|---|
| α -humulene | Merck, PHL83351-100MG | Terpenoid | Sesquiterpene | 139 |
| Phenethyl 2-methylbutyrate | Merck, W363200-1KG-K | Aromatic | Ester | 120 |
| Methyl isoeugenol | Merck, W247618-SAMPLE-K | Aromatic | Alcohol | 269 |
| Caryophyllene oxide | Merck, W509647-SAMPLE | Terpenoid | Sesquiterpenoid oxide | 427 |
| Ethyl laurate | Merck, W244104-SAMPLE-K | FAD | Ester | 167 |
| Farnesol | Merck, W247804-SAMPLE-K | Terpenoid | Sesquiterpenoid alcohol | 352 |
| Benzyl benzoate | Merck, W213802-SAMPLE-K | Aromatic | Ester | 186 |

**Table S3**: GC-EAD responses by *Plodia interpunctella* to a test mixture of 4 pheromone components. Bolded response values with * were significant above the background response.

| **Retention time (GC-FID, minutes)** | **Peak number; compound identity if known** | ***Plodia* response (significant hit frequency, spikes/second)** |
|---:|---|---:|
| 1.065 | 1 | **26.67 *** |
| 1.230 | 2 | 0 |
| 1.290 | 3 | 0 |
| 2.405 | 4 | 0.08 |
| 9.555 | 5 | **30.00 *** |
| 10.335 | 6 | 0.95 |
| 10.565 | 7 | 0.56 |
| 11.490 | 8 | **2.22 *** |
| 11.615 | 9 | **8.33 *** |
| 12.290 | 10 | **5.00 *** |
| 13.515 | 11 | **1.67 *** |
| 13.630 | 12 | **6.67 *** |
| 13.735 | 13 | **1.67 *** |
| 13.955 | 14 | **1.67 *** |
| 14.080 | 15 | **2.08 *** |

**Table S4:** GC-EAD responses by *Acleris comariana* to a natural pheromone gland extract. Bolded response values with * were significant above the background response. Note the particularly significant response (significant hit frequency 50.00 spikes/second) to the bioactive pheromone component (*E*)-11,13-tetradecadienal (peak 7).

| **Retention time (GC-FID, minutes)** | **Peak number; compound identity if known** | ***Acleris* response (significant hit frequency, spikes/second)** |
|---:|---|---:|
| 2.895 | 1 | 0.58 |
| 5.015 | 2 | **2.62 *** |
| 5.095 | 3 | **3.15 *** |
| 11.540 | 4 | 2.50 |
| 11.970 | 5 | 0.56 |
| 12.025 | 6 | 1.00 |
| 12.325 | 7; (*E*)-11,13-tetradecadienal | **50.00 *** |
| 12.520 | 8 | **3.61 *** |
| 12.725 | 9 | **2.00 *** |
| 12.840 | 10 | **2.78 *** |
| 13.365 | 11 | **2.00 *** |
| 13.660 | 12 | 0.56 |
| 13.960 | 13 | 0.56 |
| 14.235 | 14 | 0.67 |
| 14.350 | 15 | **2.08 *** |
| 14.555 | 16 | 1.19 |
| 14.960 | 17 | 0.56 |
| 15.210 | 18 | 0.83 |
| 15.825 | 19 | **5.45 *** |
| 16.715 | 20 | 2.14 |
| 18.530 | 21 | 2.14 |
| 19.785 | 22 | 1.11 |
| 20.090 | 23 | 1.25 |

**Table S5:** Literature sources demonstrating previous electroantennogram activity or lack thereof in *Manduca sexta* and *Bombus terrestris* to the compounds used in this study. Note the frequent discrepancies in activity (especially for activity in *B. terrestris*), even for compounds with multiple citations. Compounds with at least one active citation in either species are highlighted in bold. Compounds shaded in yellow were EAD-active in more than 2/3 of *M. sexta* males or females and/or *B. terrestris* worker individuals from this study.

| **Compound** | **Electroantennographically active in *Manduca sexta* males or females according to previous studies? (citation(s) in superscript)** | **Electroantennographically active in *Bombus terrestris* workers according to previous studies? (citation(s) in superscript)** |
|---|---|---|
| Ethyl L-lactate | *(no citations found)* | *(no citations found)* |
| **(*Z*)-3-hexen-1-ol** | *(no citations found)* | **Active[1]; inactive[2]** |
| **Isoamyl acetate** | **Active[3]** | **Active[4,5]** |
| Ethyl valerate | *(no citations found)* | *(no citations found)* |
| **(1R)-(+)-α-pinene** | Inactive[3] | **Active[1]; inactive[6]** |
| Camphene | *(no citations found)* | Inactive[6] |
| **Benzaldehyde** | **Active[3,7,8,9]** | **Active[10,11,12]; inactive[2,6]** |
| **(*Z*)-3-hexenyl acetate** | **Active[8,9,13]** | **Active[14]; inactive[2]** |
| **(R)-(+)-limonene** | **Active[9]; inactive[3]** | **Active[4]; inactive[2,6]** |
| **Phenylacetaldehyde** | **Active[13]** | **Active[2]** |
| **Acetophenone** | **Active[3,9]** | **Active[2,11]** |
| **Methyl benzoate** | **Active[3,7,9]** | **Active[10,11]; inactive[2,6]** |
| (1R)-(+)-nopinone | *(no citations found)* | *(no citations found)* |
| **Benzyl acetate** | **Active[3]** | Inactive[2] |
| 1-phenylethyl acetate | *(no citations found)* | Inactive[2] |
| (1S)-(-)-verbenone | *(no citations found)* | *(no citations found)* |
| **Geraniol** | **Active[3,8,9]** | **Active[4,15,16,17]** |
| (*E*)-cinnamaldehyde | *(no citations found)* | *(no citations found)* |
| **Indole** | **Active[13]** | Inactive[2] |
| 3-phenylpropyl acetate | *(no citations found)* | *(no citations found)* |
| **β-caryophyllene** | **Active[8,9]; inactive[3]** | **Active[1,18]; inactive[19]** |
| α –humulene | Inactive[3,8] | *(no citations found)* |
| Phenethyl 2-methylbutyrate | *(no citations found)* | *(no citations found)* |
| Methyl isoeugenol | Inactive[3] | *(no citations found)* |
| **Caryophyllene oxide** | **Active[9]** | *(no citations found)* |
| Ethyl laurate | *(no citations found)* | No electroanntenography data, but is present in *B. terrestris* labial gland[20,21] so likely active as a pheromone |
| **Farnesol** | **Active[3,9]** | **Active[15,16]; note that this is part of the recruitment (foraging) pheromone of *B. terrestris*[22]** |
| **Benzyl benzoate** | **Active[3]; perhaps active[7]** | Inactive[14] |

**Table S3 citations:**

8. Fandino, R. A., A. Haverkamp, S. Bisch-Knaden, J. Zhang, S. Bucks, T.-A. Nguyen Thi, A. Werkenthin, J. Ryback, M. Stengl, M. Knaden, B. S. Hansson, and E. Große-Wilde. 2019. Mutagenesis of *orco* impairs foraging but not oviposition in the hawkmoth *Manduca sexta*. *bioRxiv* 541201.

16. Schmalz, F. D. 2022. Processing of behaviorally relevant stimuli at different levels in the bee brain. PhD thesis, Julius-Maximilians-Universität Würzburg, Würzburg, Germany.
