## Appendix 1 for "Quantitative analysis of gas chromatography-coupled electroantennographic detection (GC-EAD) of plant volatiles by insects"

**Short title: Byers & Jacobs – Quantitative insect GC-EAD for plant interaction analysis**

### APPENDIX 1: PROTOCOL FOR COLLECTION AND ANALYSIS OF GC-EAD DATA

#### Equipment and materials

- ☐ **System for separating volatile samples**
    - A gas chromatography-coupled flame ionization detection system (GC-FID, e.g. Agilent 8890 GC with FID, Agilent, Stockport, UK) modified with a glass Y-splitter (e.g. Agilent Ultrainert PressFit Y-Splitter) to separate column eluent in two parts: one part is sent to the GC-FID, the other part to the insect antenna; these are passed through deactivated column (e.g. Agilent CP803210) of equal lengths
    - Appropriate gases for the GC-FID, e.g. hydrogen as the carrier (from either compressed gas bottles or from a hydrogen generator), nitrogen as the make-up, and air for the FID
    - Heated transfer tube installed into the GC-FID wall (e.g. Syntech EC-03, Ockenfels Syntech, Buchenbach, Germany)
    - GC-FID analysis software allowing comma-separated values (CSV) file export, e.g. Agilent OpenLab CDS v2.5
    - A gas chromatography-coupled mass spectrometer detection system (GC-MS), unmodified (e.g. Agilent 7890B GC and 5977/A/B MS, for determining volatile compound identify for unknown samples, e.g. plant headspace)
    - Equivalent column types and lengths for the GC-FID and GC-MS (e.g. Phenomenex Zebron ZB-5HT column, Phenomenex UK Ltd, Macclesfield, UK)
- ☐ **System for collecting electroantennography data**
    - Antenna electrode fork OR pulled glass capillaries and capillary holders with silver wire (e.g. Syntech PRG-2)
    - Electrode gel (with fork, e.g. Signa Gel, Parker Laboratories, Fairfield, New Jersey, USA) OR 0.1N potassium chloride (with capillaries)
    - Electrode mount on a micromanipulator (e.g. Syntech MP-15 with EAG Combi Probe), with a second electrode mount and micromanipulator and dissecting microscope if using glass capillary electrodes
    - Electroantennogram signal amplifier and acquisition system (e.g. Syntech IDAC-2)
    - Constant and pulsed air delivery system (e.g. Syntech CS-55)
    - Silicone tubing and barb fittings to fit the air delivery system and bent tube
    - Glass or metal bent tube to connect air delivery to the output of the GC-FID and the insect
    - Electroantennogram signal acquisition and analysis software (e.g. Syntech GcEAD 2014 v1.2.5)

- ☐ **Insect preparation supplies**
  - 15-50ml conical tubes and/or butterfly envelopes
  - Filter paper (round or square)
  - Red light source (a headlamp is ideal) and dark room for working with stinging insects such as bees
  - Dissection supplies: microdissection scissors, fine scalpel, broad forceps, and pin on a stick
  - Dissection microscope with appropriate lighting

**Sample and insect choice**

- ☐ Volatile samples to separate with the GC to test with insect antennae , here a mixture of purchased chemical standards – this could be mixtures of purchased or synthesized chemical standards, headspace collected from living plant tissue, direct solvent extraction of plant tissue, etc.
- ☐ A live insect of the species and sex or caste of interest, here female and male *Manduca sexta* hawkmoths and *Bombus terrestris audax* bumble bee workers – these could be captive-reared or wild-collected insects (note that prior foraging experience is unlikely to affect results, unlike with behavioural studies).
- ☐ An alkane ladder to facilitate comparison of Kovats retention indices between GC-FID and GC-MS for compound identification.
- ☐ A positive control compound or mixture that the insect is known to respond to – consider searching the literature to find this out. A concentration of 10µg/µl is sufficient.

**Experimental data collection**

1. Ensure the GC-FID is set up with an appropriate method and that detectors, inlet, and oven are at appropriate temperatures and the status is ready. Also turn on the heated transfer tube and set to 250°C or as appropriate.
2. Prepare an odor cartridge as follows: place a small piece of filter paper in a glass Pasteur pipette, ensuring that it is reachable with a regular micropipette tip. Plug the open large end with a bit of cotton wool.
3. Set up the air delivery system with the silicone tubing and barbs (this only needs doing at the first run) so that the pulsed air is flowing through a silicone tube with an open end and the constant air is flowing through the bent tube.
4. Set up the air delivery system for a low to moderate constant flow (e.g. 8 on the Syntech CS-55) and a low to moderate pulsed flow, with the same value as the constant flow. Set up the pulsed flow to repeat three times in fairly short succession each time the button or foot pedal is pressed.
5. Gently push the GC-FID column end going through the heated transfer tube 1-2 millimetres into the bent tube.
6. Collect the insect from the colony or via netting in the wild and place into a 15 or 50ml conical tube or butterfly envelope. If it is a stinging insect such as a bee or ant, consider placing in the

refrigerator at 4°C or on ice for 30 minutes and work with the insect under red light in a dark room to avoid flight.

7. Holding the insect by hand (for larger insects) or with forceps (for stinging insects) over the filter paper, cut off a single antenna using the microdissection scissors such that it lands on the filter paper. For insects with elbowed antennae (e.g. bees), try to include the initial antennal segments before the bend. Place the insect back in the tube or envelope, and back in the fridge if stinging.
8. Transfer the filter paper to the dissection microscope stage.
9. Under good lighting, remove the last half segment of the antenna (or more for some larger antennae) using the microscalpel. Try to get a slight diagonal angle on the cut if using the glass electrodes.
10. If using an antennal fork and gel, load the fork with mustard seed-sized dollops of gel on each of the two ends. If using glass capillary electrodes and electrode holders, load the tubes with potassium chloride solution by dipping the narrower ends into the solution and place them into the electrode holders.
11. Pick up the antenna by gently touching it with the electrode fork and gel assembly or by gently allowing one end of it to be sucked into one of the glass electrodes.
12. If using gel and the antennal fork, work the gel with the pin on a stick to ensure both ends of the antenna are fully enclosed within the gel. Make sure the middle of the antenna between the two ends is not covered with gel and that the gel is not touching between the two sides of the fork.
13. Place the fork or electrode holder into the electrode mount.
14. If using the electrode holders, use a second dissecting microscope and the pin on a stick, along with the second electrode mount micromanipulator with the electrode holder and capillary mounted, to coax the other end of the antenna into the second capillary.
15. Move the antennal assembly to within ca. 1cm of the bent tube outlet (longer antennae should be moved slightly further away to ensure the air stream reaches the entire antennal body).
16. With the antenna seated, open the electroantennogram software and preview the signal. Ensure that the antenna is responsive by gently exhaling on it and seeing if there is a change in the signal.
17. Allow the antenna to rest for 5 minutes while preparing the odor cartridge and GC-FID injection material.
18. While the antenna is resting, load 10 µl of positive control solution onto the filter paper in the odor cartridge using a micropipette and place the open end of the odor cartridge into the silicone tubing coming from the pulsed barb of the air delivery system. The narrow end of the odor cartridge should be placed in the hole in the side of the bent tube that is not taken up by the GC-FID outlet.
19. Once 5 minutes has passed, begin recording with the electroantennogram software.
20. Stimulate the antenna by activating the pulsed air flow through the odor cartridge and make sure the antenna responds. It isn't critical if the response is a peak or a trough, either is fine.
21. If the antenna isn't responding, ensure it is still connected via the gel or capillaries. If this is the case and it is still unresponsive, discard and use the other antenna from the same insect. Never

use both antennae from the same insect as two good samples, as this introduces pseudoreplication into the analysis.

22. Prepare the GC-FID for injection, then inject 3µl of the test sample into the GC-FID. Write down the injection time according to the electroantennogram software.
23. At the end of the GC-FID run, or after all peaks have eluted, stimulate the antenna with the positive control odor cartridge again. If it is still responsive, another GC-FID run can be conducted (antennae typically stay responsive for 1-2 hours, depending on the insect and the quality of the preparation). If it is no longer responsive, the sample must be discarded and the GC-FID run must be considered a failure as it is not clear when the antenna stopped responding.
24. Stop the recording and export it from the electroantennogram software as a CSV file.
25. This protocol can be conducted with multiple insects in a row; attempt to capture at least three replicate insects per species and sex/caste, with more being ideal.

**Statistical analysis**

1. Determine the identities (names) of the GC-FID peaks by comparison to the GC-MS data.
2. Transfer the two CSV files (GC-FID output and electroantennogram software output) to the analysis computer.
3. Using the provided Perl script, identify the peak onsets and offsets in the GC-FID output, setting the second argument (the user-defined threshold for peak onsets and offsets) to a value that captures legitimate peaks but excludes contaminants where possible. If this is unsuitable, use the GC-FID data analysis software to automatically identify peak onsets and offsets and insert into the R script (see step 4).
4. Edit the provided R script to include the filenames, GC-FID onsets and offsets, GC peak identities/names, injection time, run length (the default is 20 minutes), how many windows you would like to calculate the threshold across (the default is 5 windows in 20 minutes), which peaks are within which window, what y-axis limits should be used for the plots, and what output filenames should be used for the plots and data output.
5. Execute the R script in your preferred R workflow software.
6. The output file (e.g. "insect-10cmin-hits-date.csv") will contain the GC-FID peak retention time, the name of the GC-FID peak, the significant hit frequency of the electroantennogram response, and whether the hit was significant (via a color code with black as non-significant and red as significant).
7. Report significant peaks (those peaks with a significant hit frequency larger than the background hit frequency) with their significant hit frequencies.

**Perl script and R script**

Please see https://github.com/plantpollinator/GCEADAnalysis/ for the two scripts cited above.
